# Long-term hunter-gatherer continuity in the Rhine-Meuse region was disrupted by local formation of expansive Bell Beaker groups

**DOI:** 10.1101/2025.03.24.644985

**Authors:** Iñigo Olalde, Eveline Altena, Quentin Bourgeois, Harry Fokkens, Luc Amkreutz, Marie-France Deguilloux, Alessandro Fichera, Damien Flas, Francesca Gandini, Jan F. Kegler, Lisette M. Kootker, Kirsten Leijnse, Leendert Louwe Kooijmans, Roel Lauwerier, Rebecca Miller, Helle Molthof, Pierre Noiret, Daan C. M. Raemaekers, Maïté Rivollat, Liesbeth Smits, John R. Stewart, Theo ten Anscher, Michel Toussaint, Kim Callan, Olivia Cheronet, Trudi Frost, Lora Iliev, Matthew Mah, Adam Micco, Jonas Oppenheimer, Iris Patterson, Lijun Qiu, Gregory Soos, J. Noah Workman, Ceiridwen J. Edwards, Iosif Lazaridis, Swapan Mallick, Nick Patterson, Nadin Rohland, Martin B. Richards, Ron Pinhasi, Wolfgang Haak, Maria Pala, David Reich

## Abstract

The first phase of the ancient DNA revolution painted a broad-brush picture of European Holocene prehistory, whereby 6500-4000 BCE, farmers descending from western Anatolians mixed with local hunter-gatherers resulting in 70-100% ancestry turnover, then 3000-2500 BCE people associated with the Corded Ware complex spread steppe ancestry into north-central Europe. We document an exception to this pattern in the wider Rhine-Meuse area in communities in the wetlands, riverine areas, and coastal areas of the western and central Netherlands, Belgium and western Germany, where we assembled genome-wide data for 109 people 8500-1700 BCE. Here, a distinctive population with high hunter-gatherer ancestry (∼50%) persisted up to three thousand years later than in continental European regions, reflecting limited incorporation of females of Early European Farmer ancestry into local communities. In the western Netherlands, the arrival of the Corded Ware complex was also exceptional: lowland individuals from settlements adopting Corded Ware pottery had hardly any steppe ancestry, despite a characteristic early Corded Ware Y-chromosome. The limited influx may reflect the unique ecology of the region’s river-dominated landscapes, which were not amenable to wholesale adoption of the early Neolithic type of farming introduced by Linearbandkeramik, making it possible for previously established groups to thrive, and creating a persistent but permeable boundary that allowed transfer of ideas and low-level gene flow. This changed with the formation-through-mixture of Bell Beaker using populations ∼2500 BCE by fusion of local Rhine-Meuse people (9-17%) and Corded Ware associated migrants of both sexes. Their expansion from the Rhine-Meuse region then had a disruptive impact across a much wider part of northwest Europe, including Britain where its arrival was the main source of a 90-100% replacement of local Neolithic peoples.

---

Whole genome ancient DNA (aDNA) analysis has illuminated long-standing debates about cultural and demographic transformations in Holocene Europe. Two major prehistoric events have been characterized: the spread of genetic ancestry originating from western Anatolian farmers into Europe associated with the introduction of farming in the Early Neolithic^1–3^, and the spread of ancestry characteristic of Yamnaya steppe pastoralists during the 3rd millennium BCE^4–7^, mediated by the dispersal of the Corded Ware (CW) and Bell Beaker (BB) complexes. However, the demographic processes on a regional level are still imperfectly understood, and, have been shown to follow variable patterns. For example, while the spread of Anatolian ancestry in central Europe was primarily propelled by the expansion of Linearbandkeramik (LBK) farmers^2,4,8–10^, in the Baltic region and Scandinavia, there was late adoption of the farming lifestyle and even, in some cases, a return to hunting, gathering, and fishing^11^. This shift was in part paralleled by a resurgence of local hunter-gatherer ancestry^12–14^.

This paper focuses on the unique trajectory of communities from water-rich environments in the wider Rhine-Meuse area in western and central Netherlands, Belgium, and northern and northwestern Germany. Around 5500 BCE, the southern part of this region witnessed the arrival of LBK-associated farmers, who settled across the fertile loess soils in the south of the Netherlands and parts of Belgium, Germany and France. Within these communities, there is evidence of contact with hunter-gatherer groups, as documented by Limburg and La Hoguette pottery^15^, although the origin and significance^16^ of these contacts is debated^17^. Once established, these LBK communities developed into regional variants such as the Blicquy, Rössen, Villeneuve-Saint-Germain, Bischheim, and the later southern Michelsberg groups.

North of the loess, large rivers such as the Scheldt, Meuse, and Rhine created a dynamic landscape that included fertile soils favoured by farmers, alongside coastlines, beach barriers, river delta wetlands and forested river dunes that continued to support hunting, gathering and fishing practices after 4200 BCE^17–21^. This contrasts with other areas of Europe (with the exception of northern Scandinavia, the Baltic region and the eastern European taiga), where farming practices quickly became dominant^22^. In the Rhine-Meuse area, the wetland communities of the Swifterbant (5^th^ millennium BCE) and Hazendonk cultures (4000-3500 BCE) settled on elevated areas (river and coastal dunes, crevasse splays, and river levees) in a region dominated by water courses and peat bogs. They relied mostly on hunting, gathering, and fishing, but also practiced farming. Around 3500 BCE, the Vlaardingen culture succeeded the Swifterbant/Hazendonk tradition, while remaining settled in approximately the same region^23^. Simultaneously, farmers associated with the Funnelbeaker culture (or TRB for *Trechterbekercultuur* in Dutch) settled on the Friesian-Drenthian plateau in the northeast and its surrounding sandy uplands, in regions where no evidence of earlier habitation, neither burials nor settlements, has been found. The Swifterbant, Hazendonk and Vlaardingen cultures were all located near water streams, while TRB farmers settled mostly on forested sandy plateaus and their fringes, as did the Michelsberg communities to the south.

A mixed subsistence strategy of hunting, gathering, and farming persisted in the western/central Netherlands until the 3rd millennium BCE, when a more intense farming-based economy emerged in association with the Late Vlaardingen complex and the introduction of the ard plough around 3000 BCE^24^. The spread of CW influence to the wider Rhine-Meuse area was more complex than in many areas of central and eastern Europe. In the uplands, where skeletal material tends to be poorly preserved and no ancient DNA data are available, the complete CW package emerged; marked by the construction of CW burial mounds, the general absence of settlements, and sparse pottery finds^25,26^. By contrast, in wetland areas along the coast, the Rhine-Meuse delta^27^, and other low-lying regions^28^, CW-associated pottery was incorporated into Vlaardingen settlement contexts, but the characteristic CW-style burials were not^27,29,30^. South of the Rhine-Meuse rivers, Seine-Oise-Marne (SOM) groups, mostly known through burials and megaliths^31^, had little to no CW-associated material^32,33^.

The arrival of the BB complex around 2500 BCE marked another major cultural transition, as settlements spread across the wetlands and coastal areas, replacing Vlaardingen/CW settlements, though generally not using the same sites^28^. The BB economy was similar to the previous CW one and consisted of predominantly farming mixed with low-intensity hunting and gathering. In the sandy uplands, there was a continuation of the barrow ritual, but with distinct BB characteristics and material culture replacing the CW repertoire^34,35^. BB groups were also well attested south of the Rhine, as evident in BB burial mounds on the sandy soils of the southern Netherlands and Belgium^32,36,37^. BB settlement sites remain just as elusive in this area as CW settlements. However, the presence of ploughland dated to the Late Neolithic suggests that the lack of settlement evidence is not the result of nomadism but rather of settlements in lower lying places where there is little chance for detection by archaeologists^28^.

Archaeogenetic data have the potential to deepen our understanding of the nature of these dynamic changes in the Rhine-Meuse region. We generated genome-wide data using in-solution enrichment for more than a million single nucleotide polymorphisms (SNPs) from 41 individuals dated between 8500-1700 BCE, sampling cultural contexts that fill gaps in the ancient DNA record of this region (Figure 1; Supplementary Table 1). Together with 68 published individuals^6,10,38–41^, the time-transect includes 109 individuals. We also report twelve new direct radiocarbon dates on newly analyzed individuals (Supplementary Table 8).

**Figure 1.**
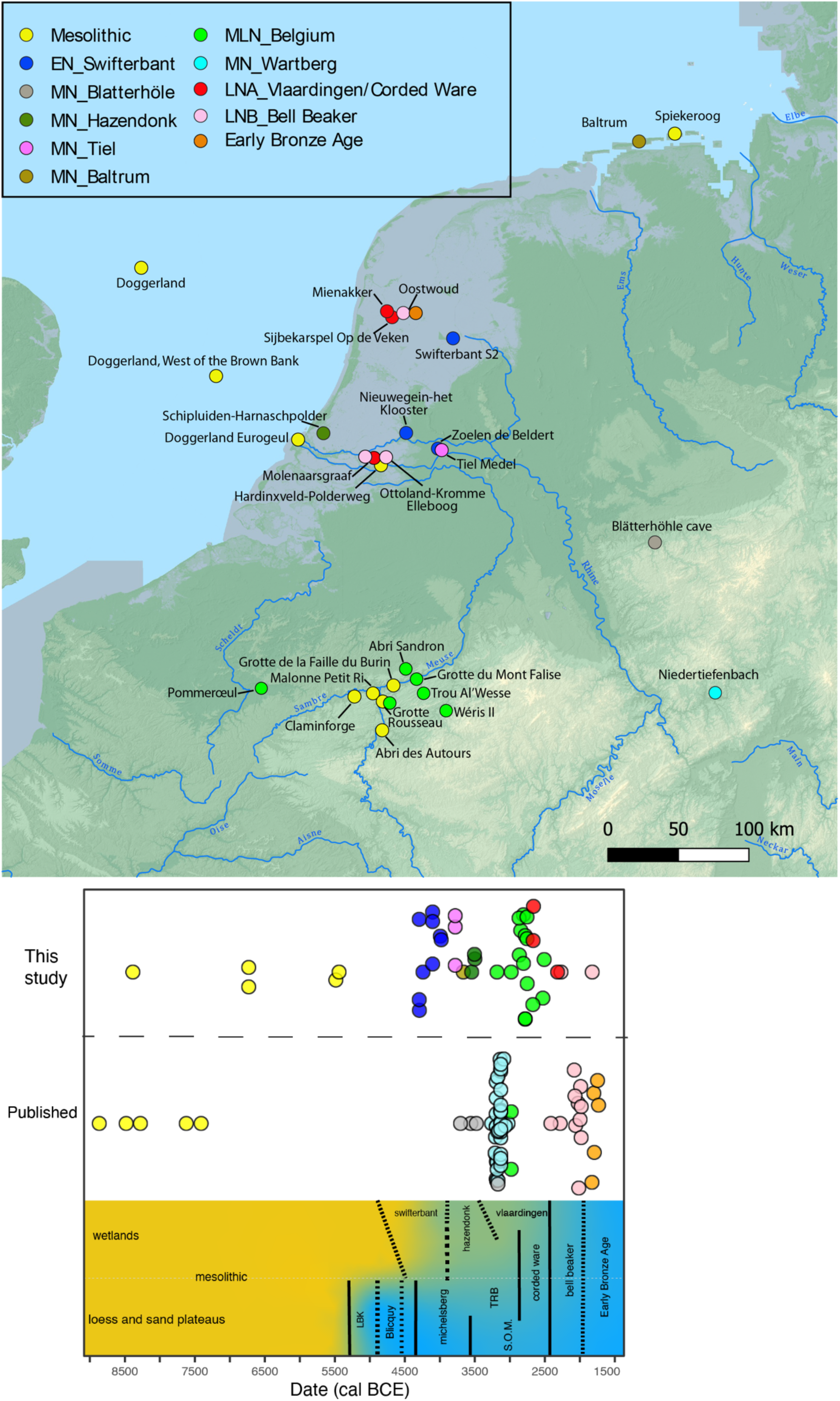
**a)** Map showing archaeological sites with genome-wide data in the Rhine-Meuse area and adjacent regions. **b)** Chronological placement of the individuals from the Rhine-Meuse region included in this study. In the bottom panel the local chronology of archaeological cultures. Black lines indicate the degree of changes between the time periods, with dashed lines representing a gradual and lesser change compared to a solid line indicating a more abrupt change in material culture. The colour gradient indicates the general reliance on hunting and gathering (yellow) to farming (blue).

## Late persistence of Western hunter-gatherer ancestry in the Rhine-Meuse area

Based on Principal Component Analysis (PCA) (Figure 2), Neolithic individuals from the Rhine-Meuse area fall along the central/western European Neolithic cline, but much closer to Mesolithic western hunter-gatherers (WHG) than most European Neolithic farmers. This suggests elevated WHG-related ancestry, which we confirmed through modelling using *qpAdm* (Supplementary Table 3). We find that the earliest Neolithic individuals (4400-3800 BCE), associated with the Swifterbant culture, are genetically highly heterogeneous, with a mother and her daughter (I12093-I12094; Nieuwegein) entirely descending from hunter-gatherer populations, three individuals (I12091-I17968 from Nieuwegein and I33739 from Zoelen de Beldert) with 60-70% of such ancestry (Figure 3; Supplementary Table 3) and four individuals (SWA001-SWA002-SWA004 from Swifterbant S2 and I33738 from Zoelen de Beldert) with 35-45%. These results differ from the overall patterns of hunter-gatherer and farmer admixture elsewhere in central and western Europe, where the arrival of a farming economy was generally associated with <30% local WHG ancestry. However, the results perhaps make more sense in light of the equally limited economic transformation, which combined farming with continued core reliance on the rich wild resources from the Rhine-Meuse wetlands and river valleys. Genetic mixing of local groups with high WHG ancestry continued for the next ∼1500 years, with stable proportions of ∼40-50 % WHG and 50-60% early European farmer (EEF) ancestry. Rare exceptions include one Middle Neolithic individual from the island of Baltrum (BLR001) and one published individual from the Blätterhöhle cave (I1565), both with >75% hunter-gatherer ancestry, and also some Wartberg-associated individuals with a range of WHG ancestry reaching as low as 25%. The fact that this relatively high WHG ancestry extended not just to the Rhine-Meuse wetlands, but also further along the Rhine and Meuse rivers and the northern coast, is consistent with archaeological evidence of continued cultural engagement of people across this region^42^. Three individuals from Tiel Medel who can be indirectly dated to ∼3700 BCE (Supplementary S2; Supplementary Table 7) form an exception to this pattern, with only ∼20% WHG ancestry (Figure 3), possibly representing new arrivals from neighboring parts of northwest Europe with lower WHG-associated ancestry.

**Figure 2.**
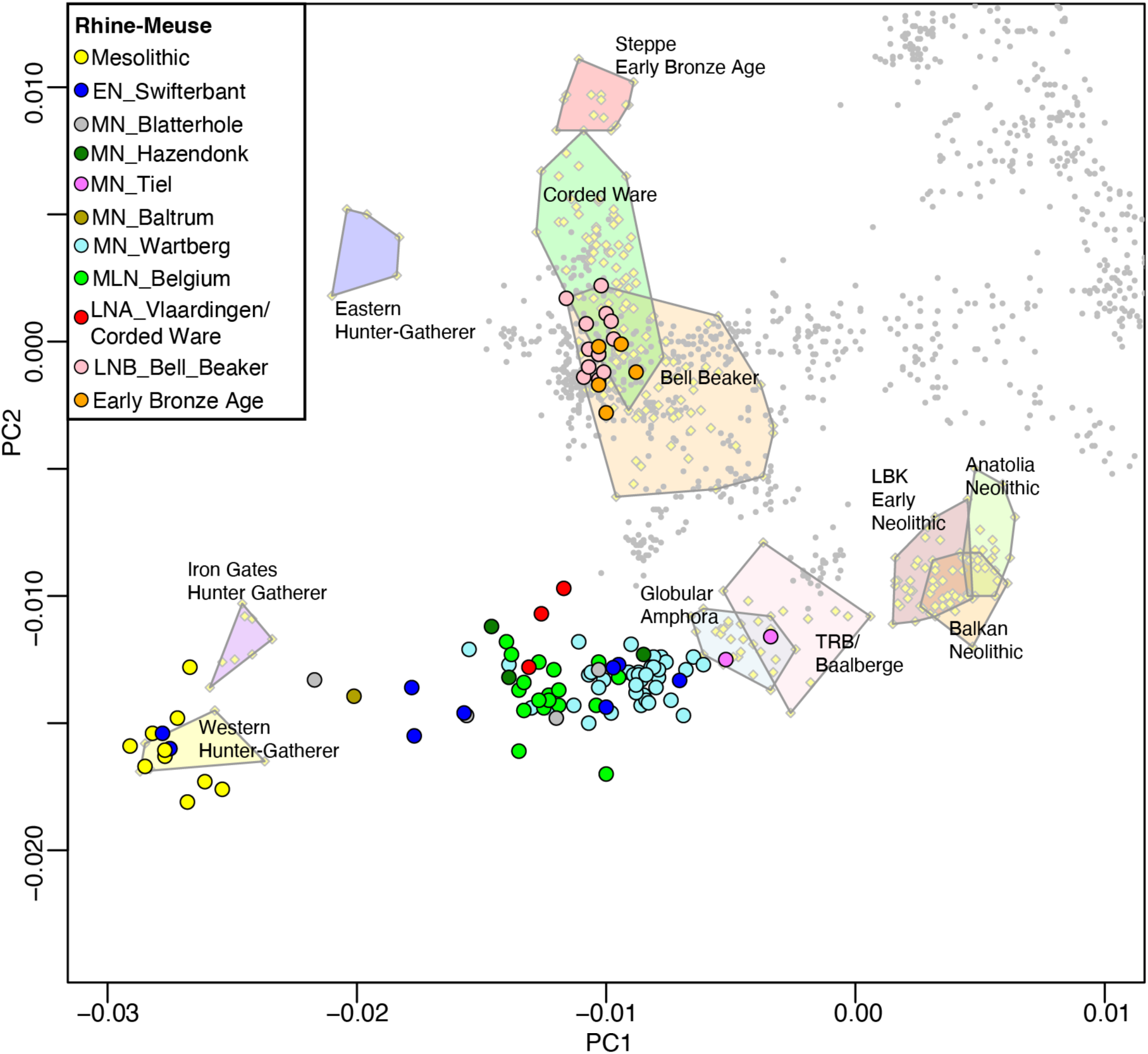
Principal Component Analysis with the ancient individuals projected onto the principal components computed on present-day individuals from West Eurasia.

**Figure 3.**
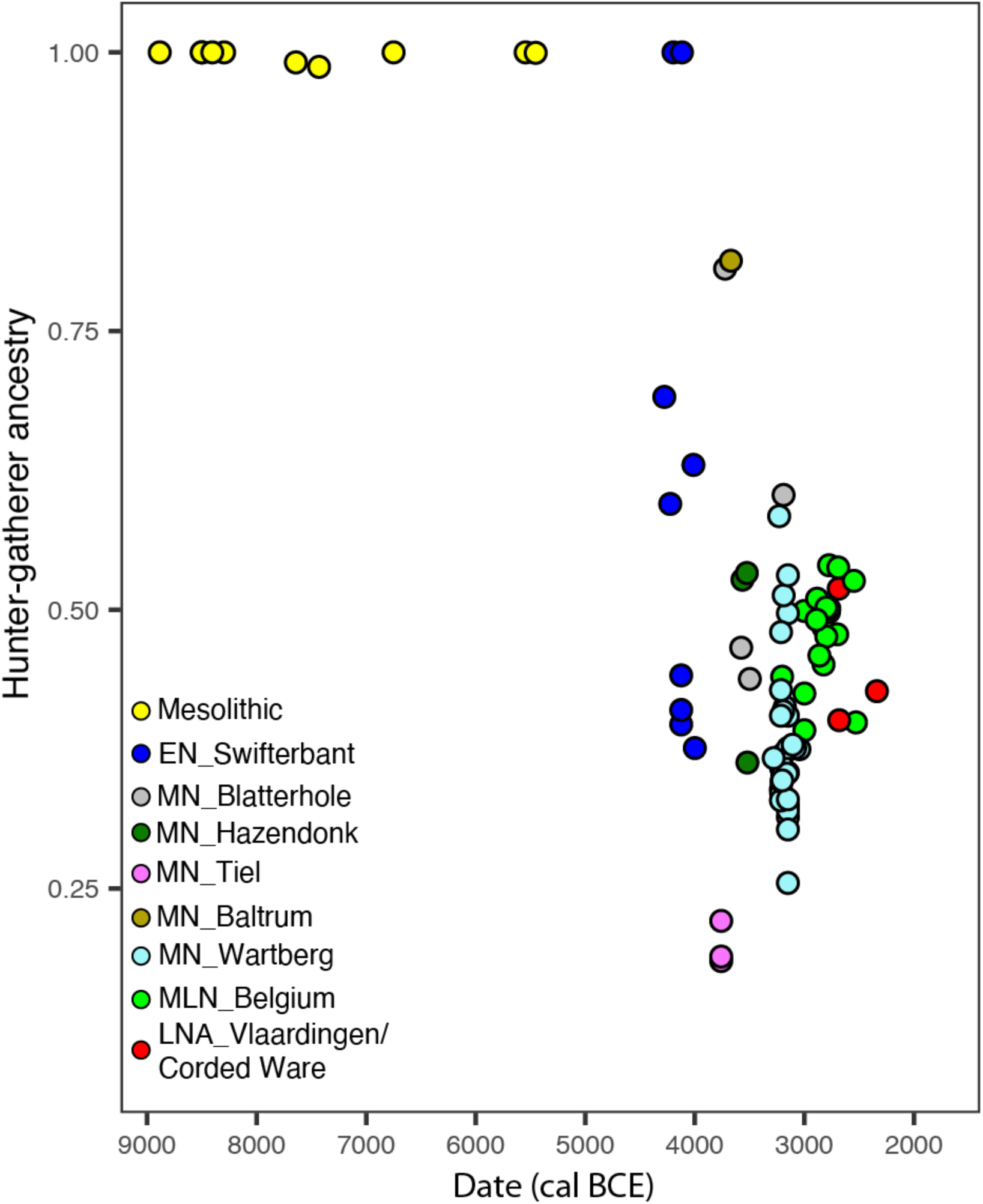
Hunter-gatherer ancestry proportions across time in the Rhine-Meuse area, estimated using *qpAdm* (Supplementary Table 3).

Compared to other regions of central, southern, and western Europe where farming was practiced (Figure 4), the Rhine-Meuse area stands out for its long survival of high proportions of WHG-related ancestry on a population scale (as opposed to isolated cases^43–45)^ until the BB transition, halfway through the 3^rd^ millennium BCE. To identify other instances where WHG ancestry on a whole-population scale endured in such high proportions to the dawn of the Bronze Age, it is necessary to go to cultures where farming was never adopted: to parts of the Baltic coast where populations with high EEF ancestry never made significant impact^12^, and to Scandinavia where hunter-gatherers with full WHG ancestry persisted until the early 3^rd^ millennium BCE alongside EEF-rich farmers^13^ (Figure 4).

**Figure 4.**
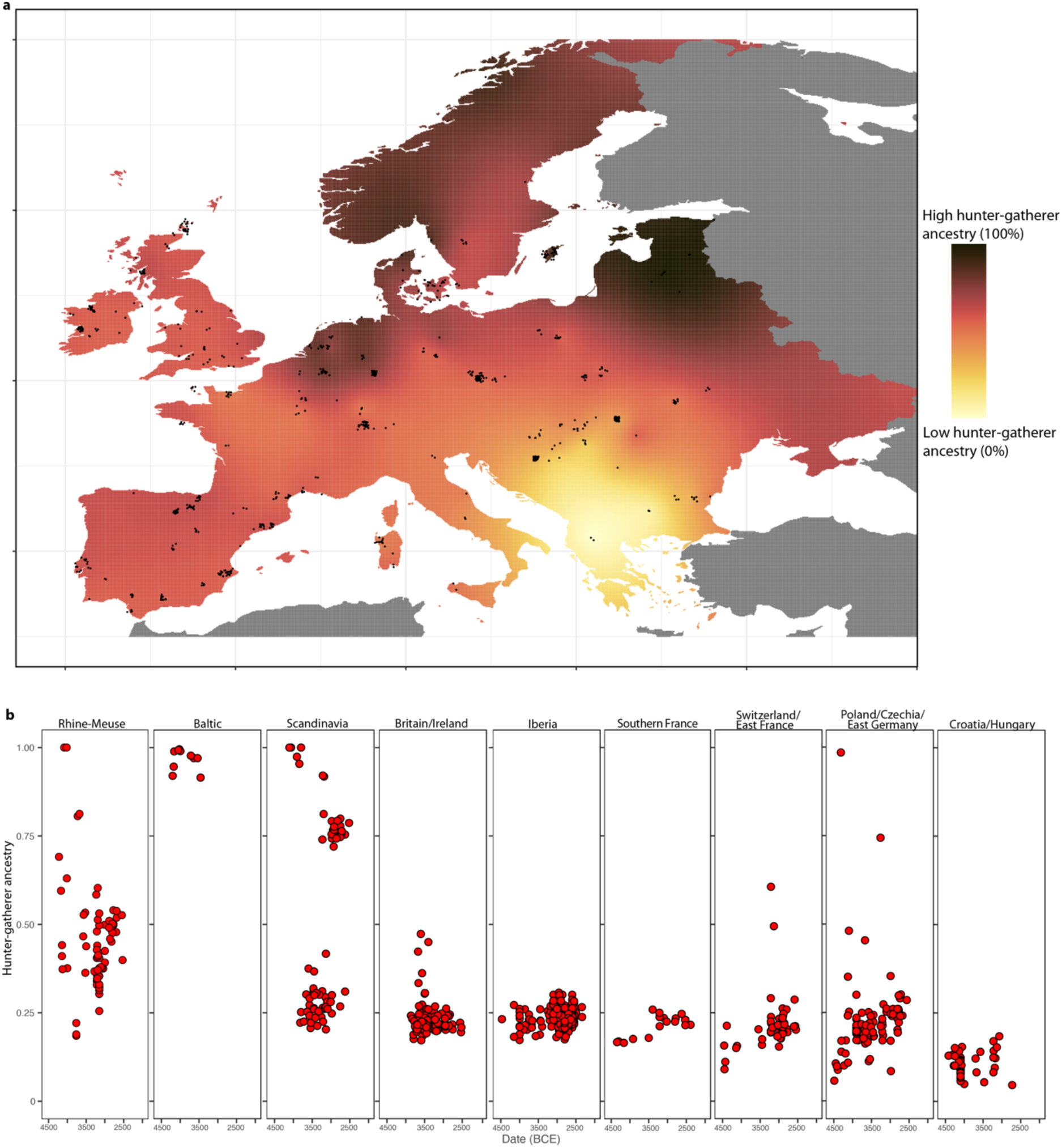
Hunter-gatherer ancestry proportions across Europe between 4500-2500 BCE, estimated using *qpAdm*. **a)** Spatial kriging of hunter-gather ancestry. The colors represent the predicted ancestry proportion at each point in the grid. **b)** Hunter-gatherer ancestry levels in individuals from different European regions.

## A genetically interconnected region with female-mediated gene flow

We find that the EEF ancestry proportions in Rhine-Meuse area Neolithic people were significantly higher on chromosome X than the autosomes (normally distributed Z-score between 2 and 3) (Supplementary Table 4), indicating a higher ancestral contribution from women with EEF ancestry. Independent confirmation is provided by analysis of the two uniparentally inherited parts of the genome (Supplementary Table 6). All Neolithic men (*n*=42 excluding close relatives) belong to Y-chromosome lineages common in Mesolithic hunter-gatherers (haplogroups I2a, R1b-V88 and C1a). In contrast, the maternally transmitted mitochondrial lineages are predominantly of Neolithic farmer origin (50 out of 71, based on their absence in sampled European Mesolithic individuals^2,6,12,13,38,45–57^. For example, the earliest individual with EEF ancestry, a Swifterbant female dated to ∼4342-4171 cal BCE (I17968, Nieuwegein) at the start of the transition to farming in the region^19,21^, harbors only 30% EEF ancestry in her autosomes but farmer-associated mitochondrial haplogroup H+152. A previous study^57^ reported similar sex-biased admixture in Neolithic farmers of Iberia and in Funnel Beaker farmers of northern Europe. A plausible scenario is that in all three regions, hunter-gatherer communities incorporated farmer women, who plausibly acted as vectors for exchange of ideas and technologies related to farming. This scenario of sex-biased admixture of Neolithic ancestry contrasts with one of complete displacement of local cultures by incoming farmers and migration of entire groups, a process that occurred in other parts of early Neolithic Europe.

The Neolithic populations of the Rhine-Meuse region were highly genetically interconnected, as reflected in large segments (>12 cM) of the genome being identical by descent (IBD), which is only expected to be observed for individuals who share common ancestors in the last dozens of generations^58^ (Supplementary Table 7). We also find several cases of IBD segments >20 cM, suggesting even closer relationships between sites such as Blätterhöhle, Niedertiefenbach, and Abri Sandron, as well as between sites in the Rhine-Meuse area and nearby areas of central Europe and Northern France (Figure 5). A striking case is a relationship (∼50 cM in IBD) between an individual from Blätterhöhle, modern western Germany, and a father-daughter pair from Mont-Aimé^44^ in modern northern France, who are also clear ancestry outliers exhibiting more hunter-gatherer ancestry than other individuals from Mont-Aimé.

**Figure 5.**
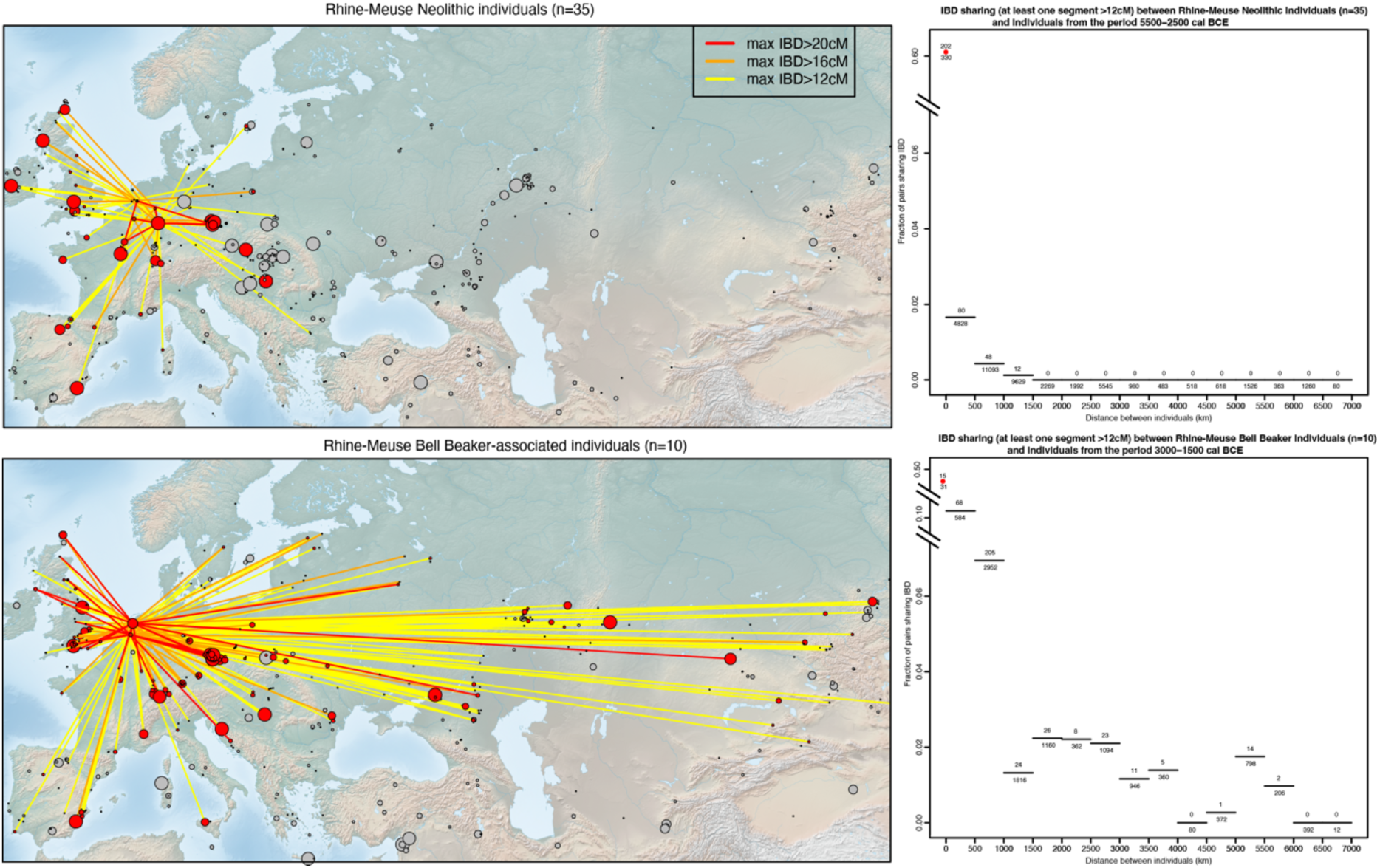
On the left, IBD connections of Rhine-Meuse Early, Middle and Middle-Late Neolithic individuals (top) and Rhine-Meuse BB-associated individuals (bottom). Sites are represented by circles with size proportional to the number of individuals amenable to IBD calling. Grey circles indicate archaeological sites between 5500-2500 BCE (top) and 3000-1500 BCE (bottom) with no IBD connections to Rhine-Meuse individuals. On the right, decay of IBD connections with geographic distance for Rhine-Meuse Early, Middle and Middle-Late Neolithic individuals (top) and Rhine-Meuse BB-associated individuals (bottom).

### Minimal steppe admixture associated with the appearance of Corded Ware pottery

In many areas of Europe, the emergence of the CW complex is associated with large-scale demographic change due to the arrival of groups carrying steppe ancestry. The ancestry change in three sampled individuals from Vlaardingen/CW contexts in the Western Netherlands is far smaller. These individuals were buried within settlements that used CW complex material goods, but they were not part of the typical CW single grave burials, which are absent from the Vlaardingen culture region. One female (I12896 from the site Molenaarsgraaf) is consistent with having no steppe-related ancestry at all and instead sharing ancestry with local late Neolithic farmers of the region. However, the other two individuals, from Mienakker and Sijbekarspel Op de Veken, north of the Rhine River delta, can be modeled as a mixture of ∼11% ancestry associated with the main cluster of CW groups, and 89% derived from Rhine-Meuse area Neolithic populations with high hunter-gatherer ancestry, similar to Late Neolithic Belgium (Supplementary Table 5). Despite a low steppe ancestry proportion in the autosomes, the male I12902 from Mienakker, who yields one of the earliest CW complex associated dates on bone in Europe outside of Bohemia and the Baltic region (2852-2574 cal BCE), carries Y haplogroup R1b-U106, also known in one early CW-associated individual from Bohemia^7^. These results suggest that the male ancestor who brought this Y haplogroup to the Rhine-Meuse region was part of the early CW expansion.

While limited to three people, the IBD analysis for the Vlaardingen/CW individuals revealed two additional notable signals. First, the two individuals with ∼11% CW ancestry, excavated at nearby sites, have an IBD match that represents approximately a 3^rd^ cousin relation, hinting at a small community size. Second, the geographical range of their IBD links extends much further east than previous groups (Extended Data Figure 1). Among their closest connections (one segment of 19cM) is a Yamnaya-associated individual from Samara in far eastern Europe^59^ (ID: I6733) and CW-associated individuals from present-day Poland^60^ (ID: pcw362) and Czechia^7^ (ID: STD002). We also detect connections to other Rhine-Meuse area Late Neolithic individuals, providing direct evidence of a major local ancestry component in Vlaardingen/CW individuals. These patterns reflect their dual sources of ancestry: a minor component (potentially completely along the male line) from central/eastern CW groups (and through them to Yamnaya steppe pastoralists) and a major component from the local Neolithic population.

## Bell Beaker complex: population influx and local admixture in the Rhine-Meuse delta

The advent of the BB complex in the Rhine-Meuse delta after 2500 BCE is associated with a demographic change of a much larger scale than events in the previous millennia. All 13 available BB-associated individuals appear genetically close to CW-associated groups, but not to the preceding local Rhine-Meuse area Vlaardingen/CW individuals in the PCA. They can be modeled with ∼83% ancestry from the main cluster of CW-associated individuals (Supplementary Table 5), and their remaining ancestry from a Wartberg Neolithic-related group, representing the Neolithic population from the Rhine-Meuse area with the lowest level of hunter-gatherer ancestry, or as a mixture between Middle and Late Neolithic groups from outside the Rhine-Meuse area (*e.g.* Poland Globular Amphora, Czechia TRB, Germany Baalberge, Iberia Neolithic-Chalcolithic) and Late Neolithic populations from Belgium. All these scenarios point to a ∼83-91% (but not 100%) ancestry change associated with the arrival of the BB complex in the Rhine-Meuse region (Supplementary Table 5). A contribution of local Rhine-Meuse delta farmers, with their distinctive signature of high hunter-gatherer ancestry, is essential to model the formation of Rhine-Meuse delta BB-associated individuals. Even at 9-17%, we can be confident this local admixture occurred: models lacking this unique Rhine-Meuse delta farmer genetic contribution are rejected with high significance. This suggests that the observed mixture between CW culture associated groups and European farmers that formed the genetic profile of Rhine-Meuse delta BB must have occurred in the region itself. Radiocarbon dates further suggest that the Rhine-Meuse area was one of the earliest places where the BB cultural phenomenon arose^61^. While the earliest appearance of BB cultural material has been located in Iberia^62^, our results show that early formation of BB-associated groups, influenced not just culturally but also genetically by CW users, also occurred in the Rhine-Meuse area.

The advent of the BB complex in the Rhine-Meuse region contrasts to the previous ancestry changes in our time transect, as it was more transformative in a demographic sense, with large ancestry turnover involving both sexes. Among BB men, all yielded R1b-L151 haplogroups, which were absent in earlier Neolithic European populations but present in early CW-associated individuals from Czechia in central Europe. All nine available BB-associated men from Oostwoud and Ottoland-Kromme Elleboog belonged to the derived R1b-L151-P312 lineage, the major lineage among BB groups across Europe. Two P312 individuals could be further subtyped to DF19, a minor P312 subtype today mostly present in central/northern European populations (https://www.yfull.com/tree/R-DF19/). At Molenaarsgraaf, the only man with enough resolution to determine a L151 subtype belonged to R1b-L151-U106 (I13025), matching the Vlaardingen/CW associated male from Mienakker, referred to above, and suggesting a similar CW-related source for the patrilineal ancestry in both Vlaardingen/CW and BB men in the Rhine-Meuse area. This would be consistent with the hypothesis that even if there was limited local continuity within the lowlands, the same male lineages that were associated with the arrival of CW pottery to the region (in the Vlaardingen/CW context) were at least partially associated with subsequent BB emergence. This suggests the possibility that the BB associated population in the Rhine Meuse delta emerged from sustained influx of ‘classic’ CW-related groups to the region (such as those documented in the uplands to the east, where skeletal preservation is absent but classic CW burial features are present), which mixed with the local Vlaardingen or other Rhine-Meuse Neolithic populations that had high hunter-gatherer ancestry.

Further evidence for a major influx from outside regions during the BB period comes from inspection of IBD networks, which, after this point, expand thousands of kilometers further to the east and northeast (Figure 5). These show strong links to CW and BB-associated individuals from Bohemia, as well as evidence of distant Early Bronze Age relatives from England and Scotland, corroborating findings of a shared origin between these groups located on opposite shores of the North Sea^6^. The BB horizon is the first period in our time transect when people of the Rhine-Meuse region became intensively integrated within a much wider European IBD network, in contrast to previous more regional patterns, yet another indication for the high level of mobility that has become evident from isotope studies^63,64^.

After this period of profound demographic change, Early Bronze Age individuals from the Rhine-Meuse area had similar ancestry to BB predecessors (Figure 2), with ∼6% additional Neolithic-related ancestry (Supplementary Table 5), potentially reflecting small-scale admixture with neighboring populations. Local continuity from the BB period to the Middle Bronze Age is also reflected in the abundant familial links connecting Early Bronze Age individuals with both earlier BB individuals and later Middle Bronze Age ones (Supplementary Table 7), with pairs sharing up to 100 cM IBD as expected for ∼6th-degree relations.

## Expansion from the Rhine-Meuse area led to disruptive demographic change in Britain

The study highlights the Rhine-Meuse delta region as a candidate for a secondary expansion of BB users that had a greater demographic impact than the postulated initial Iberian expansion. To test this hypothesis, we used our statistical modelling framework to re-examine the genetic evidence of the arrival of BB users in Britain. Previous work showed a minimum ∼90% ancestry change, but had not been able to distinguish models in which the EEF admixture came from Britain itself or elsewhere^6^. We analyzed 30 BB-associated individuals from Great Britain from the main homogeneous cluster and obtained the same result as for Rhine-Meuse delta BBs (Supplementary Table 5; Table 1). Both require ∼9-20% ancestry from Rhine-Meuse delta Neolithic populations with high levels of hunter-gatherer ancestry, plus ancestry from the main CW genetic cluster. This is consistent with no contribution at all from British Neolithic farmers. However, in an outlier subset of four BB associated individuals from Britain (including the high-status “Amesbury Archer”) with lower proportions of steppe ancestry than the main cluster, models featuring CW Complex associated groups and Rhine-Meuse farmers provide a poor fit. We cannot rule out the possibility that some of the ancestry of these outliers could derive from local British Neolithic populations, but it could also plausibly come from separate migratory streams into Britain, such as the one suggested for the Amesbury Archer whose isotopic genetic signatures indicate an origin in the Alps^65^. By the Early Bronze Age, when ancestry proportions in Britain stabilized, we estimate 91% Rhine-Meuse BB ancestry and at most 9% local British Neolithic ancestry but possibly as little as 0%, providing new information about the magnitude of the demographic transition associated with the BB transition in Britain.

**Table 1.** Modelling the ancestry of Rhine-Meuse BB-associated individuals and the main group of British BB-associated individuals using *qpAdm* models involving a CW group from present-day Germany and different Neolithic populations from within and outside the Rhine-Meuse area. See Supplementary Table 5 for further details.

|  | **Rhine-Meuse BB** | **British BB** |
| --- | --- | --- |
| P-value for models with Iberian farmers | 0.0073 | 0.0104 |
| P-value for models with French farmers | 0.0000068 | 0.0000011 |
| P-value for models with German farmers | 0.00022 | 0.00026 |
| P-value for models with Globular Amphora farmers | 0.0028 | 0.0028 |
| P-value for models with MLN Belgium (Rhine-Meuse) | 0.02060 | 0.00056 |
| P-value for models with MN Wartberg (Rhine-Meuse) | 0.62 | 0.69 |
| Proportion of ancestry from LN Wartberg | 17.4% | 19.6% |
| Minimum proportion of ancestry from MLN Belgium allowing 3-way *qpAdm* models with CW and other farmer sources outside the Rhine Meuse area | 9.4% | 8.8% |

## Discussion

A striking finding in our study is that the long-term persistence of high hunter-gatherer ancestry was not limited to the core wetlands of the Netherlands, but was also a feature of inland regions of the Rhine (Blatterhöhle cave and Wartberg culture) and Meuse (Belgian Neolithic sites). In the context of archaeological evidence, this suggests the co-existence of two distinct but interdigitated Neolithic spheres in this region, persisting well into the 3^rd^ millennium BCE. One was the communities centered around water, not only those in the wetland core but also connected to it through waterways, suggesting the presence of semi-agrarian groups far inland^17,66^. Water links were central for these communities, and connections to people living along the waterways were often more important than connections to physically closer neighbors. Surrounding the waterways, early farming communities preferred the fertile loess soils and kept to culturally specific traditions of settlement, housing, material culture, and burial. This supports the “frontier mobility” model proposed by Zvelebil^67^, albeit in a more geographically restricted context. These communities exchanged ideas, women introduced EEF genetic ancestry and probably also new technological knowledge in the more hunting-gathering practicing communities, but their distinct cultures persisted for millennia.

The genetic distinctiveness of the waterway-based people through the Neolithic began with the Swifterbant and Hazendonk communities, continued into the Vlaardingen and Seine-Oise- Marne groups, and persisted through the time when the CW complex began to influence the region. CW pottery is present in the Rhine-Meuse delta, but other aspects of this culture are lacking, in particular the characteristic burial rituals^27,34,35^. This was accompanied by limited population influx and high retention of the ancestry of the previously established groups.

The advent of the BB complex marked a break from the previously established pattern, with evidence of whole-group migration involving people of both sexes along with some degree of admixture with local Rhine-Meuse delta populations (9-17%). In our transect east of the Rhine-Meuse delta, there is a ∼300 year gap between the three analyzed Vlaardingen/CW-associated individuals and the 13 BB-associated individuals. Thus, the time course and exact location in the region where people with a CW-associated genetic profile and a Rhine-Meuse Neolithic genetic profile came together remains unclear. However, the homogeneity of ancestry within the three BB-associated sites we sampled show that the mixture had largely taken place by that time (∼2300-1900 BCE) and that the cumulative effect was transformative.

The evidence for a large-scale demographic change in the Rhine-Meuse region by the time of the spread of Beaker users is important in light of the evidence from Britain, where Beaker users spread at around the same time. Since the British Neolithic populations encountered by Beaker users practiced cremation and thus did not yield samples amenable for aDNA analysis, it has been unclear whether there was a sharp population break or whether there was a period of extended co-existence ^68^. In the Rhine-Meuse region, in contrast, the practice of cremation by previously established groups was infrequent and the turnover was very substantial. This raises the plausibility of a similar process in Britain, where our analyses show an even more profound demographic change, with the great majority of local BB burials being consistent with no local British Neolithic ancestry and the contribution of the local British Neolithic population by the Early Bronze Age estimated at ≤9%. While we do not know what triggered this large-scale mobility, the genetic legacy of local populations both in the Rhine-Meuse area and Britain collapsed relatively rapidly. This could have been facilitated by local populations already in decline due to agricultural crisis or epidemic events^69,70^, but also by inter-group violence^71^.

Despite the evidence of a cultural break, in both the Rhine-Meuse area and Britain, important “indigenous” cultural traditions and knowledge remained intact. In Britain, the BB immigrants continued to use particular traditional sources of (Langdale) flint^72^, continued to build and alter Stonehenge and other monuments, to build round houses, and practiced forms of agriculture that were similar to the earlier Grooved Ware complex^73^. In the Rhine-Meuse area, BB groups used the same settlement areas, and continued to settle in river valleys, on crevasse splays, and along river dunes in a way that was oriented explicitly towards a hunting-farming mixed economy, highlighting continuity as well as change in this unique part of Europe.

## Methods

### Sampling, extraction, library preparation, capture, sequencing

Our initial selection for aDNA analysis included 118 ancient individuals for aDNA analysis, including one previously reported individual for whom we generated additional libraries. We performed laboratory work in dedicated clean rooms. We removed the outer layer of teeth and long bones, and collected powder from beneath the cleaned surface. This process minimized the risk of exogenous DNA contamination, with low-speed drilling used to prevent heat-induced DNA damage^74^. In the case of temporal bones, we removed cochleae through sandblasting^75^, and then milled them. We incubated the resulting powder in lysis buffer, and cleaned and concentrated the DNA from one-fifth of the lysate. We did this either manually or using an automated protocol with silica magnetic beads^76^ and Dabney Binding Buffer^77,78^ for manual extraction. The samples from Trou Al’Wesse, Abri Sandron, and Grotte du Mont Falise were prepared and extracted following the method outlined in Dulias *et al*.^79^.

We built 219 libraries (Supplementary Table 2) using two different protocols. Double-stranded barcoded libraries were prepared with truncated adapters from the extract and subjected to partial (“half”) uracil–DNA–glycosylase (UDG) treatment before blunt-end repair to significantly reduce the characteristic damage pattern of aDNA^80,81^. Single-stranded libraries were prepared using automated scripts following Gansauge *et al*. ^82^. A fraction was subjected to USER treatment.

DNA libraries were enriched for human DNA using probes that target 1,233,013 SNPs (‘1240k capture’^83^) or 1,352,535 SNPs (‘Twist’ BioSciences^84^), and the mitochondrial genome. We performed two rounds of capture for the ‘1240k’ reagent and one for the ‘Twist’ BioSciences reagent. Captured libraries were sequenced on an Illumina HiSeq X10 instrument with 2x101 cycles and 2x7 cycles to read out the two indices^85^ or on an Illumina NextSeq 500 instrument with 2x76 cycles and 2x7 cycles to read out the two indices.

### Bioinformatics: Demultiplexing, adapter removal, mapping, PCR duplicate removal

Reads for each sample were extracted from the raw sequencing data based on sample-specific indices introduced during wet-lab processing, permitting up to one mismatch. Adapters were removed, and paired-end sequences were merged into single-ended sequences with a required 15-base-pair overlap (allowing one mismatch with high quality bases or three mismatches with low quality bases). This process was applied by selecting the highest-quality base in the overlapping region. Reads that could not be merged were discarded before aligning to the human reference genome (hg19) and the RSRS version of the mitochondrial genome using the ‘samse’ command in bwa (version 0.6.1)^86^. We removed duplicates based on the alignment coordinates and orientation of the aligned reads. Aligned sequences from different libraries of the same sample were merged accordingly into a single bam file. The computational pipelines are available on GitHub (https://github.com/dReichLab/ADNA-Tools, https://github.com/dReichLab/adna-workflow).

### Evaluation of authenticity

We established aDNA authenticity using several criteria. Libraries with a deamination rate below 3% at the terminal nucleotide were excluded from further analysis. We computed the ratio of Y-chromosome to X- and Y-chromosome reads. Libraries with ratios above 0.03 and below 0.32 were excluded from further analysis. We estimated mismatch rates to the consensus mitochondrial sequence using contamMix- 1.0.1051^87^, and X-chromosome contamination estimates using ANGSD^88^ in males with sufficient coverage. Libraries with evidence of contamination were excluded from further analysis. Finally, individuals without a minimum of 20,000 targeted 1240k SNPs with at least one overlapping sequence were discarded from population genetic analysis. After applying these filters, 86 libraries from 42 individuals remained, and we merged data from the libraries to increase sequencing coverage.

### Analysis datasets

In addition to the 41 individuals with newly generated data and the previously reported individual from Oostwoud^41^ with additional data, we also included data from 67 previously published individuals^6,10,38–41^ from the Rhine-Meuse region, dated from the Mesolithic to the Early Bronze Age, for a total of 109 individuals from the Rhine-Meuse region and adjacent areas between 8500-1700 BCE (Figure 1; Supplementary Table 1). The time-transect dataset includes new Mesolithic individuals from Belgium, the Netherlands and northwest Germany, as well as published data from now submerged areas of Doggerland^38^; nine Early-Middle Neolithic individuals from semi-agrarian Swifterbant contexts (4500-4000 BCE) (Netherlands) - the first data from this unique culture; the first three Middle Neolithic individuals from Hazendonk archaeological contexts (Netherlands); three likely Middle Neolithic individuals from Tiel (Netherlands) and one Middle Neolithic individual from Baltrum island (northwest Germany); four published Middle Neolithic individuals from Blätterhöhle cave (3500-3000 BCE)^10^ (northwest Germany); 40 published Middle Neolithic individuals from a Wartberg context (3500-2800 BCE)^39^ (Niedertiefenbach, northwest Germany); 18 Late Neolithic individuals buried in caves from the Ardennes region (3300-2500 BCE) (Belgium); three Late Neolithic individuals from Vlaardingen/CW contexts including the first data from this culture from the Rhine-Meuse area (3000-2500 BCE) (Netherlands); 13 Late Neolithic individuals from a BB context (2500-2000 BCE) (Netherlands); and five individuals from Early Bronze Age contexts (2000-1700 BCE) (Netherlands).

To aid the analysis of the Rhine-Meuse area individuals, the analysis dataset was further complemented by previously published data from ancient individuals from other regions. For genome-wide analyses, we assembled two datasets. The HO dataset included the ancient individuals, and 1,036 present-day West Eurasian individuals genotyped on the Affymetrix Human Origins Array^2,89,90^. We kept 591,642 SNPs shared between the 1240k capture and the Human Origins Array. The HOIll dataset included only the ancient individuals and 1,233,013 SNPs in common between 1240k and Twist reagents. In both datasets, we randomly sampled one allele at each SNP position for each individual, discarding the first and the last two nucleotides of each sequence.

### Haplogroup assignment of uniparentally inherited markers

We created consensus mitochondrial haplotypes with samtools and bcftools. We restricted to sequences with a mapping quality of more than 30 and a base quality of more than 30. We then called haplogroups with Haplogrep3 (Supplementary Table 1). We called Y-chromosome haplogroups (Supplementary Table 1) following the methodology in Lazaridis *et al*.^91^, based on the YFull v.8.09 phylogeny (https://www.yfull.com/). Haplogroups found in Neolithic individuals were classified as either “hunter-gatherer-related” if they were already present among Mesolithic hunter-gatherers (mitochondrial haplogroups U5, U4’9, U2, U* and K1e; Y-chromosome haplogroups I2a, C1a, and R1b-V88), or “Neolithic related” if they were most likely brought by incoming Early European farmer populations (all mitochondrial and Y-chromosome haplogroups except those mentioned above ) (Supplementary Table 6). With this approach, we understand that we might be underestimating the number of lineages contributed by Neolithic farmers, both in the mtDNA and Y-chromosome, as some lineages considered as Mesolithic hunter-gatherer-related, based on their presence during the Mesolithic period, might have been incorporated by farmer populations during their path from Anatolia to the Rhine-Meuse area.

### Molecular sex determination

Genetic sex was determined by calculating the ratio of reads mapped to Y-chromosome SNP positions to the total reads mapped to sex-chromosome SNP positions. Individuals with a ratio <0.03 were classified as female, while those with a ratio >0.32 were classified as male.

### Biological relatedness

To estimate close biological relatedness up to the 3^rd^ degree, mismatch rates were computed between all possible pairs of Rhine-Meuse area individuals, randomly sampling one read for each individual at each of the 1.15 million autosomal SNPs. Mismatch rates were converted to relatedness coefficients following Fowler *et al*.^92^, using three different baseline mismatch rate values to account for the different ancestral backgrounds found in the dataset. If both individuals in the pair had fully Mesolithic hunter-gatherer ancestry, we use a baseline mismatch rate of 0.225. If at least one individual has EEF ancestry and both lack steppe-associated ancestry, we use a baseline mismatch rate of 0.252. If at least one individual has steppe-associated ancestry, we use a baseline mismatch rate of 0.258. Close kinship relationships are annotated in Supplementary Table 1.

### Runs of homozygosity

We called runs of homozygosity using hapROH^93^ for individuals with more than 300,000 available SNPs.

### Principal component analysis

We ran Principal Component Analysis on the HO dataset using the smartpca software from the EIGENSOFT package^94^. We computed PCs on 1036 present-day West Eurasians genotyped on the Affymetrix Human Origins Array. Ancient individuals were projected onto those PCs using lsqproject:YES and shrinkmode:YES.

### Data Access

Genotype data for individuals included in this study can be obtained from the Harvard Dataverse repository through the following link (https://xxx). The DNA sequences reported in this paper are deposited in the European Nucleotide Archive under accession number xxx. Other newly reported data, such as radiocarbon dates and archaeological context information, are included in the manuscript and supplementary files.

## Acknowledgments

We thank Nicole Adamski, Nasreen Broomandkhoshbacht, Elizabeth Curtis, Ilana Greenslade, Kristin Stewardson, and Fatma Zalzala for laboratory work. We thank Provinciaal depot voor archeologie van Noord-Holland (Martin Veen, Rob van Eerden), Provinciaal archeologisch depot Zuid-Holland (Inge Riemersma, Mark Phlippeau), Archeologisch Depot Gelderland (Stephan Weiss-König), and National Museum of Antiquities Leiden (Luc Amkreutz) for granting permission to sample ancient remains and assistance in sampling. The ancient DNA data generation and analysis was supported by the National Institutes of Health (R01-HG012287); the John Templeton Foundation (grant 61220); a private gift from Jean-Francois Clin; by the Allen Discovery Center program, a Paul G. Allen Frontiers Group advised program of the Paul G. Allen Family Foundation; by the Howard Hughes Medical Institute (DR); by the Max Planck Society, by the French Research Foundation and German Research Foundation (to M.R., W.H. and M.-F.D., project INTERACT, ANR-17-FRAL-0010 and DFG-HA-5407/4-1, 2018-21); and The Leverhulme Trust (AF, FG, CJE, MBR, MP). The accepted version of this article (before the editing, proofreading, and formatting changes following the paper being accepted) is subject to the Howard Hughes Medical Institute (HHMI) Open Access to Publications policy; HHMI lab heads have previously granted a non-exclusive CC BY 4.0 license to the public and a sublicensable license to HHMI in their research articles. Pursuant to those licenses, the accepted manuscript can be made freely available under a CC BY 4.0 license immediately upon publication.

## Author Contributions

I.O., E.A., Q.B., H.F. and D.R. wrote the manuscript with the input of all other coauthors. M.B.R., R.P., W.H., M.P. and D.R. supervised parts of the study. I.O., I.L., N.P. and M.R. analysed genetic data. E.A., Q.B. and H.F. edited archaeological information. L.A., M.-F.D., A.F., D.F., F.G., J.F.K., L.M.K., K.L., L.L.K., R.L., R.M., H.M., P.N., D.C.M.R., M.R., L.S., J.R.S., T.tenA., M.T.. C.J.E. sampled anthropological remains and/or contributed to the creation of the archaeological supplement. G.S., M.M., A.M. and S.M. processed bioinformatic data. K.C., O.C., T.F., L.I., J.O., I.P., L.Q., J.N.W., C.J.E. and N.R. carried out wet laboratory work.

## Ethics Statement

The individuals studied here were all analyzed with the goal of minimizing damage to their skeletal remains, with permission from local authorities in each location from which they came. Every sample is represented by stewards, such as archaeologists or museum curators, who are either authors or thanked in the Acknowledgments. Open science principles require making all data used to support the conclusions of a study maximally available, and we support these principles here by making fully publicly available not only the digital copies of molecules (the uploaded sequences) but also the molecular copies (the ancient DNA libraries themselves, which constitute molecular data storage). Those researchers who wish to carry out deeper sequencing of libraries published in this study should send a request to the corresponding author DR. We commit to granting reasonable requests as long as the libraries remain preserved in our laboratories, with no requirement that we be included as collaborators or co-authors on any resulting publications.

---

**Extended Data Figure 1.**
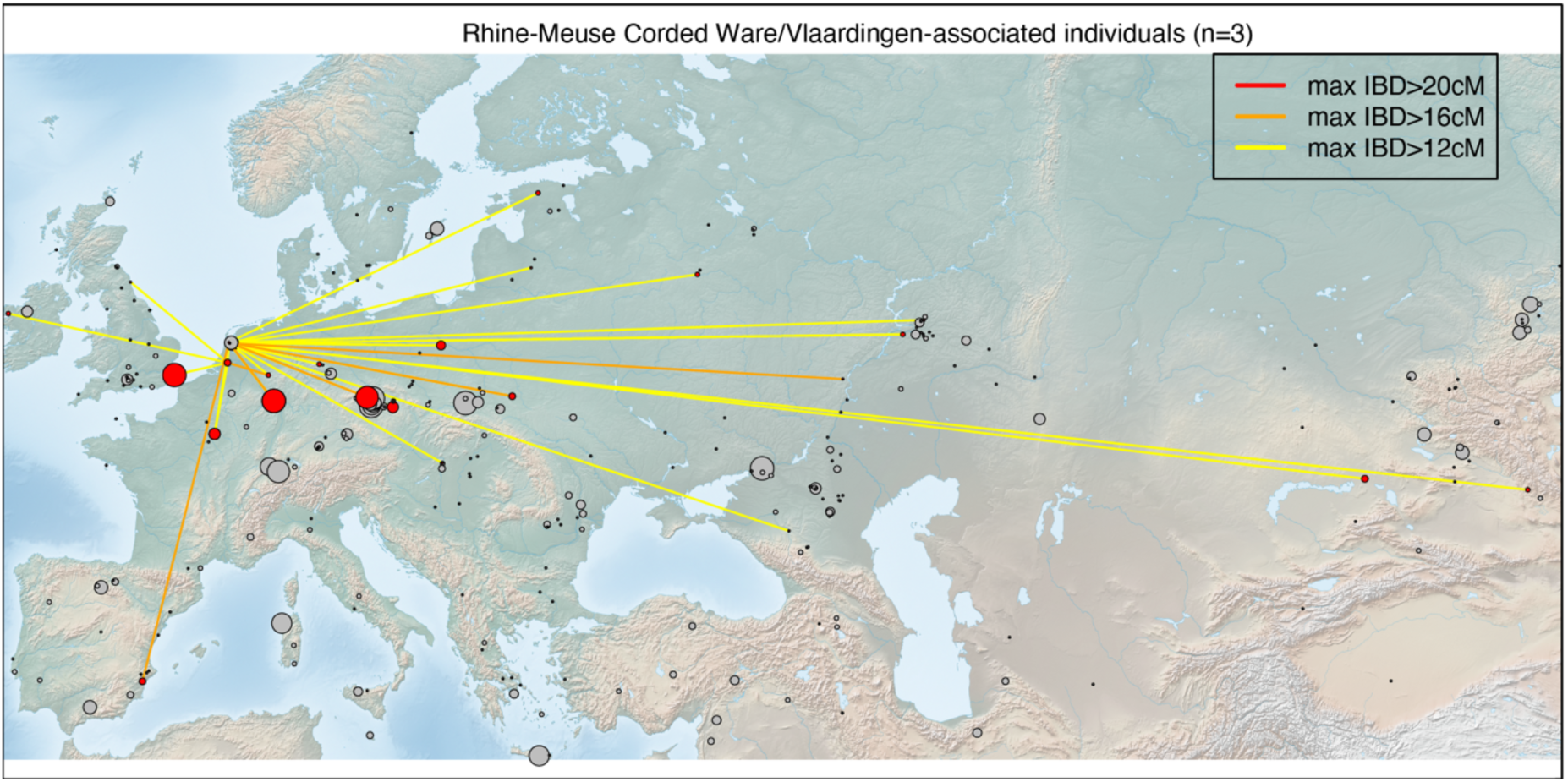
IBD connections of Rhine-Meuse CW/Vlaardingen individuals. Circles represent archaeological sites, with size proportional to the number of individuals amenable to IBD calling. Grey circles indicate archaeological sites between 3000-2000 BCE with no IBD connections to CW/Vlaardingen individuals.

## References

1. Olalde, I. et al. A common genetic origin for early farmers from Mediterranean cardial and Central European LBK cultures. Mol. Biol. Evol. 32, 3132–3142 (2015).

2. Lazaridis, I. et al. Ancient human genomes suggest three ancestral populations for present-day Europeans. Nature 513, 409–413 (2014).

3. Skoglund, P. et al. Origins and genetic legacy of Neolithic farmers and hunter-gatherers in Europe. Science 336, 466–469 (2012).

4. Haak, W. et al. Massive migration from the steppe was a source for Indo-European languages in Europe. Nature 522, 207–211 (2015).

5. Allentoft, M. E. et al. Population genomics of Bronze Age Eurasia. Nature 522, 167–172 (2015).

6. Olalde, I. et al. The Beaker phenomenon and the genomic transformation of northwest Europe. Nature 555, 190–196 (2018).

7. Papac, L. et al. Dynamic changes in genomic and social structures in third millennium BCE central Europe. Sci. Adv. 7, (2021).

8. Brandt, G. et al. Ancient DNA reveals key stages in the formation of Central European mitochondrial genetic diversity. Science 342, 257–261 (2013).

9. Mathieson, I. et al. Genome-wide patterns of selection in 230 ancient Eurasians. Nature 528, 499–503 (2015).

10. Lipson, M. et al. Parallel ancient genomic transects reveal complex population history of early European farmers. Nature 551, 368–372 (2017).

11. Robb, J. Material Culture, Landscapes of Action, and Emergent Causation: A New Model for the Origins of the European Neolithic. Curr. Anthropol. 54, 657–683 (2013).

12. Mittnik, A. et al. The genetic prehistory of the Baltic Sea region. Nat. Commun. 9, 1–11 (2018).

13. Skoglund, P. et al. Genomic Diversity and Admixture Differs for Stone-Age Scandinavian Foragers and Farmers. Science 344, 747–750 (2014).

14. Malmström, H. et al. Ancient DNA Reveals Lack of Continuity between Neolithic Hunter-Gatherers and Contemporary Scandinavians. Curr. Biol. 19, 1758–1762 (2009).

15. Hofmann, D. The changing role of La Hoguette pottery in an LBK context. in 191–224 (2016).

16. Kirschneck, E. The Phenomena La Hoguette and Limburg – Technological Aspects. Open Archaeol. 7, 1295–1344 (2021).

17. 17. Amkreutz, L. W. S. W. Persistent traditions: a long-term perspective on communities in the process of Neolithisation in the Lower Rhine Area (5500-2500 cal BC). (Sidestone Press, 2013).

18. Crombé, P. et al. New evidence on the earliest domesticated animals and possible small-scale husbandry in Atlantic NW Europe. Sci. Rep. 10, 1–15 (2020).

19. Brusgaard, N. Ø. et al. Early animal management in northern Europe: multi-proxy evidence from Swifterbant, the Netherlands. Antiquity 98, 654–671 (2024).

20. Kooijmans, L. P. L. & Jongste, P. F. B. A neolithic settlement on the Dutch North Sea coast c. 3500 CAL BC. (Analecta Praehistorica Leidensia S., 2006).

21. Dreshaj, M., Dee, M., Brusgaard, N., Raemaekers, D. & Peeters, H. High-resolution Bayesian chronology of the earliest evidence of domesticated animals in the Dutch wetlands (Hardinxveld-Giessendam archaeological sites). PLoS One 18, 1–23 (2023).

22. Robb, J. Material Culture, Landscapes of Action, and Emergent Causation: A New Model for the Origins of the European Neolithic. Curr. Anthropol. 54, 657–683 (2013).

23. Menne, J. & Brunner, M. Transition from Swifterbant to Funnelbeaker: A Bayesian Chronological Model. Open Archaeol. 7, 1235–1243 (2021).

24. Goossens, T. . Opgraving Hellevoetsluis-Ossenhoek. Een nederzetting van de Vlaardingen-groep op een kwelderrug in de gemeente Hellevoetsluis. (Leiden (Archolrapport 87), 2009).

25. Furholt, M. Mobility and Social Change: Understanding the European Neolithic Period after the Archaeogenetic Revolution. Journal of Archaeological Research 29, (Springer US, 2021).

26. Bourgeois, Q. Monuments on the Horizon: The Formation of the Barrow Landscape throughout the 3rd and 2nd Millennium BC. (2013).

27. Beckerman, S. M. Corded ware coastal communities: Using ceramic analysis to reconstruct third millennium BC societies in the Netherlands. (Sidestone Press, 2015).

28. Fokkens, H., Steffens, B. J. W. & van As, S. F. . Farmers, fishers, fowlers, hunters: knowledge generated by development-led archaeology about the Late Neolithic, the Early Bronze Age and the start of the Middle Bronze Age (2850–1500 cal BC) in the Netherlands. Ned. Archeol. Rapp. 53, 978–990 (2016).

29. Kroon, E. J. Serial learners: interactions between Funnel Beaker West and Corded Ware communities in the Netherlands during the third millennium BCE from the perspective of ceramic technology. (Sidestone Press, 2024).

30. Furholt, M. Re-integrating Archaeology: A Contribution to aDNA Studies and the Migration Discourse on the 3rd Millennium BC in Europe. Proc. Prehist. Soc. 85, 115–129 (2019).

31. Toussaint, M. Les sépultures néolithiques du Bassin Mosan Wallon et leurs rélations avec les bassins de la Seine et du Rhin. Arch. Mosellana 7, 9–51 (2007).

32. Lanting, J. N. De NO-Nederlandse/NW-Duitse Klokbekergroep: Culturele achtergrond, typologie van het aardewerk, datering, verspreiding en grafritueel. Palaeohistoria 49–50, 11–326 (2008).

33. Cauwe, N., Vander Linden, M. & Vanmontfort, B. The Middle and Late Neolithic. Anthropol. Praehist. 112, 77–89 (2001).

34. Wentink, K. Stereotype. The role of grave sets in Corded Ware and Bell Beaker funerary practices. (Leiden: Sidestone Press, 2020).

35. Bourgeois, Q. Monuments on the Horizon: The Formation of the Barrow Landscape throughout the 3rd and 2nd Millennium BC. (2013).

36. Dyselinck, T. et al. Gent-Hogeweg, Vlakdekkende opgraving. Mariakerke. (BAAC-rapport, A-11.0045, 2013).

37. Vander Linden, M. What linked the Bell Beakers in third millennium BC Europe? Antiquity 81, 343–352 (2007).

38. Posth, C. et al. Palaeogenomics of Upper Palaeolithic to Neolithic European hunter-gatherers. Nature 615, 117–126 (2023).

39. Immel, A. et al. Genome-wide study of a Neolithic Wartberg grave community reveals distinct HLA variation and hunter-gatherer ancestry. *Commun*. Biol. 4, (2021).

40. Veselka, B., et al. Assembling ancestors : the manipulation of Neolithic and Gallo-Roman skeletal remains at Pommerœul , Belgium. 98, 1576– 1591 (2024).

41. Patterson, N. et al. Large-scale migration into Britain during the Middle to Late Bronze Age. Nature 601, 588–594 (2021).

42. ten Anscher, T. J., Knippenberg, S., van der Linde, C. M., Roessingh, W. & Willemse, N. W. Doorbraken aan de Rijn. Een Swifterbant-gehucht, een Hazendonk-nederzetting en erven en gravenuit de bronstijd in Medel- De Roeskamp. in (RAAP-rapport 6519 / Archol rapport 742 / ADC rappo, 2023).

43. Arzelier, A., et al. Neolithic genomic data from southern France showcase intensified interactions with hunter-gatherer communities. iScience 25, (2022).

44. Seguin-Orlando, A. et al. Heterogeneous Hunter-Gatherer and Steppe- Related Ancestries in Late Neolithic and Bell Beaker Genomes from Present-Day France. Curr. Biol. 1–12 (2021). doi:10.1016/j.cub.2020.12.015

45. Rivollat, M. et al. Ancient genome-wide DNA from France highlights the complexity of interactions between Mesolithic hunter-gatherers and Neolithic farmers. Sci. Adv. 6, 1–17 (2020).

46. Allentoft, M. E. et al. Population genomics of post-glacial western Eurasia. Nature 625, 301–311 (2024).

47. Brace, S. et al. Ancient genomes indicate population replacement in Early Neolithic Britain. *Nat*. Ecol. Evol. 3, 765–771 (2019).

48. González-Fortes, G. et al. Paleogenomic Evidence for Multi-generational Mixing between Neolithic Farmers and Mesolithic Hunter-Gatherers in the Lower Danube Basin. Curr. Biol. 27, 1801–1810 (2017).

49. Yu, H. et al. Genomic and dietary discontinuities during the Mesolithic and Neolithic in Sicily. iScience 25, (2022).

50. Günther, T. et al. Population genomics of Mesolithic Scandinavia : Investigating early postglacial migration routes and high-latitude adaptation. PLoS Biol. 1–22 (2018).

51. Brunel, S. et al. Ancient genomes from present-day France unveil 7 , 000 years of its demographic history. PNAS 1–8 (2020). doi:10.1073/pnas.1918034117

52. Simões, L. G. et al. Genomic ancestry and social dynamics of the last hunter- gatherers of Atlantic France. Proc. Natl. Acad. Sci. 121, e2310545121 (2024).

53. Olalde, I. et al. The genomic history of the Iberian Peninsula over the past 8000 years. Science 363, 1230–1234 (2019).

54. Olalde, I. et al. Derived immune and ancestral pigmentation alleles in a 7,000-year-old Mesolithic European. Nature 507, 225–228 (2014).

55. Fu, Q. et al. The genetic history of Ice Age Europe. Nature 534, 200–205 (2016).

56. Villalba-Mouco, V. et al. Survival of Late Pleistocene Hunter-Gatherer Ancestry in the Iberian Peninsula. Curr. Biol. 29, 1–9 (2019).

57. Mathieson, I. et al. The genomic history of southeastern Europe. Nature 555, 197–203 (2018).

58. Ringbauer, H. et al. Accurate detection of identity-by-descent segments in human ancient DNA. Nat. Genet. 56, 143–151 (2023).

59. Lazaridis, I. et al. The genetic origin of the Indo-Europeans. Nature (2025). doi:10.1038/s41586-024-08531-5

60. Linderholm, A. et al. Corded Ware cultural complexity uncovered using genomic and isotopic analysis from south-eastern Poland. Sci. Rep. 10, 1– 13 (2020).

61. Heyd, V. Yamnaya, Corded Wares, and Bell Beakers on the move. in Yamnaya Interactions. Proceedings of the International Workshop held in Helsinki, 25–26 April 2019. The Yamnaya Impact on Prehistoric Europe (eds. Heyd, V., Kulcsár, G. & Preda-Bălănică, B.) 383–414 (Archaeolingua Alapítvány, 2021).

62. Jeunesse, C. The dogma of the Iberian origin of the Bell Beaker: attempting its deconstruction. J. Neolit. Archaeol. 16, 158–166 (2015).

63. Price, T. D., Knipper, C., Grupe, G. & Smrcka, V. Strontium Isotopes and Prehistoric Human Migration: The Bell Beaker Period in Central Europe. Eur. J. Archaeol. 7, 9–40 (2004).

64. Massy, K. et al. Patterns of Transformation from the Final Neolithic to the Early Bronze Age: A Case Study from the Lech Valley South of Augsburg. in Appropriating innovations: entangled knowledge in Eurasia, 5000-1500 BCE (eds. Maran, J. & Stockhammer, P.) 241–261 (Oxbow Books, 2017).

65. Evans, J. A., Chenery, C. A. & Montgomery, J. A summary of strontium and oxygen isotope variation in archaeological human tooth enamel excavated from Britain. J. Anal. At. Spectrom. 27, 754–764 (2012).

66. van der Sloot, P. et al. Le Mésolithique et le Néolithique du site Saint-Lambert à Liège dans leur contexte chronologique, géologique et environnemental. Synthèse des données et acquis récents. Notae Praehistoricae 23, 79–104 (2003).

67. Zvelebil, M. The Social Context of the Agricultural Transition in Europe. in Archaeogenetics: DNA and the population prehistory of Europ (eds. Renfrew, C. & Boyle, K.) 57–79 (McDonald Institute monographs, 2000).

68. Armit, I. & Reich, D. The return of the Beaker folk? Rethinking migration and population change in British prehistory. Antiquity 95, 1464–1477 (2021).

69. Colledge, S., Conolly, J., Crema, E. & Shennan, S. Neolithic population crash in northwest Europe associated with agricultural crisis. Quat. Res. 92, 686–707 (2019).

70. Rascovan, N. et al. Emergence and Spread of Basal Lineages of Yersinia pestis during the Neolithic Decline. Cell 176, 295–305.e10 (2019).

71. Kristiansen, K. The Decline of the Neolithic and the Rise of Bronze Age Society. in The Oxford Handbook of Neolithic Europe (eds. Fowler, C., Harding, J. & Hofmann, D.) 1093–1118 (Oxford University Press, 2015). doi:10.1093/oxfordhb/9780199545841.013.057

72. Woodward, A. Beaker age bracers in England: sources, function and use. Antiquity 80/309, 530–543 (2006).

73. Parker Pearson, M., et al. Beaker people in Britain: migration, mobility and diet. Antiquity 90, 620–637 (2016).

74. Adler, C. J., Haak, W., Donlon, D. & Cooper, A. Survival and recovery of DNA from ancient teeth and bones. J. Archaeol. Sci. 38, 956–964 (2011).

75. Pinhasi, R., Fernandes, D. M., Sirak, K. & Cheronet, O. Isolating the human cochlea to generate bone powder for ancient DNA analysis. Nat. Protoc. 14, 1194–1205 (2019).

76. Rohland, N., Glocke, I., Aximu-Petri, A. & Meyer, M. Extraction of highly degraded DNA from ancient bones, teeth and sediments for high-throughput sequencing. Nat. Protoc. 13, 2447–2461 (2018).

77. Dabney, J. et al. Complete mitochondrial genome sequence of a Middle Pleistocene cave bear reconstructed from ultrashort DNA fragments. Proc. Natl. Acad. Sci. U. S. A. 110, 15758–63 (2013).

78. Korlević, P. et al. Reducing microbial and human contamination in dna extractions from ancient bones and teeth. Biotechniques 59, 87–93 (2015).

79. Dulias, K. et al. Ancient DNA at the edge of the world: Continental immigration and the persistence of Neolithic male lineages in Bronze Age Orkney. Proc. Natl. Acad. Sci. U. S. A. 119, 1–10 (2022).

80. Briggs, A. W. et al. Removal of deaminated cytosines and detection of in vivo methylation in ancient DNA. Nucleic Acids Res. 38, e87 (2009).

81. Rohland, N., Harney, E., Mallick, S., Nordenfelt, S. & Reich, D. Partial uracil – DNA – glycosylase treatment for screening of ancient DNA. Philos. Trans. R. Soc. London B 370, (2015).

82. Gansauge, M. T., Aximu-Petri, A., Nagel, S. & Meyer, M. Manual and automated preparation of single-stranded DNA libraries for the sequencing of DNA from ancient biological remains and other sources of highly degraded DNA. Nat. Protoc. 15, 2279–2300 (2020).

83. Fu, Q. et al. An early modern human from Romania with a recent Neanderthal ancestor. Nature 524, 216–219 (2015).

84. Rohland, N. et al. Three assays for in-solution enrichment of ancient human DNA at more than a million SNPs. Genome Res. 32, (2022).

85. Kircher, M., Sawyer, S. & Meyer, M. Double indexing overcomes inaccuracies in multiplex sequencing on the Illumina platform. Nucleic Acids Res. 40, 1–8 (2012).

86. Li, H. & Durbin, R. Fast and accurate short read alignment with Burrows– Wheeler transform. Bioinformatics 25, 1754–1760 (2009).

87. Fu, Q. et al. A revised timescale for human evolution based on ancient mitochondrial genomes. Curr. Biol. 23, 553–9 (2013).

88. Korneliussen, T. S., Albrechtsen, A. & Nielsen, R. ANGSD : Analysis of Next Generation Sequencing Data. BMC Bioinformatics 15, 1–13 (2014).

89. Patterson, N. et al. Ancient admixture in human history. Genetics 192, 1065–93 (2012).

90. Biagini, S. A. et al. People from Ibiza: an unexpected isolate in the Western Mediterranean. Eur. J. Hum. Genet. 941–951 (2019). doi:10.1038/s41431-019-0361-1

91. Lazaridis, I. et al. The genetic history of the Southern Arc: A bridge between West Asia and Europe. Science 377, (2022).

92. Fowler, C. et al. A high-resolution picture of kinship practices in an Early Neolithic tomb. Nature 601, 584–587 (2021).

93. Ringbauer, H., Novembre, J. & Steinrücken, M. Parental relatedness through time revealed by runs of homozygosity in ancient DNA. Nat. Commun. 12, 1–11 (2021).

94. Patterson, N., Price, A. L. & Reich, D. Population structure and eigenanalysis. PLoS Genet. 2, e190 (2006).

