## Supplementary Information for "Long-term hunter-gatherer continuity in the Rhine-Meuse region was disrupted by local formation of expansive Bell Beaker groups"

#### Table of contents

SI 1. Archaeological overview of the Rhine-Meuse region 8500-1700 BCE

SI 2. Archaeological context information about the newly reported and published individuals from the Rhine-Meuse area

SI 3 Analytical details Sr-O-C isotope analysis

SI 4. *qpAdm* modeling of ancestry proportions

SI 5. IBD sharing analysis

### SI 1. Archaeological overview of the Rhine-Meuse region 8500-1700 BCE

#### A mosaic of 'cultures': an archaeological survey 8500-1700 BCE of the Low Countries

Harry Fokkens, Quentin Bourgeois, Eveline Altena, Luc Amkreutz

#### Contents

1 Mesolithic 8500-5000 BCE ... 4

1.1 Short summary of the sample context for Table 1; for the extended version we refer to SI 2 ... 4

1.2 Cultural dynamics in the Mesolithic 8500-5000 BCE ... 6

2 The first farmers (5000-4000 BCE) ... 6

2.1 Summary of the archaeological context for the individuals in Table 2; for the extended version we refer to SI 2 ... 7

2.2 Cultural dynamics in the Early Neolithic 5000-4000 BCE ... 8

3 Middle Neolithic farmers (4000-3500 BCE) ... 9

3.1 Summary of the context information for the individuals presented in Table 3; for the extended version we refer to SI 2 ... 9

3.2 Cultural dynamics in the Middle Neolithic 4000-3500 BCE ... 10

4 Middle – Late Neolithic farmers (3500-3000 BCE) ... 11

4.1 Summary of the context information for the individuals presented in Table 4; for the extended version we refer to SI 2 ... 12

4.2 Cultural dynamics at the end of the Middle Neolithic 3500-3000 BCE ... 13

5 Late Neolithic A 3000-2500 BCE. ... 15

5.1 Summary of the archaeological context of the samples in table 5; for the extended version we refer to SI 2 ... 15

5.2 Cultural dynamics in the Late Neolithic A: 3000-2500 BCE ... 17

6 Late Neolithic and Early Bronze Age: AOO, Bell Beaker and Barbed Wire Beaker ... 19

6.1 Summary of the archaeological context of the samples in table 6; for the extended version we refer to SI 2 ... 20

6.2 Cultural dynamics in the Late Neolithic B and Early Bronze Age: 2500-1700 BCE: Bell Beaker and Barbed Wire Beaker ... 20

In this overview we sketch the broad cultural developments in the Rhine-Meuse delta in a temporal framework. The goal of this supplement is to provide an archaeological background for the interpretation of the DNA results. Since the Neolithic is characterized by regionally different manifestations of similar archaeological cultures, we discuss how the data from the 'Low Countries' fit within this complex mosaic.

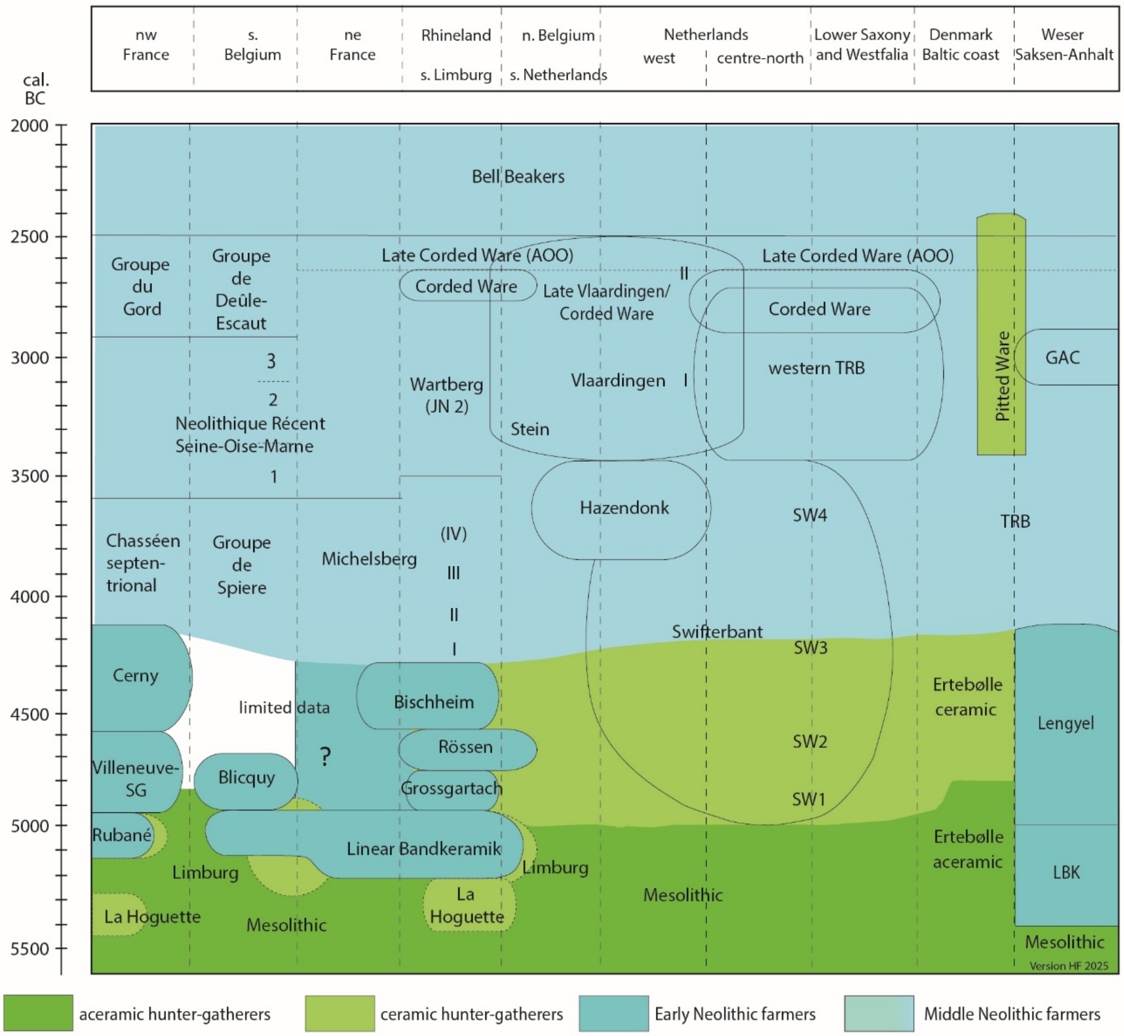

*Figure 1 Schematic overview of archaeological formations in the Early and Late Neolithic in northwest Europe (adapted and updated from Louwe Kooijmans 2006)*[1].

Figure 1 presents the mosaic of cultures that archaeologists recognize against the background of their economic bases and the concept of Neolithization, emphasizing that this was a fluid process in most areas, and not at all a synchronous one. It tries to visualize transitions from hunter-gatherer economies to farming economies in different regions. Early Neolithic farmers are separated from Middle Neolithic farmers in this model in order to indicate the difference between Early Linearbandkeramik (LBK) immigrant farmers and their descendants and a much more diffuse trajectory of adopting farming elements by 'indigenous' hunter-gatherer communities. This does not imply that Middle Neolithic farmers 'slowly' adopted farming, but that their economy was regionally specific, often relying to some extent on hunting, gathering and fishing. We argue that in wetlands and in river valleys of the Rhine and Meuse the nature of their settlement environment contributed to this different adoption of 'the' Neolithic than on the more eastern sandy uplands and southern loess zones.

### 1 Mesolithic (8500-5000 BCE)

| Genetic ID | Archaeological ID | Full Date | Locality |
|---|---|---|---|
| AAT001 | AA3 | 9160-8623 calBCE (9500±75 BP, OxA-4917) | Abri des Autours (BE) |
| I7015 | BELG_6598 | 9100-8400 BCE | ClaminForge (BE) |
| MPR001.AG | MPR-1 | 8731-8294 calBCE (9270±90 BP, OxA-5042) | Malonne Petit Ri (BE) |
| I7018 | BELG_7764 | 8547-8293 calBCE (9190±45 BP, PSUAMS-7873) | Grotte_Rousseau (BE) |
| DOG002 | U 2014/12.4; A10-007_V003_M006 | 8421-8238 calBCE (9091±37 BP, MAMS-34582) | Doggerland, West of Brown Bank |
| I7010 | BELG_265 | 8000-5500 BCE | Grotte de la faille du burin (BE) |
| DOG001 | A10-007_V002_M003 | 7730-7586 calBCE (8627±35 BP, MAMS-48201) | Doggerland, Eurogeul |
| DOG007 | U 2014/12.3; A10-007_V001_M001 | 7576-7201 calBCE (8370±50 BP, GrA-11642) | Doggerland |
| SPI001 | 2212/2:1 | 5558-5373 calBCE (6510±40 BP, Poz-103001) | Spiekeroog (GE) |
| I13024 | V28.578 Trijntje | 5802-5626 calBCE (6820±50 BP, GrA-9804), corrected to 5500-5400/5300 calBCE (Dreshay *et al.* 2023) | Hardinxveld-Polderweg (NL) |

*Table 1 Summary of samples used for analysis from the period 8500-5000 BCE*

#### 1.1 Summary of the archaeological context for the individuals in Table 1; for the extended version we refer to SI 2

The Belgian samples are all from (open) cave contexts. Sample I7015 is from a small cave site on a tributary of the Sambre (Claminforge), excavated by Michel Toussaint in 1995[2]. Radiocarbon dates on two individuals from this collection of burials place them well into the Mesolithic. Sample I7018 is from a cave site for which we lack reliable archaeological information (Grotte Rousseau). However, five radiocarbon dates are available for this site. The one that we generated directly on the sample we analyzed for DNA is the only one dating to the Mesolithic, and the genetic profile is typical for Mesolithic western European hunter-gatherers.

All the other dates are from the Middle and late Neolithic and the related individuals have substantial proportions of ancestry from Anatolian Neolithic farmers. Le Grotte du faille du Burin is a small cave in which the remains at least 12 individuals were found. Four dates are available, all falling in the Mesolithic [3]. It is not clear whether I7010 was one of those, but the genetic profile is typical for Mesolithic western European hunter-gatherers

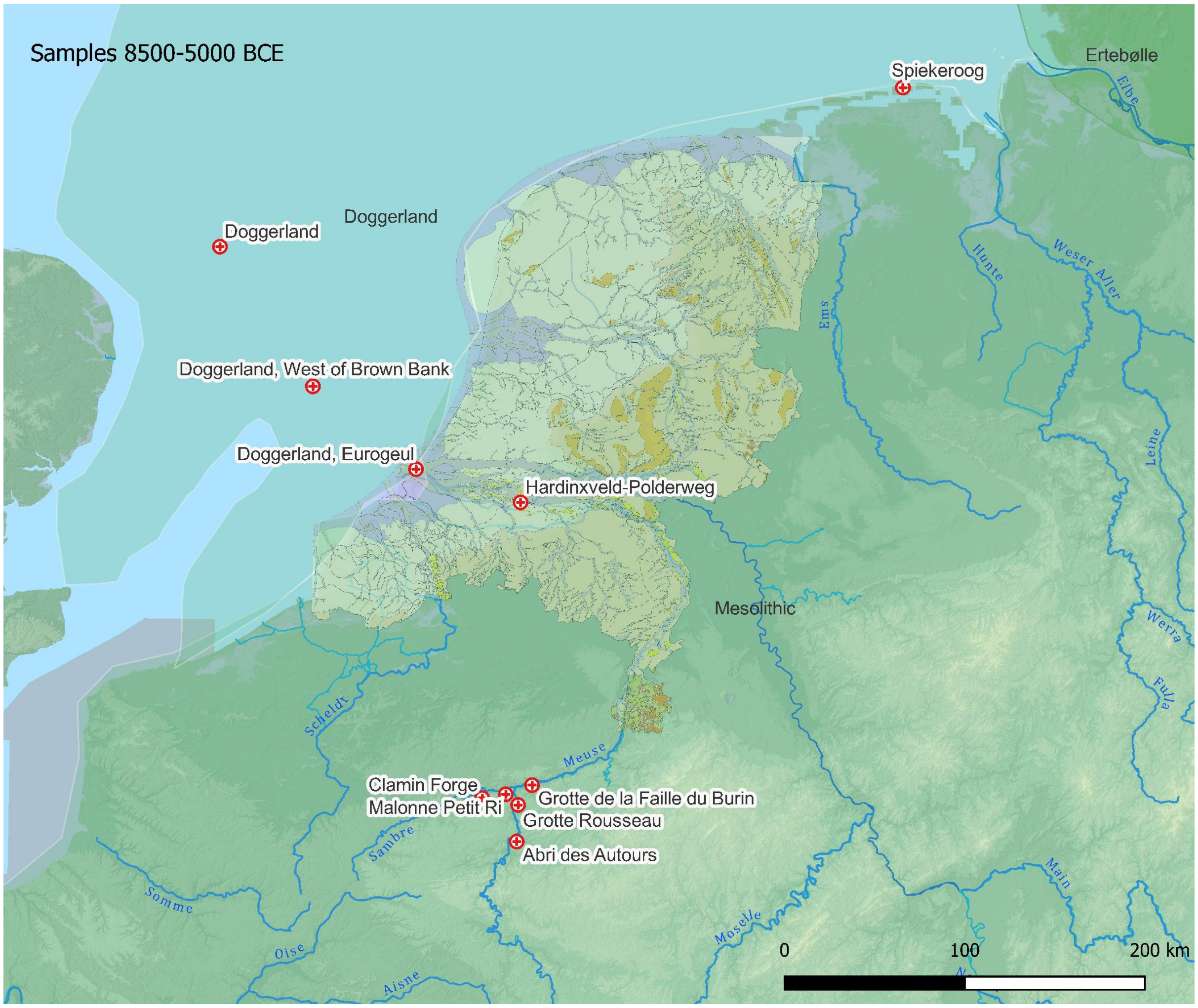

*Figure 2 Schematic distribution of cultural spheres and the geographic locations of samples 8500-5000 BCE. The paleogeographic reconstruction of the Netherlands in 9000 BCE is after Vos et al.*[4]*; the extent of Doggerland is after Coles*[5]*. The elevation map is from* https://www.mapsforeurope.org/datasets/euro-dem *(the grey area in the English Channel is uncharted).*

The samples from Doggerland, Abri des Autours and Malonne Petit Ri have been published by Posth *et al.* 2023[4]. The Abri des Autours is a cave site on the right bank of the Meuse near the town of Dinant. It contains burial structures from the Mesolithic and the Middle Neolithic [4,5]. The Grotte du Petit Ri is located in Malonne, near Namur (Belgium). The remains of several individuals, lithics and faunal remains were found [5,6]. The Doggerland individuals are from different locations in the North Sea, mostly discovered by fishermen or in sand supplies from the North Sea [7,8]. The sample from Spiekeroog is a beach find from the northern coast of the island. The Hardinxveld-Polderweg sample I13024 is from a well-documented grave from a Mesolithic context on a (now) submerged river dune [9,10].

##### 1.2 Cultural dynamics in the Mesolithic 8500-5000 BCE

Before early farmers of the LBK reached northwestern Europe, the area —and especially the river deltas and seaside locations— was occupied by Late Mesolithic hunter-gatherer communities with slightly different regional traditions. Especially the vast area of North Sea Doggerland is thought to have been densely occupied[5,11,12]. The Rhine-Meuse delta, bounded in the west by the Scheldt and in the north the Vecht, once formed the eastern fringe of Doggerland. During the Holocene Doggerland gradually drowned due to rising sea levels and changed in an 'archipelago' in front of the Thames and Rhine-Meuse estuaries, extending far into the present North Sea[13]. After the Storrega landslide tsunami around 6200 BCE, this process of drowning accelerated[12,13]. Around 5500 BCE Doggerland had completely drowned [13], and the sea had advanced to about the present-day coastline of the southern North Sea (*Figure 2*). Its inhabitants likely migrated toward these new coastlines and connected inland deltas.

In Denmark and on the Baltic coast, the Late Mesolithic Ertebølle hunter-gatherer-fishers formed a '*coastal adaptation with a focus on marine resources, especially fish*'[14], in particular cod. In northern and eastern Jutland, the Ertebølle Culture is best known from its (seasonal) shell middens with oyster shells (*kjøkkenmøddinger*):[15], but many more sites are known without shell middens, both on the coast and inland [14,16]. In the Dutch coastal areas and deltas the hunter-gatherers of the Swifterbant culture practiced a similar economy until c. 4200 BCE. From that period onwards they also practiced arable farming and husbandry while continuing to exploit freshwater resources and living on raised areas in, or bordering the wetlands [17,18].

Hunter-gatherers started to produce Swifterbant pottery from between 5100 and 4800 BCE (Dreshaj *et al.* 2023), at approximately the same time (4800 BCE) as in Belgium [19] and Ertebølle contexts [20], demonstrating that these hunter-gatherer-fisher communities were in close contact with each other. Farming was not adopted until a few hundred years later[21].

In some respects, the Mesolithic occupation in Belgium resembles the Dutch situation, with characteristics very similar to the Swifterbant hunter-gatherers along the Scheldt and Meuse River valleys[22].

#### 2 The first farmers (5000-4000 BCE)

| Genetic ID | Archaeological ID | Full Date | Locality |
|---|---|---|---|
| I12091 | NGKL10-ID1 | 4400-3900 BCE | Nieuwegein-het Klooster (NL) |
| I12093 | NGKL10-ID5 | 4200-4000 BCE | Nieuwegein-het Klooster (NL) |
| I12094 | NGKL10-ID6 | 4200-4000 BCE | Nieuwegein-het Klooster (NL) |
| I17968 | NGKL10-ID4 | 4342-4171 calBCE (5410±30 BP, ICA-6954) | Nieuwegein-het Klooster (NL) |
| | | 4200-3800 BCE | |
| I33738 | skelet I | 4227-3808 calBCE (5190±50 BP, UtC-1961 | Zoelen_de Beldert (NL) |
| I33739 | skelet II | 4200-3800 BCE | Zoelen_de Beldert (NL) |
| SWA001 | Skelet II | 4180-4030 BCE | Swifterbant S2 (NL) |
| SWA002 | Skelet III | 4180-4030 BCE | Swifterbant S2 (NL) |
| SWA004 | Skelet IV | 4180-4030 BCE | Swifterbant S2 (NL) |

*Table 2 Summary of the samples used for analysis from the period 5000-4000 BCE.*

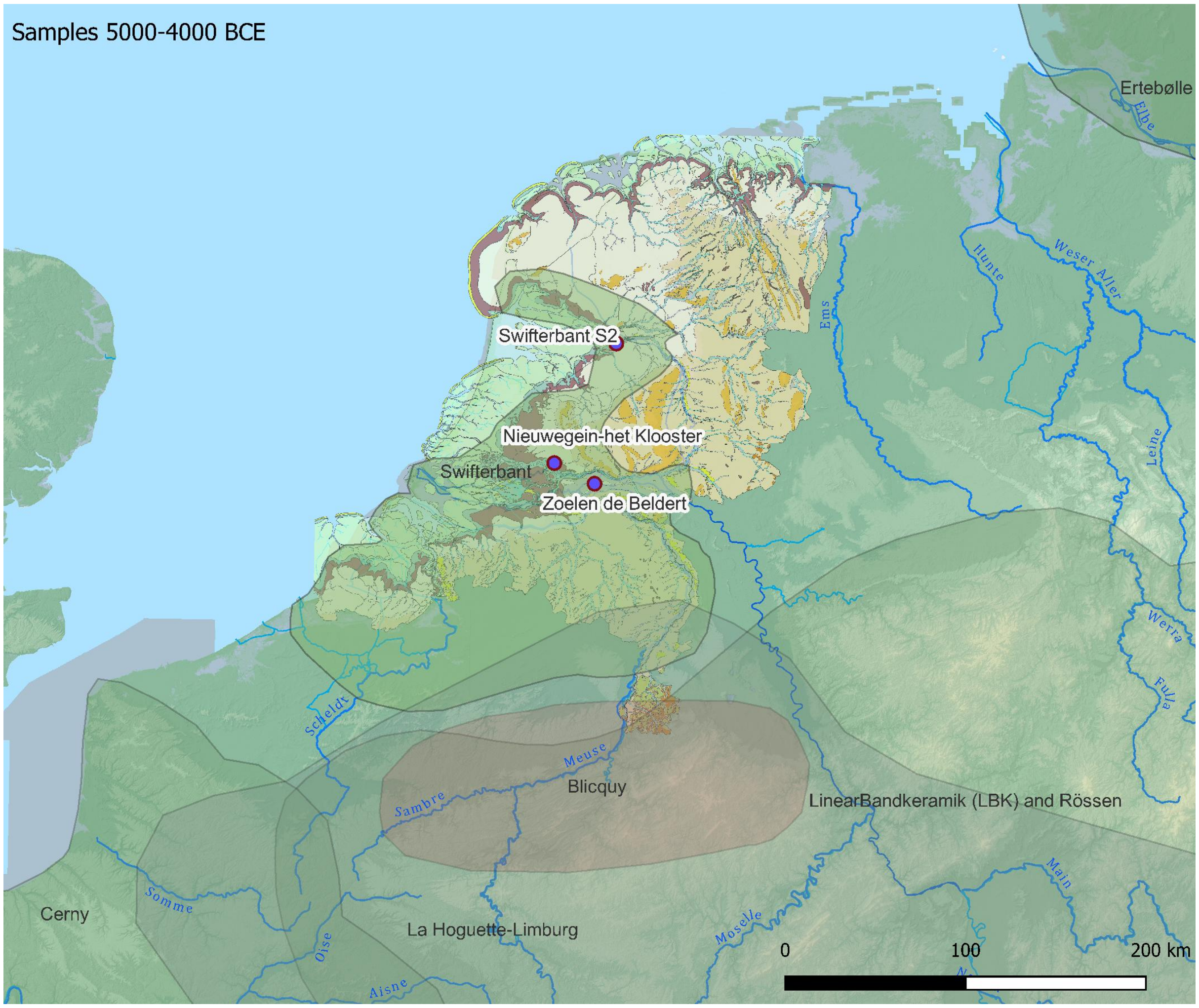

*Figure 3 Schematic distribution of cultural spheres and the geographic locations of samples from 5000-4000 BCE. The paleogeographic reconstruction of the Netherlands 5500 BCE is after Vos et al.*[4]. *The elevation map is from* *https://www.mapsforeurope.org/datasets/euro-dem* *(the grey area in the English Channel is uncharted).*

#### 2.1 Summary of the archaeological context for the individuals in Table 2; for the extended version we refer to SI 2

Three individuals were sampled from one of the eponymous sites at Swifterbant (site S2). These are located on a small river system. Sites S3 and S4 are within meters distance from one another, while S2 is located at some 500 m distance. The site remains make clear that the occupants practiced hunting and animal husbandry, gathering, and cultivation. High resolution Bayesian modelling showed that the site is most likely dated between 4180 and 4030 BCE [21], which places it in a later phase of the Swifterbant culture (SW2).

The Nieuwegein site is unique in the Netherlands because it is one of the few Early Swifterbant sites known. At Nieuwegein several graves were found, one of which was of a baby and her mother (I12093 and I12094). Both lacked Early European farmer genetic ancestry, but the other two sampled individuals harbor a mixture of a major ancestry component associated with Mesolithic hunter-gatherers and a minor component associated with Anatolian farmers. The same is true for the Late Swifterbant Zoelen-de Beldert individuals, who have a contemporary date.

The individuals analyzed from Zoelen-de Beldert are from a rescue excavation of a pit or grave that contained the remains of three people. Sample I33739 is from skeleton II, a woman lying on the bottom of the pit. On top of that was the secondary deposition of the remains from another woman (sample I33738)[23]. Finally, at the top of the pit a poorly preserved skeleton was found of a child, which did not yield sufficient genetic data.

#### 2.2 Cultural dynamics in the Early Neolithic 5000-4000 BCE

The first Linearbandkeramik (LBK) farmers arrived on the loess plateaus of southern Limburg and eastern Belgium west of the Rhine around 5300 BCE[12,24] (*Figure 3*). From the archaeological evidence it is clear that, while their economy was fully based on farming, it was strictly bound to the fertile loess soils. The sandy uplands north of this zone were much less fertile, were exhausted more quickly, and were probably less suitable for the cultivation of emmer (*Triticum monococcum*) and einkorn (*Triticum dicoccum*) wheat, which were staples for LBK farmers[24–27]. From settlement evidence it is clear that LBK farmers also raised sheep, goats, pigs and cattle to supplement their diet[24]. The descendants of these cultures (*Figure 3*), like the Grossgartach, Rössen, and Bischheim farmers in the east, and the Villeneuve-Saint Germain, Cerny and Groupe de Blicquy farmers in the west, basically grew the same crops and kept similar livestock, although Cerny and Rössen farmers also added bread wheat (*Triticum aestivum*) to their diet[24,28]. The early farming traditions in general kept to the loess belts of eastern and southern Belgium, northern France, the Paris Basin, and the Rhineland[e.g. 29]. There is, however, still a lot of debate about the role of La Hoguette and Limburg pottery. This non-LBK ware occurs also outside the loess areas, and does not seem to have a clear connection with LBK pottery even though it occurs on several LBK sites[30]. Constantin *et al*[31]. therefore think this style is a development from within LBK, while others think it might (also) be related to hunter-gatherer communities [30,32]. From the archaeological evidence it is not clear what happened in Southern Belgium and Northeastern France (*cf. Figure 1*). In the region situated between the distribution of the Rubané/LBK and Blicquy cultures, the archaeological record is scarce in Southern Belgium and part of Northeastern France, with no Mesolithic and little Neolithic material[33–36].

Further east, in central Europe, the settlers of the LBK were followed by the early farmers of the Lengyel culture who had their origins probably in Poland [37–39]. Lengyel farmers were still bound to the loess soils and practiced agriculture in a similar way to their LBK predecessors.

While the LBK was thriving across large parts of northern Europe, from France and Belgium in the west to Poland and Ukraine in the east, the river deltas and seashores of the southern North Sea and the Baltic were inhabited by 'indigenous' hunter-gatherer-fisher communities. Even though these communities adopted aspects of the Neolithic way of life —first pottery making, and later also elements of farming— it would not be correct to characterize them as (full) farmers just because they practiced farming as well. For this phase of the Neolithic in the wetlands of the Rhine-Meuse delta Louwe Kooijmans[40] introduced the term "extended broad spectrum economies". In his later models he adopted the term 'semi-agrarian' especially for the communities that in addition to farming, still had hunting and fishing as central parts of their economies[12]. As this appears to have been a constant feature of all communities living in wetlands throughout the Middle Neolithic, even until the Late Neolithic and the Bronze Age[41].

### 3 Middle Neolithic farmers (4000-3500 BCE)

| Genetic ID | Archaeological ID | Full Date | Locality |
|---|---|---|---|
| I1565 | Bla8+Bla9+Bla11+Bla24+Bla26(x)+Bla45 (Excavation 2004) \| Bla8+Bla9+Bla11+Bla24+Bla26(x)+Bla45 (Excavation 2014) | 3725-3655 calBCE (4965±15 BP) | Blätterhöhle Cave (GE) |
| I1563 | Bla5+Bla7+Bal13+Bla26(o)+Bla30+Bla54 (Excavation 2004) \| Bla5+Bla7+Bal13+Bla26(o)+Bla30+Bla54 (Excavation 2014) | 3626-3378 calBCE (4726±17 BP) [R_combine: (4580±30, KIA-28844, Bla5); (4860±30, KIA-45011, Bla7); (4730±25, KIA-45010, Bla13)] | Blätterhöhle Cave (GE) |
| I1593 | Bla16+Bla27+Bla59 (Excavation 2004) \| Bla16+Bla27+Bla59 (Excavation 2014) | 3644-3528 calBCE (4810±23 BP) | Blätterhöhle Cave (GE) |
| I35542 | grave 4_S1137 | 3800-3600 BCE | Tiel_Medel (NL) |
| I35543 | grave 1_S1147 | 3800-3600 BCE | Tiel_Medel (NL) |
| I35544 | grave 3_S2386 | 3800-3600 BCE | Tiel_Medel (NL) |
| I38121.TW | 04HP_M120 | 3800-3600 BCE | Schipluiden-Harnaschpolder (NL) |
| I38447.TW | Grave 5, Ind. 6; 04HP_M113 | 3630-3500 calBCE (5170±40 BP, GrA-26672) | Schipluiden-Harnaschpolder (NL) |
| I38448.TW | Grave 6, Ind. 7; 04HP_M115 | 3550-3490 calBCE (5005±40 BP, GrA-26650) | Schipluiden-Harnaschpolder (NL) |

*Table 3 Summary of samples used for analysis from the period 4000-3500 BCE*

#### 3.1 Summary of the context information for the individuals presented in Table 3; for the extended version we refer to SI 2

The individuals analyzed from Blätterhöhle Cave are from the German Mittelgebirge, at a site with both Mesolithic and Neolithic graves[42,43]. The samples analyzed for DNA came from a scatter of nine bones in the cave all of which yielded DNA and which were treated as independent samples in the study that first reported mitochondrial DNA, but for which genome-wide data showed they belonged to two distinct individuals[44].

The site of Tiel-Medel is situated on a stream ridge or crevasse of the Rhine. It has Swifterbant, but also Hazendonk period occupation and graves. A third phase at the site can be dated to the late 3rd-2nd Millennium BCE. The individuals reported here were published as Bronze Age individuals[45] but the complete lack of Steppe related ancestry in these individuals and IBD analysis suggest that an association with the Middle Neolithic Hazendonk culture is more likely (see S2. for more information).

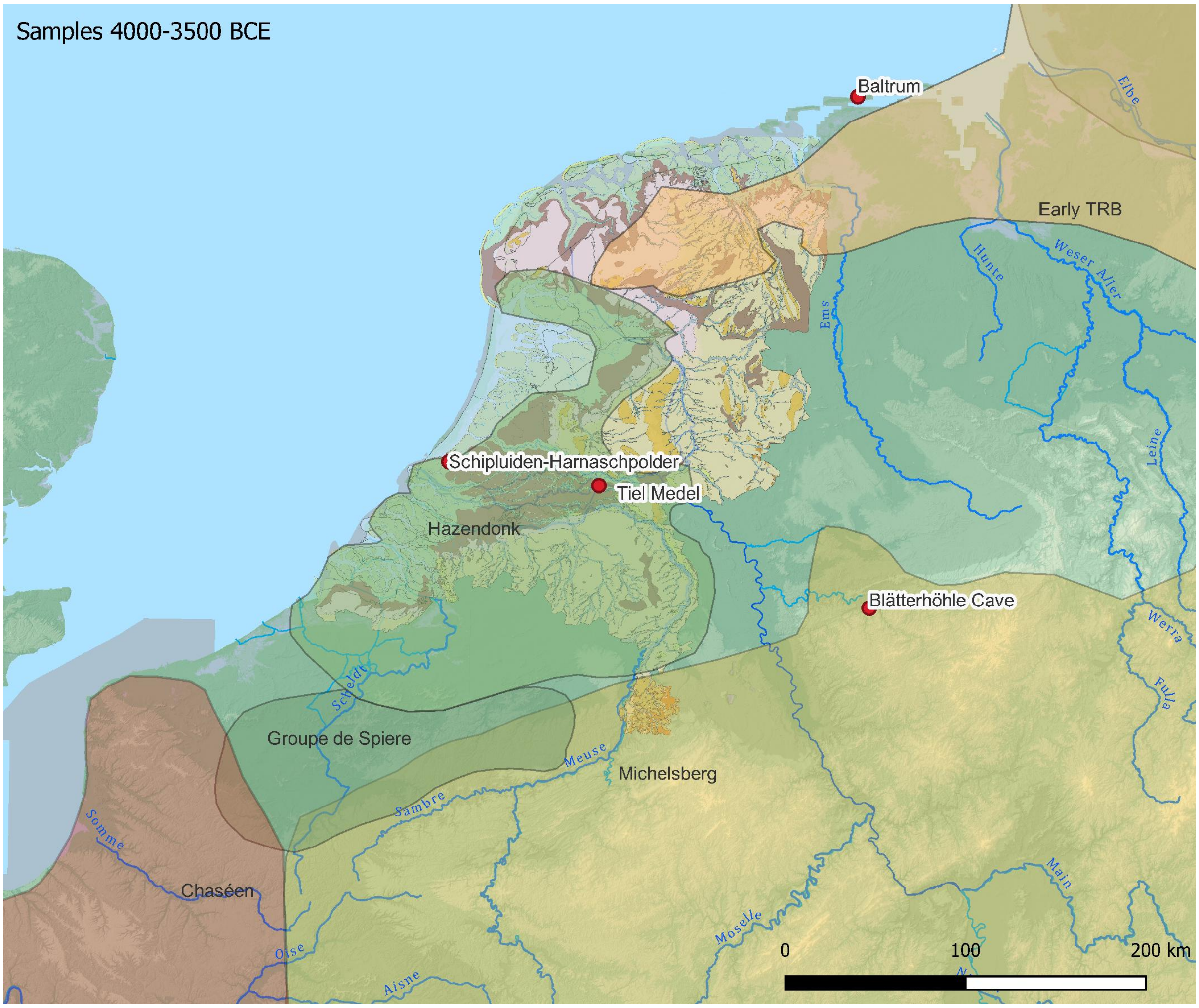

*Figure 3 Schematic distribution of cultural spheres and the geographic locations of samples from 4000-3500 BCE. The paleogeographic reconstruction of the Netherlands in 3850 BCE is after Vos et al.*[4]*. The elevation map is from [https://www.mapsforeurope.org/datasets/euro-dem](https://www.mapsforeurope.org/datasets/euro-dem) (the grey area in the English Channel is uncharted).*

The individuals from Schipluiden are from burials in a Middle Neolithic settlement with huts or houses, situated near the coast on a beach barrier[46]. The habitation on the site must have ended before 3400 BCE [47,48].

#### 3.2 Cultural dynamics in the Middle Neolithic 4000-3500 BCE

In most regions the Middle Neolithic was characterized by the first use of flint and stone axes, exploitation of flint mines, and diversification of agricultural practices and crops[24]. The Funnel Beaker Culture (TRB) successors to the Lengyel Culture, for instance, practiced swidden cultivation on the lighter sandy soils north of the loess belt from c. 4000 BCE onwards[38]. The transition from hunting to farming societies in northern Europe took place in a very short period between 4200 BCE and 3800 BCE. The first TRB pottery appears in a Late Ertebølle context around 4100 BCE without replacing other aspects of Ertebølle material. This suggests that Ertebølle hunters-fishers adopted TRB pottery, yet not their farming practices. *Figure 3* shows the distribution of cultural formations that are recognized for this period. The distribution of Early TRB sites is sketchy because only few are known. We assume that most of the forested uplands of NE Netherlands and NW Germany were uninhabited (although perhaps exploited), based on the very limited archaeological indications for this, even though the area has been investigated. Exceptions are early 'pre-Drouwen' TRB occupation in lower lying areas (Schokland-P14[49]; Westingermaar[50]).

The Rhine-Meuse delta is dominated by sites of the Late Swifterbant and Hazendonk traditions. Only the later Hazendonk 3 phase is nowadays distinguished as a local variant of the Swifterbant and Michelsberg cultures (*Figure 1*[51,52]). Its distribution is mainly restricted to the river dunes (*donken*) in the central Rhine-Meuse delta. The only individuals sampled that were associated with Hazendonk 3 pottery, are the individuals from the Schipluiden site.

The Blätterhöhle cave is situated within the northern distribution of the Michelsberg Culture. That is contemporary with similar regional cultures such as the Groupe de Spiere in Belgium[53] and the Chasséen in northern France[54] (see Bakels[24] for an extensive overview of economy and settlement). Even though these groups all practiced farming, their economy was not a full farming economy across the region, with the Groupe de Spiere sites being characterized by wetland locations, and by a mix of farming, hunting, fishing, and gathering[55,56].

#### 4 Middle – Late Neolithic farmers (3500-3000 BCE)

| Genetic ID | Archaeological ID | Full Date | Locality |
|---|---|---|---|
| I1594 | Bla28 (Excavation 2004) | 3338-3024 calBCE (4465±30 BP, KIA-28846) | Blätterhöhle Cave (BE) |
| KH150613_KH180043 | NT107 | 3346-3098 calBCE (4499±26 BP, KIA-53045) | Niedertiefenbach (GE) |
| KH150614_KH150615 | NT142.1 | 3334-3025 calBCE (4462±24 BP, KIA-53046) | Niedertiefenbach (GE) |
| KH150618 | NT48 | 3336-3028 calBCE (4468±25 BP, KIA-52275) | Niedertiefenbach (GE) |
| KH150619 | NT42 | 3333-3021 calBCE (4455±24 BP, KIA-52276) | Niedertiefenbach (GE) |
| KH150620 | NT148 | 3344-3094 calBCE (4491±25 BP, KIA-53047) | Niedertiefenbach (GE) |
| KH150621 | NT50 | 3500-2800 BCE | Niedertiefenbach (GE) |
| KH150622 | NT130 | 3264-2926 calBCE (4417±19 BP, KIA-53048) | Niedertiefenbach (GE) |
| KH150623 | NT135 | 3500-2800 BCE | Niedertiefenbach (GE) |
| KH150625 | NT83 | 3348-3096 calBCE (4497±27 BP, KIA-52277) | Niedertiefenbach (GE) |
| KH150626 | NT58 | 3500-2800 BCE | Niedertiefenbach (GE) |
| KH150627 | KI11 | 3335-3024 calBCE (4462±27 BP, KIA-52278) | Niedertiefenbach (GE) |
| KH150628 | NT150.1 | 3334-2938 calBCE (4448±26 BP, KIA-53049) | Niedertiefenbach (GE) |
| KH150629 | NT30 | 3500-2800 BCE | Niedertiefenbach (GE) |
| KH150630 | KI12 | 3487-3106 calBCE (4564±25 BP, KIA-53050) | Niedertiefenbach (GE) |
| KH150633 | KI13 | 3344-3036 calBCE (4486±29 BP, KIA-52279) | Niedertiefenbach (GE) |
| KH150635 | KI14 | 3322-2924 calBCE (4425±28 BP, KIA-52280) | Niedertiefenbach (GE) |
| KH150637 | NT110 | 3328-2926 calBCE (4432±28 BP, KIA-52281) | Niedertiefenbach (GE) |
| KH150639 | NT49 | 3500-2800 BCE | Niedertiefenbach (GE) |
| KH150640 | NT98 | 3334-3024 calBCE (4461±25 BP, KIA-53053) | Niedertiefenbach (GE) |

| | | | |
|---|---|---|---|
| KH150641 | KI15 | 3339-3028 calBCE (4473±28 BP, KIA-52282) | Niedertiefenbach (GE) |
| KH180044 | NT136.1 | 3500-2800 BCE | Niedertiefenbach (GE) |
| KH180045 | NT146 | 3500-2800 BCE | Niedertiefenbach (GE) |
| I13627 | 1 / x4; AF004 | 3334-3028 calBCE (4466±21 BP, OxA-39060) | Trou Al'Wesse, Modave (BE) |
| I18068 | T26-C | 3011-2890 calBCE (4320 ± 27 BP, RICH27887) | Pommeroeul (BE) |
| I21570 | T26-J | 3017-2906 calBCE (4278 ± 27 BP, RICH27891) | Pommeroeul (BE) |
| I7014 | BELG_273 | 3100-2900 BCE | Weris_II (BE) |

*Table 4 Summary of samples used for analysis from the period 3500-3000 BCE*

##### 4.1 Summary of the context information for the individuals presented in Table 4; for the extended version we refer to SI 2

The samples cited above are from Germany, from a cave site in the Mittelgebirge (Blätterhöhle GE), a gallery grave near Niedertiefenbach (GE), from an open air site (Pommereuil BE), a cave site (Trou Al'Wesse) and an *allée couverte* (Weris II). The two German sites are associated with the Wartberg Culture, a late variant of the Michelsberg Culture (Immel *et al.* 2021), while the three Belgian sites are probably associated with the Seine-Oise-Marne (SOM) culture, even though only Wéris II is associated with context finds[57]. The individual from Trou Al'Wesse was excavated in a cave (in the Ardennes; Miller *et al.* 2011; 2012), while the sample from Weris II originated from a megalithic grave of a type that had a wide distribution all over western Europe in this period.

The Pommereuil samples come from a composite burial that was discovered in a Gallo-Roman cemetery as the only inhumation grave[58]. Osteo-archaeological and DNA analysis, combined with intensive dating of the separate bones, revealed that it was composed of at least seven individuals, with Late Neolithic dates for the postcranial skeletal parts and a Gallo-Roman date for the cranium[58].

The Blätterhole, Niedertiefenbach and Pommereuil sites were published earlier, whereas the new samples published here are from Wéris II and Trou Al'Wesse in Belgium. The Niedertiefenbach site stood out because the 42 sampled individuals showed a relatively high (34-58%) proportion of WHG ancestry despite their relatively young date of 3300 and 3200 BCE[59].

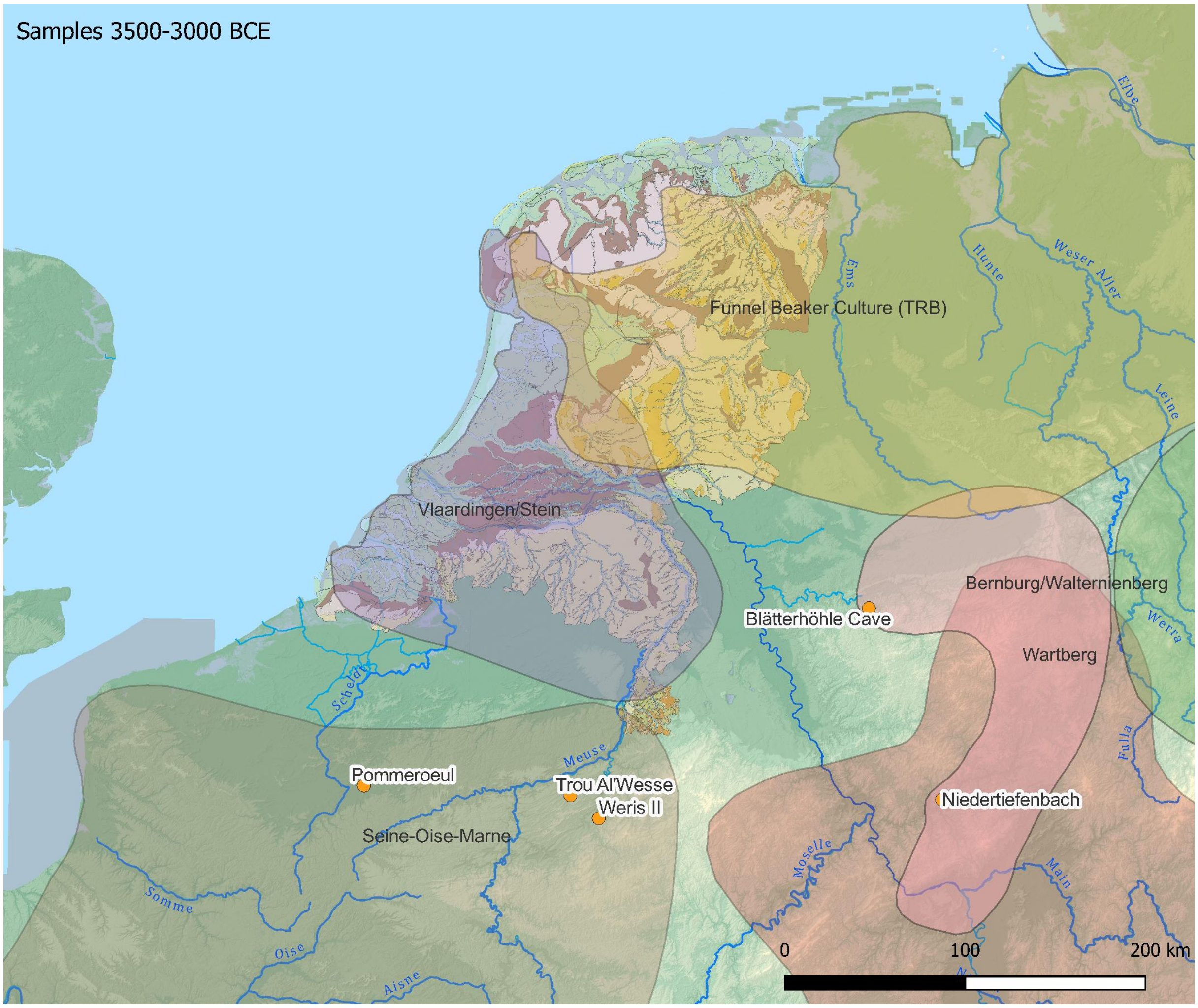

*Figure 4 Schematic distribution of cultural spheres and the geographic locations of newly analyzed and published samples 3500-3000 BCE. The paleogeographic reconstruction of the Netherlands 3850 BCE is after Vos et al.*[4]*. The elevation map is from* [https://www.mapsforeurope.org/datasets/euro-dem](https://www.mapsforeurope.org/datasets/euro-dem) *(the grey area in the English Channel is uncharted).*

##### 4.2 Cultural dynamics at the end of the Middle Neolithic 3500-3000 BCE

The Late Neolithic begins at different moments between 3500 and 3000 BCE in most regions. Several archaeological cultures have been distinguished, mainly on the basis of burial traditions, pottery styles and flint assemblages.

In the Paris Basin, in the north of France, and the adjacent Rhineland, the late 4th and early 3rd millennium are characterized by the megalithic "Seine-Oise-Marne" Culture (SOM) (*Figure 4*). Dates for SOM vary but this complex should likely be placed between 3350 and 3000 BCE[60]. This places SOM in the same temporal framework as, for instance, the Vlaardingen, Late Michelsberg, Wartberg and Funnel Beaker (TRB) cultures. From a material culture perspective these regional groups or cultures are closely related, for instance SOM, Wartberg and TRB all built megalithic structures of a kind, which were then often reused until the end of the 3rd millennium BCE. Cultural similarities are mirrored in pottery forms (not or sparsely decorated) and technology (even though these are regionally different), and in economy.

The only site in the Netherlands that has been associated with SOM is a cellar with a stone floor in the Meuse valley near Stein[61–63]. As such it is the most northern expression of a tradition of collective graves in megalithic monuments or in caves and abris that was typical for the SOM tradition.

Habitation in the deltas of the rivers Scheldt, Meuse, Rhine and Vecht is characterized by Vlaardingen settlements on beach barriers, on Pleistocene river dunes and on levees and crevasse splays but also further inland along the river Meuse and its tributaries (*Figure 5*). Their economy consisted of small-scale farming on these elevated areas, but also hunting, gathering and fishing. In the few cemeteries that have been found near these settlements the dead are buried in a crouched position in 'flat' graves without grave gifts apart from amber beads[49,64].

North and east of the river deltas, on the ice-sculpted and sand-covered landscape of the Veluwe and the northern and eastern Netherlands, the cultural situation appears to have been a little different. These 'uplands' were probably heavily forested, and we have little evidence for Neolithic habitation on the plateaus before the TRB tradition started to build their megalithic monuments around 3500 BCE[50]. In northwest Germany and the northeast Netherlands, a separate branch of the TRB developed: the western TRB[65]. There are indications that the first TRB sites developed from 3500 onwards in riverine environments, not unlike those in northwest Germany (e.g. Hüde)[49,50,66]. These sites are all located in the north of the Netherlands and in the Vecht basin. Megalithic monuments were built only in the ice-sculpted areas of the northeastern Netherlands. Only there had the land-ice of the penultimate glaciation brought sufficient boulder material to build these monuments. West of that area, TRB settlement sites have been found only on the western fringes of the sandy 'uplands' of the Veluwe, but none in the south of the Netherlands.

Although they were contemporary with the TRB people occupying the Pleistocene 'uplands', the overall impression is that the Vlaardingen traditions kept to their traditional living areas west of the sandy uplands. We think they were almost inextricably bound to the river deltas and maintained socially defined cultural boundaries with the TRB farmers. It is reasonable to hypothesize that those (immigrant) TRB populations would have relatively high proportions of Anatolian farmer-associated ancestry, similar to the TRB individuals that have been genetically analyzed from Denmark[67], but we are not aware of any TRB groups sampled in our research area.

Genetic studies demonstrate that the spread of the TRB culture, where it has been sampled, was associated with a new genetic signature, leading to the conclusion that '…individuals with hunter-gatherer ancestry persisted for decades and perhaps centuries after the arrival of farming groups in Denmark, although they have left only a minor genomic imprint on the population of the subsequent centuries'[67]. Nevertheless, the hunter-gatherer tradition was not completely 'eradicated' by the new wave of farming tradition. Along the Baltic, the southern Swedish and eastern Danish sea coasts, hunter-gatherer-fishers of the Narva, Nema, Zedmar and Pitted Ware traditions kept their own economic strategies and specific material culture until as late as 2200 BCE[68,69]. The Pitted Ware tradition appears in Scandinavia around 3400 BCE, with clear cultural similarity to economies further east. Genetic evidence also shows that Pitted Ware communities had mostly hunter-gatherer ancestry and did not mix with contemporary TRB, GAC and Corded Ware populations[67,70], even if they exchanged goods and probably practiced some agriculture as well [71].

### 5 Late Neolithic A (3000-2500 BCE)

| Genetic ID | Archaeological ID | Full Date | Locality |
|---|---|---|---|
| I12902 | R9560-11_V2764 | 2848-2501 calBCE (4088±19 BP) [R_Combine: (4100±20 BP, PSUAMS-8438), 2676-2433 calBCE (4010±50 BP, GrA-15698)] | Mienakker (NL) |
| I12896 | h 1973/3_18,19 | 2864-2500 calBCE (4100±30 BP, PSUAMS-7806) | Molenaarsgraaf (NL) |
| I33741 | A16-005_M94 | 2571-2468 calBCE (4015±20 BP) [R_Combine: (4015±20 BP, PSUAMS-12657), (3890±50 BP, GrA-15696)] | Sijbekarspel_de_Veken (NL) |
| I13631 | 213 / 15.019; AF008 | 2950-2600 BCE | Grotte du Mont Falise (BE) |
| I13638 | 213 / 3252; AF014 | 2886-2668 calBCE (4180±25 BP, PSUAMS-12142) | Grotte du Mont Falise (BE) |
| I13649 | 212 / 1x.021; AF026 | 3050-2600 BCE | Grotte du Mont Falise (BE) |
| I13651 | 212 / 3245; AF028 | 3050-2600 BCE | Grotte du Mont Falise (BE) |
| I13629 | 213 / 3259; AF006 | 2950-2600 BCE | Grotte du Mont Falise (BE) |
| I13631 | 213 / 15.019; AF008 | 2950-2600 BCE | Grotte du Mont Falise (BE) |
| I13633 | 97; AF010 | 2916-2781 calBCE (4260±25 BP, PSUAMS-12141) | Abri Sandron (BE) |
| I13635 | 89 / 6763; AF012 | 2660-2468 calBCE (4035±30 BP PSUAMS-11920) | Abri Sandron (BE) |
| I13654 | X1; AF031 | 2950-2650 BCE | Abri Sandron (BE) |
| I13655 | 88 / 6165; AF032 | 2950-2650 BCE | Abri Sandron (BE) |
| I13656 | 90; AF033 | 2950-2650 BCE | Abri Sandron (BE) |
| I13657 | 91; AF034 | 2917-2786 calBCE (4265±25 BP, PSUAMS-12143) | Abri Sandron (BE) |
| I13659 | 94; AF036 | 2911-2706 calBCE (4245±25 BP, PSUAMS-12144) | Abri Sandron (BE) |
| I13660 | 95; AF037 | 2865-2502 calBCE (4103±29 BP, UBA-42622) | Abri Sandron (BE) |
| I7012 | BELG_267 | 2618-2468 calBCE (4020±25 BP, PSUAMS-7872) | Grotte Rousseau (BE) |

*Table 5 Summary of samples used for analysis from the period 3000-2500 BCE*

#### 5.1 Summary of the archaeological context of the samples in table 5; for the extended version we refer to SI 2

Three individuals are from a Dutch context: two from the northwest, and one from the central Netherlands (I12896, I12902 and I33741). IBD analysis indicates that I33741 from Sijbekarspel_de Veken and dated to 2571-2468 calBCE had a distant kinship (approximately 8$^{th}$ degree) with I12902 found in Mienakker and dated to 2848-2501 calBCE. The individual from Molenaarsgraaf was a chance find and virtually without cultural context[72]. A tooth from the upper jaw of this individual of 30-40 years old (RM1973/18,19) yielded sample I12896 dated directly to 2864-2500 calBCE (4100±30 BP, PSUAMS-7806).

The Belgian samples cited above are all from the Ardennes, near the river Meuse or its tributaries, from caves that were used for multiple interments over long periods of time. A challenge with these collective burials is that most of them were accessible for long time periods and they may represent multiple burial events, sometimes separated by millennia. However, while this tradition of burying people in caves in this region had already started in the Mesolithic, the vast majority of them date to the turn of the 4$^{th}$ to the 3$^{rd}$ Millennium BCE[73,74].

Abri Sandron was excavated in the late 19th Century and details are limited. Eight individuals are reported in this study and five have been directly radiocarbon dated to the Late Neolithic. In other publications three additional radiocarbon dates on human remains confirm this general time frame and suggest a short period of use for this cave[73]. While we cannot exclude that some of those without direct radiocarbon dates may date to other time periods, their genetic profile corresponds with this time-frame. One individual sampled from the Grotte Rousseau, was directly radiocarbon dated to the early 3rd Millennium BCE.

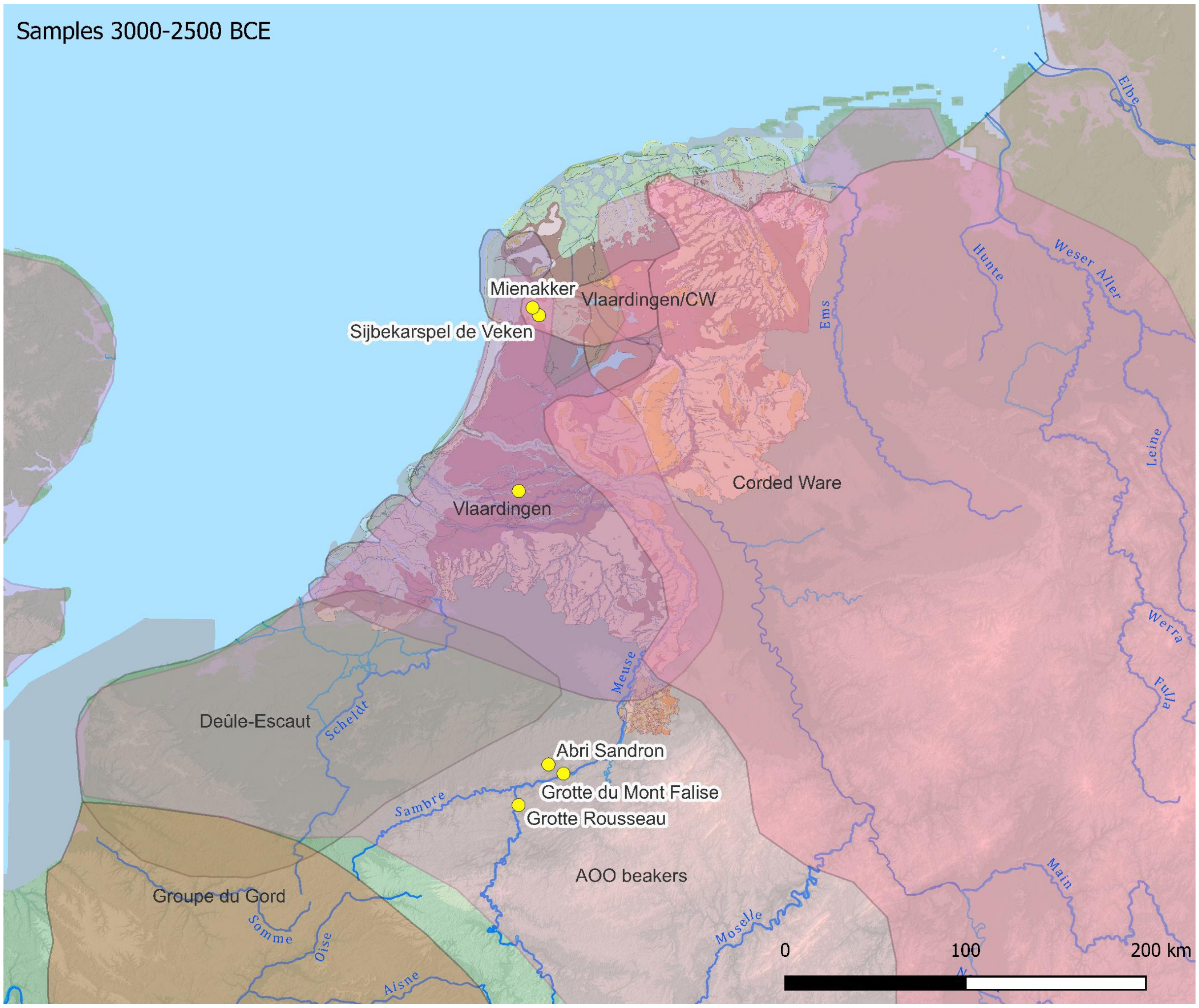

*Figure 5 Schematic distribution of cultural spheres and the geographic locations of samples from 3000-2500 BCE. The paleogeographic reconstruction of the Netherlands 2850 BCE is after Vos et al.*[4]. *The elevation map is from* *[https://www.mapsforeurope.org/datasets/euro-dem](https://www.mapsforeurope.org/datasets/euro-dem)* *(the grey area in the English Channel is uncharted).*

Although other samples from the same cave have been directly dated to the Mesolithic (see I7018), additional radiocarbon dates suggest that most individuals buried in this cave dated to the early part of the 3rd Millennium BCE[73].

The Grotte du Mont Falise was excavated in the late 19th, and early 20th centuries[75]. The context is not very clear but multiple individuals were discovered inside the cave. We included five individuals in this study, with one being directly radiocarbon dated to the first half of the 3rd Millennium BCE. Two additional radiocarbon dates from this site, support this time frame[74]. As with Abri Sandron, we cannot be certain that all sampled individuals date to this period, although it is most likely based on their genetic profile. 5.2 Cultural dynamics in the Late Neolithic A: 3000-2500 BCE

In the 3rd millennium several new traditions are recognized. The geographic distribution of these 'groups' is not very clear, however (*Figure 5*). Blanchet[76] and Brunet *et al*[77] have published generalized site distribution maps that situate the (late) SOM in the Paris Basin, reaching up to along the Oise to the River Meuse[76,77]. The distribution of the Groupe du Gord appears to concentrate in the Marne and Oise valleys, though the eponymous site lies near Compiegne, on the Somme river[76–79]. The Groupe du Deûle-Escaut is situated in the area north of the Somme[77], though most known sites are situated north of the heights of Artois, in the river valleys of the Deûle, Leie and Scheldt (Escaut)[76,77,80,81]. The location of the settlements is similar to those of the Vlaardingen Culture: on elevated areas in the river valleys and deltas. Blanchet[80], for instance, describes both SOM and Gord / Deûle-Escaut as situated mostly in river valleys with an economy that depends on farming, but also on a substantial amount of hunting and fishing. Also in Belgium, the Deûle-Escaut group is described as locating its settlements next to rivers and marshes[82–84] for access to 'food and different sources from wet environments'[85].

Both the Groupe du Gord and du Deûle-Escaut are dated to between 2900 and 2550 BCE[60]. They are therefore contemporaneous with Late Vlaardingen, late TRB and the Corded Ware in the Netherlands. From different sources it is clear there were contacts between the people using these regionally slightly different ceramic assemblages. These connections are visible for instance in similar types of ceramic artifacts, notably collared flasks, clay discs, ceramic 'spoons' and similarities in the technology and forms of pottery. They are also visible in the distribution of typical 'Vlaardingen' flint axes and of flint sources, with large blades of Grand-Pressigny flint as its 'apotheosis'. Grand-Pressigny flint is one of the typical elements in Deûle-Escaut sites[82]. Its distribution connects the Netherlands with central France through the regions in which the Artenac, Gord and Deûle-Escaut traditions are situated, along major rivers in these areas, including the river Meuse[86–88].

Taken together, the second half of the 4th and the first half of the 3rd millennium in Northern France, Belgium and the Western Netherlands was characterized by farming communities that supplemented their existence with hunting, fishing and gathering, and situated their settlements accordingly. Even though few reliable structures (houses) are known, these do have similarities. Houses were rectangular, often slightly trapezoidal, substantial, and well built[83,89–92]. They represent stable settlement locations, even if these are situated in areas that in modern eyes would not optimally be suited for arable farming. Yet they offered the possibility to optimally use the sources available in these varied environments.

The dead were buried in cemeteries in or near settlements (Vlaardingen), or in megalithic monuments that were built in the 4th millennium BCE (SOM, Wartberg, TRB) and often reused in the 3rd millennium (Gord, Deûle-Escaut, Bell Beaker) (c.f. Blanchet[76]). These practices imply that we have reasonably good context data from Vlaardingen sites, even though only one well documented cemetery is known[64]. In contrast, the megalithic graves of contemporaneous communities contain open and therefore less well dated burial assemblages. This is especially the case with the collective burials in caves and rock shelters. This practice is common along the Meuse river and its tributaries in the south of Belgium and northern France (cf. Toussaint *et al*[74]). The many caves in the region were used for collective burial from the 5th and into the 3rd millennium BC[74,93,94]. The remains are often heavily commingled and there is evidence of post-mortem manipulation of the bones. Most of the burials lack accompanying archaeological material, but those that do, contain artifacts typical for this time period, such as arrowheads or pottery shards of the respective archaeological cultures[74].

In paragraph 4.2 we already sketched the differences in settlement area between the Vlaardingen culture in the west, almost inextricably bound to the wetlands of the Rhine and Meuse valleys, and the TRB farmers that arrived from the east and settled on the higher ice-sculpted plateaus. Early and Middle Neolithic remains on these 'uplands' are very scarce, if not completely absent. We assume these regions were heavily forested and used for hunting and gathering, but not (yet) for settlement. Therefore the TRB farmers are considered to be the first to reclaim these forested areas and convert them into arable land and settled areas. While the oldest TRB sites are known from the wet areas of the Vecht-basin and on the flanks of the ice-sculpted ridges of Wiering and Texel in the north-west, its main distribution, and all of their megalithic monuments, are in the northeast, on the Hondsrug area, and down to the ice-sculpted ridges of the Veluwe in the central Netherlands.

This is also the landscape in which the Corded Ware complex settled. There is increasing evidence now that TRB groups remained in the area until at least 2600 BCE, therefore co-inhabiting the region with Corded Ware people[95,96] (Kroon 2024; Bourgeois *et al.* in press.). Here we have clear evidence of a transition to the typical Single Grave burial rite that pervades throughout Europe at that time (cf. Furholt 2021a), while at the same time, there is persistence of TRB communities well into the 3rd Millennium BC (Bourgeois *et al.* in press). For instance, cremation burials from the same time horizon as early CW burials have been found associated with TRB pottery[95,96].

West of these 'uplands', in the Rhine-Meuse delta we find only settlements of the Vlaardingen culture, but also with Corded Ware pottery. This is especially the case of the West-Frisian sites in the northwest, but also in other Vlaardingen sites in the western Netherlands. For a long time, these sites have been labeled as Corded Ware sites, but it has become clear that this Corded Ware pottery was not made by immigrant potters, but was made in a Vlaardingen tradition, with Vlaardingen clays and most likely by Vlaardingen potters[95,97–99]. Interestingly Kroon also showed that a substantial number of these pots have inclusions that only could have derived from tertiary deposits like in the Ardennes[98]. Vlaardingen potters therefore most probably had connections to people living higher up in the valley of the Meuse where such tertiary inclusions are probably eroded into natural clays.

In accordance with the cultural situation outlined above, the three individuals (I12896, I12902, I33741) sampled from the western Netherlands are labeled as Vlaardingen/Corded Ware individuals. These three individuals were living on the absolute fringes of the Corded Ware expansion prior to 2600-2500 BC. They were not buried in a classical Corded Ware burial style, but without grave goods and in 'flat graves' in settlements with both Corded Ware Beakers and Vlaardingen pottery and artifacts. These Corded Ware Beakers included All Over Ornamented (AOO) Beakers, which have a somewhat wider distribution than other Corded Ware Beakers (*Figure 6*). They are for instance found in Belgium, Northern France and in the British Isles[100–102], and appear to represent the enhanced mobility that especially marks the following Bell Beaker complex.

Traditionally in the Netherlands, AOO beakers are seen as the latest phase of the CW tradition, for instance because they are still associated with flint knives or 'daggers'[103,104]. In $^{14}$C-dates, however, there is a complete overlap with other CW dates, and there is no reason to separate AOO pottery as a distinct phase of Corded Ware[95].

### 6 Late Neolithic B and Early Bronze Age (2500-1700 BCE)

| Genetic ID | Archaeological ID | Full Date | Locality |
|---|---|---|---|
| I5748 | skeleton 575 (Jan) | 2579-2211 calBCE (3945±55 BP, GrN-6650C) | Oostwoud (NL) |
| I13028 | h 1982/7._4_Skelet I | 2500-2100 BCE | Ottoland_Kromme_Elleboog (NL) |
| I12900_en | h 1982/7._4_Skelet II | 2457-2145 calBCE (3820±45 BP, GrN-6384) | Ottoland_Kromme_Elleboog (NL) |
| I13025 | h 1967/1._Skelet I | 2136-1892 calBCE (3635±40 BP, GrN-5131) | Molenaarsgraaf (NL) |
| I13026 | h 1967/1._Skelet II | 2135-1890 calBCE (3630±40 BP, GrN-5566) | Molenaarsgraaf (NL) |
| I13027 | h 1967/1._Skelet III | 2197-1983 calBCE (3700±25 BP, PSUAMS-7847) | Molenaarsgraaf (NL) |
| I4069 | skeleton 229 | 2192-1887 calBCE (3640±50 BP, GrA-6477) | Oostwoud (NL) |
| I4073 | skeleton 236 | 2197-1897 calBCE (3660±50 BP, GrA-15598) | Oostwoud (NL) |
| I4074 | skeleton 242/533 | 2281-1899 calBCE (3690±60 BP, GrA-15597) | Oostwoud (NL) |
| I4075 | skeleton 243 | 2127-1933 calBCE (3635±20 BP, PSUAMS-2337) | Oostwoud (NL) |
| I5750 | skeleton 230 extra | 2050-1950 BCE | Oostwoud (NL) |
| I20063 | skeleton 232 | 1944-1766 calBCE (3530±25 BP, GrN-8801) | Oostwoud (NL) |
| I4067_en | skeleton 127 | 1958-1646 calBCE (3500±50 BP, GrA-15602) | Oostwoud (NL) |
| I4070 | skeleton 230 | 1880-1627 calBCE (3440±40 BP, GrA-17225) | Oostwoud (NL) |
| I4071 | skeleton 231 | 1883-1634 calBCE (3450±40 BP, GrA-17226) | Oostwoud (NL) |
| I4076 | skeleton 247 | 1883-1747 calBCE (3490±20 BP, PSUAMS-2319) | Oostwoud (NL) |

*Table 6 Summary of samples used for analysis from the period 2500-1700 BCE*

#### 6.1 Summary of the archaeological context of the samples in table 6; for the extended version we refer to SI 2

The samples presented in table 6 are from three sites in the Netherlands. The Molenaarsgraaf site is one of the best documented Bell Beaker / Early Bronze Age settlement sites in the Netherlands[72]. The Molenaarsgraaf individuals were buried in this settlement context. Skeleton 1 (I13025) was buried with a Bell Beaker near his feet but the other two individuals were buried without Beakers, although one of them was accompanied by fishing gear. The settlement situation is typical for a combined farming-hunting-fishing economy. The Ottoland-Kromme Elleboog settlement site is close-by and has the same date[72]. Two individuals were found buried in flat graves in a settlement context. The Oostwoud site consists of two burial mounds; one with 12 skeletons dated to the Late Neolithic Bell Beaker Period and the Early Bronze Age, and a second one with two skeletons dated to the Early or Mid-Bronze Age. Both burial mounds were located in extensive arable land with many Late Neolithic and Early Bronze Age Beaker shards[105].

##### 6.2 Cultural dynamics in the Late Neolithic B and Early Bronze Age: 2500-1700 BCE: Bell Beaker and Barbed Wire Beaker

Around 2500 BCE the situation appears to change considerably. The CW and AOO pottery styles are replaced by the Bell Beaker complex almost everywhere in the Netherlands, both in graves and settlements. Also, a large scale genetic turnover becomes visible with the advent of Bell Beaker groups in the region. Olalde *et al*[106] already showed the presence of steppe-related ancestry in Oostwoud, but samples from previous periods were lacking in this paper. It was therefore not clear if this steppe-related ancestry was already present, or to what extent, in other groups, such as Corded Ware communities. Bell Beaker burial mounds are restricted mostly to the ice-sculpted uplands again, but also extend into the sandy parts of the southern Netherlands and Belgium. Apart from the Oostwoud burial mound, no others are known in the western Netherlands. However, settlement sites are discovered everywhere, even in the Rhine-Meuse delta.

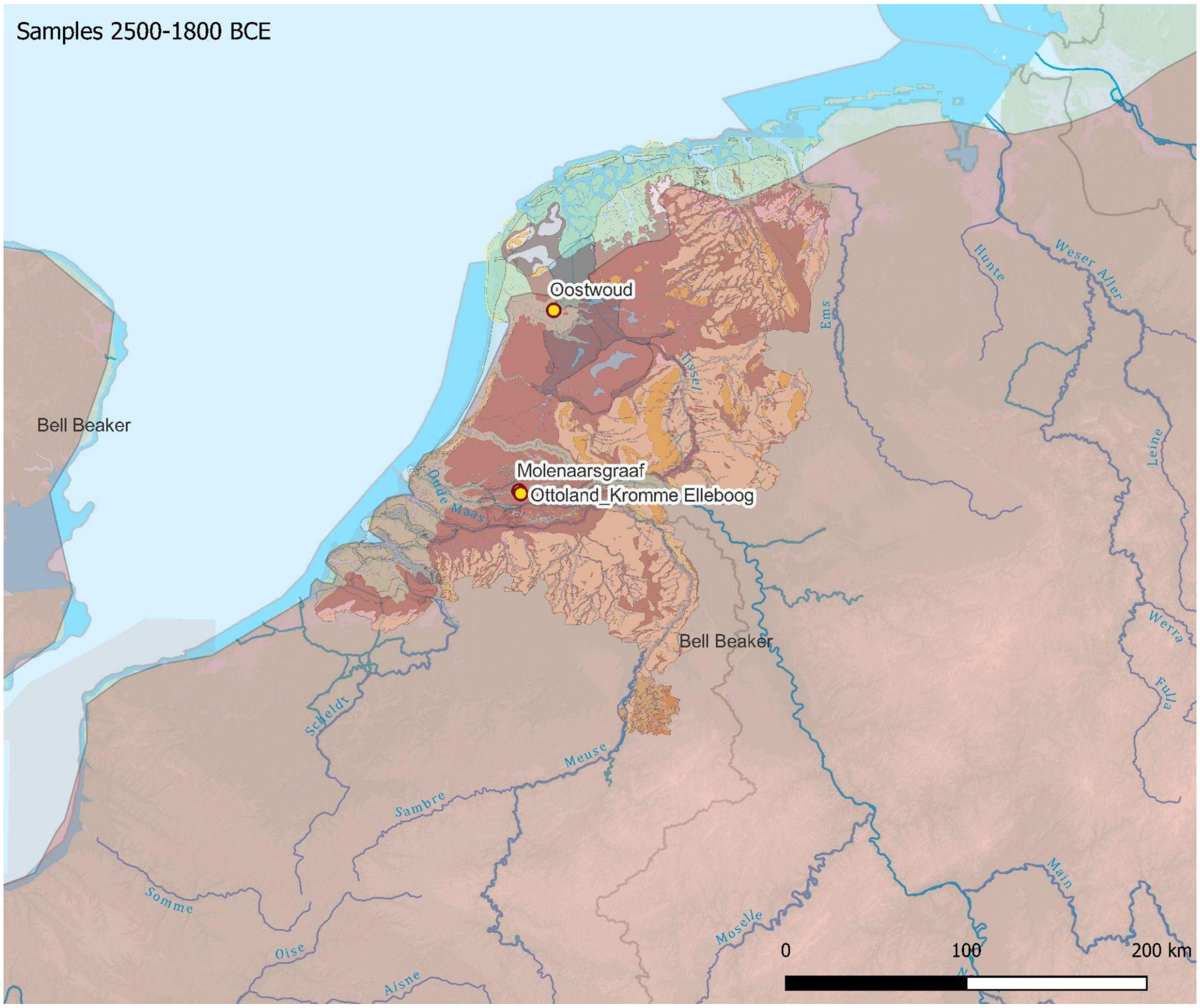

*Figure 7 Schematic distribution of cultural spheres and the geographic locations of newly analysed and published samples 2500-1800 BCE. The paleogeographic reconstruction of the Netherlands 2750 BCE is after Vos et al.*[4]*. The elevation map is from* https://www.mapsforeurope.org/datasets/euro-dem *(the grey area in the English Channel is uncharted).*

They are situated in similar locations as their Vlaardingen/Corded Ware predecessors, yet not on the exact same locations. Bell Beaker sites in the west, but also on the eastern uplands, are still situated in river valleys and in general in locations that allowed them to also hunt and fish[41].

In respect of culture therefore, settlements of the Bell Beaker complex do not offer a major break with those of the Vlaardingen/Corded Ware complex in the Rhine-Meuse delta. How they relate to settlements in the central and eastern Ice-sculpted uplands is not clear, because settlements are virtually lacking in that region. If we find them (for instance at Oldeboorn[41] and Hattemerbroek[107]), they are situated in locations one would not expect for full farming communities: in low-lying wetlands and in river valleys. They seem to be chosen to optimize the exploitation of the landscape, rather than focusing on farming. It should be noted, however, that present-day construction works are typically not conducted in these types of locations, making it less likely for them to be investigated by development-led archaeology[41].

Regionalisation of Beaker styles is visible in most countries during the course of the 3rd Millennium BCE [100–102], but this is an observation that is almost exclusively connected to pottery style. Bell Beaker settlements remain very scarce (several in the Netherlands, but very few in the adjacent countries: see Besse[108]). Around 2000 BCE a last 'universal' phase of the Beaker complex arrived with the Barbed Wire Beakers. These occur everywhere where Bell Beakers were used, from Denmark down to southern France and eastward to Poland. Yet settlements are rare and even burial sites are scarce. In the Netherlands, only the Oostwoud burial mounds yielded samples from this period, which ended around 1800-1700 BCE, to be replaced by Middle Bronze Age material culture. In the Low Countries and adjacent areas, this was the period in which the three-aisled longhouse developed as a symbol of social and economic structures that probably were much different from the Late Neolithic Beaker period[109].

### SI 2. Archaeological context information about the newly reported and published individuals from the Rhine-Meuse area

Edited by Harry Fokkens, Quentin Bourgeois and Eveline Altena

### The Netherlands

#### 2.1 Hardinxveld-Polderweg (98POL) (Zuid-Holland, the Netherlands)

**Analyzed individual:**

I13024 (V28578): 5802-5626 calBCE (6820±50 BP, GrA-9804), 5500-5400/5300 BCE after correction of reservoir effect

**Contact information**: Liesbeth Smits, Leendert Louwe Kooijmans

**Site information and excavation history:** The Mesolithic and Early Neolithic habitation of this site was located on the top of a submerged river dune (*donk* in Dutch) of which 16 x 28 m was excavated in a trench reinforced with steel sheet piling to a depth of about 8.5 m below Dutch Ordinance Datum. All excavated soil was sieved, yielding thousands of organic remains and flint artefacts. The mesolithic part of the site was interpreted as a winter base camp for family groups of Mesolithic hunter-gatherers (Hamburg & Louwe Kooijmans 2001: 31).

Two human burials were found at the site, one of which was severely disturbed, and three dog burials, of which two were disturbed as well. Additionally, several dispersed human bones were found, belonging to at least another ten individuals (Smits & Louwe Kooijmans 2001: 428).

**Summary of the sampled materials:** For this project the undisturbed grave, grave 1, was sampled. It contained the remains of a 40–60-year-old woman (nick-named Trijntje) who had been buried in supine extended position. Her skeleton (find number 28.578), was poorly preserved and therefore limited further inferences about her physical state (Smits & Louwe Kooijmans 2001). The petrous bone yielded sample **I13024.**

**Dating**: The $^{14}C$ date for grave 1 is 5738-5588 cal BCE (GrA 9804: 6820 ± 50 BP). However, the $\delta^{13}C$ value was high (-22.60 %) and could imply a reservoir effect of about two centuries (Louwe Kooijmans & Mol 2001: 68, table 3.2.). The final date after correction for the reservoir effect is 5500-5400/5300 BCE (Dreshaj et al. 2023).

**Source of the sample**: Provinciaal archeologisch depot Zuid-Holland; Inge Riemersma, Mark Phlippeau. Sample collected by Eveline Altena.

**Source of the entry**: Liesbeth Smits, Leendert Louwe Kooijmans

**References**:

Dreshaj, M., Dee, M., Brusgaard, N., Raemaekers, D., & Peeters, H. (2023). High-resolution Bayesian chronology of the earliest evidence of domesticated animals in the Dutch wetlands (Hardinxveld-Giessendam archaeological sites). *PLOS ONE, 18(1), E0280619.* https://doi.org/doi.org/10.1371/journal.pone.0280619.

Hamburg, T. D., & Louwe Kooijmans, L. P. (2001). 1. Vooronderzoek en opgraving. In L. P. Louwe Kooijmans (Ed.), *Archeologie in de Betuweroute. Hardinxveld-Giessendam Polderweg: Een Mesolithisch jachtkamp in het rivierengebied (5500-5000 v. Chr.)* (Vol. 83, pp. 13–34). NS Railinfrabeheer.

Louwe Kooijmans, L. P., & Mol, J. (2001). 3. Stratigrafie, chronologie en fasering. In L. P. Louwe Kooijmans (Ed.), *Archeologie in de Betuweroute. Hardinxveld-Giessendam Polderweg: Een Mesolithisch jachtkamp in het rivierengebied (5500-5000 v. Chr.)* (Vol. 83, pp. 55–97). NS Railinfrabeheer.

Smits, L., & Louwe Kooijmans, L. P. (2001). 14. De menselijke skeletresten. In L. P. Louwe Kooijmans (Ed.), *Archeologie in de Betuweroute. Hardinxveld-Giessendam Polderweg: Een Mesolithisch jachtkamp in het rivierengebied (5500-5000 v. Chr.)* (Vol. 83, pp. 419–442). NS Railinfrabeheer.

#### 2.2 Zoelen-de Beldert (Gelderland, the Netherlands)

**Analyzed individuals:**

I33738 (skelet I): 4200-3800 BCE

I33739 (skelet II): 4227-3808 calBCE (5190±50 BP, UtC-1961)

**Contact information:** Roel Lauwerier

**Site information and excavation history:** North of Tiel, in the villages of Zoelen and Medel, very extensive gravel and sand extractions have taken place from the nineteen eighties onwards. This has led to the discovery of many sites from all kinds of periods, several of which could be excavated. At Zoelen-de Beldert, northwest of Tiel, local archaeologist E. Verhelst discovered a pit with skeletal material that triggered a small rescue excavation in May 1991 (Hogestijn & Lauwerier, 1992, p. 108).

**Summary of the sampled materials:** In the burial pit or grave the remains of three people were buried. Lowermost buried were skeleton II and III, both were buried in an extended prone position on a bed of organic material (leaves or bark). They were the skeletons of a 50–70-year-old female and a child of approximately 7 years old (Hulst et al., 1993; Lauwerier, 1993). The adult skeleton II had been placed on top of the child. This skeleton yielded **sample I33739**. The child's remains did not contain suitable material for aDNA analysis.

The top of the pit contained the secondary deposition of the remains of another 30–60-year-old woman (skeleton I, **sample I33738**). This individual was likely originally buried elsewhere and interred in the new grave only after the body had decayed (Hulst et al., 1993, p. 69). Swifterbant style pottery was found nearby, but not in the grave itself.

**Dating:** A $^{14}$C sample of the adult skeleton II (**sample I33739**) yielded a date of 5190+/- 50 BP (Utc-1961), which calibrates to 4227-3808 calBCE (Hulst et al., 1993). Currently there is a discussion about the reliability of these older Utrecht AMS dates: they could be too old. This may imply that the actual date is in the lower range of the 2-sigma range around 3800 BCE. The IBD matches (≥1 fragment of 12cM) with this individual correspond mostly to the second half of the 5$^{th}$ millennium and the first half of the 4$^{th}$ millennium BCE.

**Isotopes:** Enamel fragments from three maxillary molars (teeth 26, 27, and 28) belonging to Skeleton II (**sample I33739**) were available for combined Sr-O-C isotope analysis (see Supplementary Information S3). The results are presented in Table S2.2. All $^{87}$Sr/$^{86}$Sr are compatible with the expected location bioavailable Sr signature in the Zoelen region, i.e., the Dutch central river area (Kootker et al., 2016). The $\delta^{13}C_{PDB}$ values are relatively low compared to the majority of the analysed individuals from the Netherlands (all periods, unpublished data from L.M. Kootker). Such values may indicate that the female's diet during her first 16 years of life mainly consisted of $C_3$ plants, which were likely sourced from forested or temperate environments. The $\delta^{18}O_{PDB}$ values vary (max-min: 1.53 ‰) and are all, except for the $M^1$, comparable to the values we believe are indicative of "the Netherlands" (Kootker et al., 2019). The –7.3 ‰ of the first molar ($M^1$) currently falls outside this range, which at first glance cannot be directly linked to breastfeeding, as the $\delta^{18}O$ of the $M^1$ in that case would usually be heavier (i.e., more positive) than the third molar ($M^3$). It is possible that the woman spent her youth somewhat more inland, possibly to the west of present-day

Germany. A limiting factor in the interpretation that should also be considered is that the variation in $\delta^{18}O$ within a single molar can be significant (up to about 2‰ in modern molars: Plomp et al., 2020).

**Table S2.2.** Sr-O-C isotope data for Skeleton II from Zoelen-de Beldert. The element numbers are according to Fédération Dentaire Internationale (FDI).

| Sample ID | Material | Element | $^{87}Sr/^{86}Sr$ | 2SE | $\delta^{13}C_{PDB}$ (‰) | SD | $\delta^{18}O_{PDB}$ (‰) | SD |
|---|---|---|---|---|---|---|---|---|
| Skelet II (I33739) | Enamel | 26 | 0.708914 | 0.000007 | -14.43 | 0.05 | -7.28 | 0.06 |
| | | 27 | 0.708736 | 0.000007 | -15.20 | 0.05 | -5.75 | 0.06 |
| | | 28 | 0.708856 | 0.000008 | -15.27 | 0.07 | -6.69 | 0.04 |

**Source of the samples:** Archeologisch Depot Gelderland; Stephan Weiss-König. Samples collected by Eveline Altena.

**Authors of the entry:** Roel Lauwerier, Harry Fokkens, Lisette Kootker

**References**:

Hogestijn, J. W. H., & Lauwerier, R. C. G. M. (1992). Zoelen Beldert. *Jaarverslag Rijksdienst Voor Het Oudheidkundig Bodemonderzoek 1991*, 108.

Hulst, R. S., Hogestijn, J. W. H., de Haan, M. J. A., Lauwerier, R. C. G. M., & Marswijk, R. W. (1993). Buren Zoelen. *Jaarverslag Rijksdienst Voor Het Oudheidkundig Bodemonderzoek 1992*, 69.

Kootker, L.M., Van Lanen, R.J., Kars, H., Davies, G.R. (2016). Strontium isoscapes in the Netherlands. Spatial variations in 87Sr/86Sr as a proxy for palaeomobility, Journal of Archaeological Science: Reports 6, 1-13.

Kootker, L.M., van Lanen, R.J., Groenewoudt, B.J., Altena, E., Panhuysen, R.G.A.M., Jansma, E., Kars, H., Davies, G.R. (2019). Beyond isolation: understanding past human-population variability in the Dutch town of Oldenzaal through the origin of its inhabitants and its infrastructural connections, Archaeological and Anthropological Sciences 11, 755-775.

Lauwerier, R. C. G. M. (1993). *Zoelen—Beldert 2-5-1991; neolithische menselijke skeletten (2)* [Intern verslag Archeozoölogie / ROB 2 februari 1993].

Plomp, E., von Holstein, I.C.C., Kootker, L.M., Verdegaal-Warmerdam, S.J.A., Forouzanfar, T., Davies, G.R., (2020). Strontium, oxygen, and carbon isotope variation in modern human dental enamel, Am J Phys Anthropol 172, 586-604.

#### 2.3 Nieuwegein-het Klooster location A (NGKL10) (Utrecht, Netherlands)

**Analyzed individuals:**

I12091 (S035_ID1): 4400-3900 BCE

| | | |
|---|---|---|
| I17968 | (S157)ID4): | 4400-3900 BCE |
| I12093 | (S292_ID5): | 4200-4000 BCE |
| I12094 | (S292_ID6): | 4200-4000 BCE |

**Contact information:** Kirsten Leijnse, Helle Molthof, Theo ten Anscher

**Site information and excavation history:** The development of a business park, 'Het Klooster' in Nieuwegein, the Netherlands, fueled a large-scale archaeological project. The nowadays flat green meadows are hiding a former river system called the Wiersch, a predecessor of the Rhine, with levees and peat-filled channels. The initial surveys indicated that the area was intensively used by Neolithic communities. One of the most promising locations, discovered by augering, was a dense scatter of archaeological indicators (pottery, flint, bone and charcoal) on a former river belt, 1,7 meters below the present-day surface. The site report, due to be published in 2025, will encompass three excavated areas; the information below is limited to the largest of these, known as 'location A'.

This Neolithic settlement was excavated in 2016 and yielded a find layer with tens of thousands of sherds, flints, animal bones and stones. Based on the typology of pottery and flint artefacts, the larger part of the site is dated to the second half of the Swifterbant culture (SW2; 4400–3900 cal BCE). In the southern part of the site there are indications for an older phase (SW1; 4900–4400 cal BCE). This latter part shows an emphasis on blade technology while relatively 'late' elements are absent, such as fragments of polished flint axes, triangular flint points with surface retouch, and transverse arrowheads. In contrast, the northern part of the site shows a marked shift in flint technology with a stronger emphasis on flake production, in line with SW2. The aforementioned 'late' elements are also present in this northern part and probably date to the period immediately after SW2, i.e. Pre-Drouwen TRB (3900 - 3400 cal BCE), in accordance with yet another shift in flint technology.

Most strikingly, location A yielded the remains of approximately 25 human individuals. These remains seem to be confined to three zones within the excavated area: zone 1 is located in the north of the site, zone 2 approx. 50 m to the south, and zone 3 is located another 100 m further south. Most individuals are represented by human bone fragments and teeth from the sieving residue of sampling units. Some of them probably represent burials that unfortunately were not recognized as such during the excavation, some are anciently disturbed graves, partial (re)burials or stray human remains. Five individuals were recognized as inhumations during fieldwork: three of these in close proximity in zone 2 (ID2, ID3 and ID4) and two together in a grave in zone 3 (ID5 and ID6). In addition to the five individuals in graves, a skull without a mandible (ID1) was found a few meters west of the group of ID2/3/4. The graves and the isolated skull were block lifted with the surrounding clay and transported to a hall for further investigation, to ensure dry and proper research conditions.

Grave goods were found with one individual (ID4; I12092/I17968), consisting of eight perforated cattle incisors[1] found in proximity to the neck. Apart from these, no grave goods were found. However, the sampling units yielded three jet objects, a pendant and two beads, which were found in or near areas with human remains. The pendant found near the human remains of zone 1 measures approx. 6 x 6 cm, which makes it the second largest prehistoric jet object of the Netherlands so far. Although the three ornaments could not be directly linked to buried individuals or human remains, all three were located near zones with human remains, while the areas in-between did not yield any jet finds. It is possible that the jet objects at Nieuwegein were originally deposited as grave goods.

**Summary of the sampled materials:** A total of thirteen samples, derived from eleven individuals, was analyzed for aDNA, but only four of the samples, two from zone 2 and two from zone 3, all petrous bones, delivered results.

The first, from zone 2, was from the isolated skull (ID1). This skull, without teeth or a mandible, belonged to an adolescent of 14–18 years and yielded sample **I12091** (male). The second sample was taken from ID4, found about four meters to the east of the skull, an almost complete skeleton of an adolescent of 16–20 years old (sample **I17968;** female). This individual was buried in an extended supine position, and around the neck eight perforated cattle incisors were found (probably domesticated, not aurochs).

In the most southern part of the site, zone 3, one grave was discovered containing the remains of two people. The first (ID5) was an almost complete skeleton of a young female adult of 20–24 years old, buried in extended supine position. Her right arm was bent, with the hand resting near the hip bone, and her head was turned towards the right. On her right side, cradled within the bent right arm, the remains of a neonate (0–3 months) were found (ID6). Both individuals were successfully analyzed for aDNA, yielding sample **I12093** (ID5: female) and **I12094** (ID6: female). The DNA analyses not only revealed that the baby is a girl, but also that the woman and the baby are first degree relatives: mother and daughter. It is the first time for the Stone Age of the Netherlands that we are able to take family ties into account and determine the sex of such a young infant.

**Dating:** The samples from zone 2 (ID1: I12091, ID4: I17968) were radiocarbon dated between 5200-4680 cal BCE. ID1 dates to 5868 ± 19 BP (GrM-13971; 4800–4680 cal BCE), while for ID4 two $^{14}C$ dates failed, but the third delivered a date of 6050 ± 40 BP (ICA-6959; 5200–4837 cal BCE). When compared to the typological dates of associated material, it can be concluded that these dates are too old and probably biased by, amongst other factors such as post depositional processes (see below), a freshwater reservoir effect, making them seem older than they are. The $\delta^{13}C$ values of these skeletons were subsequently − 22.18 ‰ and −22.36 ‰, which confirms the reservoir effect due to the consumption of freshwater fish. The flint and pottery found adjacent to the burials in zone 2 point to occupation mainly in the SW2 period (4400–3900 cal BCE).

One of the perforated incisors, found as grave goods at ID4, was $^{14}C$ dated to 5410 ± 30 BP (ICA-6954; 4345-4165 cal BCE). A concentration of charred hazelnuts found in the sieved layer above ID2/3/4 yielded a date of 5460 ± 30 BP (ICA-5755; 4365-4240 cal BCE). The latter dates, with the result for the charred hazelnuts as a *terminus ante quem*, seem to fit the other indicators much better and it can be assumed that the human remains found in zone 2 should also be placed within the SW2 phase.

Of ID5 in zone 3 (sample **I12093**) two $^{14}C$ samples, both from the adult individual, yielded dates of 5930 ± 15 BP (GrM-13461; 4845-4720 cal BCE) and 5920 ± 40 BP (ICA-6960; 4910-4695 cal BCE). As with the samples from zone 2, there was a big risk that the dates on the human material would turn out to be too old due to a reservoir effect. The $\delta^{13}C$ values were subsequently − 22.91 ‰ and −21.55 ‰, confirming these fears. Nevertheless, when looking at the flint found in zone 3, this material does point towards occupation in the SW1 phase (4,900–4,400 cal BCE), older than the northern and mid-section of the site, although later occupation (SW2 and Pre-Drouwener TRB) is found in the southern part as well. Several $^{14}C$ dated features in the vicinity of the buried mother and child also point to usage of this part of the terrain during SW1 (as well as later periods). However, these older features are, without exception, the remnants of surface hearths, covered by a layer of clay that has been deposited around 4200 cal BCE. The skeletal remains are found at exactly the same height as the nearby SW1 surface hearths, only a few meters away. Synchronicity can then only be assumed if the skeletons were also placed on the surface and remained there, undisturbed, for several centuries. The relatively sparse displacement of skeletal elements makes it more than likely that the bodies were buried in a shallow grave pit, dug in the clay layer that was dated to ca. 4200 cal BCE. Furthermore, below the mandible of ID5 a piece of charred ash was found, which was subsequently dated at 5720 ± 30 BP (ICA-6563; 4678-4458 cal BCE). Charcoal was not abundant in the fill of the presumed grave pit. This isolated fragment is interpreted as part of the infill, originating from a fire that dates before the burial event, and is therefore seen as a terminus post quem. All observations considered it is highly improbable that the burial of mother and child dates from before ca. 4200 BCE, regardless of the $^{14}C$ results from the human remains.

The $^{14}C$ dates of unburnt human bone, and unburnt bone in general, proved more than problematic; most results on these materials from Nieuwegein (and elsewhere, such as Tiel_Medel - de Roeskamp) had to be classified as unreliable. This was already expected as a result of the observed soil formation processes, in this case especially the decalcification of the uppermost part of the soil, to a depth of 50 to 100 cm below the surface. Amongst many other effects, decalcification and specifically the byproducts of calcium carbonate dissolution such as calcium oxide and calcium hydroxide, have a strong detrimental effect on bone, especially the collagen therein, but can also lead to increased exchange of elements (i.e. carbon) with the surrounding matrix (a more detailed description shall be part of the site report, including some analyses with multiple $^{14}C$ dates from the same context using different sample materials). The mentioned processes, with these effects, are not specific to Nieuwegein, but can also be expected in other clay areas (fluviatile and marine) that were originally calcareous but underwent (strong) decalcification.

**Isotopes:** The tooth enamel quality of the Nieuwegein individuals was moderate. To minimize contamination, only dental elements that produced clean, white enamel powder during sampling were selected for analysis. As a result, only two of the four individuals included in this study, ID4 and ID5, were sampled for Sr-O-C isotope analysis (Supplementary Information S3). However, only the Sr isotope data will be presented here. For ID4, all three molars from the right maxilla were sampled, while for ID5, all three molars from the left mandible were analysed. The data are presented in Table S2.3. The measured $^{87}Sr/^{86}Sr$ vary significantly, both between individuals and among the molars of a single individual (0.7086–0.7096). According to the current Sr isoscape map (Kootker et al., 2016), the lowest ratios are relatively low for the region and even for the present-day Netherlands. However, in the Neolithic, Sr isotope ratios below 0.7088 were more commonly observed in the Dutch river area (unpublished data from L.M. Kootker). This suggests that such low values likely occurred frequently in the prehistoric riverine landscape. Overall, the observed Sr isotope ratios are consistent with expectations for a geologically heterogeneous region such as present-day Nieuwegein. In contrast, the oxygen isotope data (not given here) seem to indicate possible non-local origins. However, the obtained $\delta^{18}O_{PDB}$ values require confirmation through $\delta^{18}O_{PO4}$ analysis, as phosphate-derived values from apatite are considered less susceptible to diagenetic alteration than those from carbonate.

**Table S2.3.** Sr isotope data for female individuals ID4 and ID5 from Nieuwegein-Het Klooster. The element numbers are according to Fédération Dentaire Internationale (FDI).

| Sample ID | Material | Element | $^{87}Sr/^{86}Sr$ | 2SE |
|---|---|---|---|---|
| V791.1/ID4 (I17968) | Enamel | 16 | 0.708702 | 0.000009 |
| | | 17 | 0.709166 | 0.000010 |
| | | 18 | 0.708622 | 0.000010 |
| V790.3/ID5 (I12093) | Enamel | 36 | 0.709646 | 0.000009 |
| | | 37 | 0.709075 | 0.000008 |

| | | | | |
|---|---|---|---|---|
| | | 38 | 0.709289 | 0.000009 |

The $^{87}Sr/^{86}Sr$ of female individuals ID4 (**I17968**) and ID5 (**I12093**) vary significantly among different dental elements, suggesting residential mobility during childhood or shifts in their primary food sources. For ID4, at least two periods of mobility can be identified: after the third year and around the eighth year of life. In contrast, ID5 exhibits evidence of only one mobility event, occurring after the third year. The isotopic difference between the second and third molars in ID5 is minimal (<0.0002, see Plomp et al., 2020), which is insufficient to indicate significant mobility.

**Source of the samples: RAAP archeologisch Adviesbureau.** Samples collected by Eveline Altena.

**Authors of the entry:** Kirsten Leijnse, Helle Molthof, Paul van der Kroft, Theo ten Ancher, Lisette Kootker

**References:** The site is due to be published in 2025

Kootker, L.M., Van Lanen, R.J., Kars, H., Davies, G.R. (2016). Strontium isoscapes in the Netherlands. Spatial variations in $^{87}Sr/^{86}Sr$ as a proxy for palaeomobility, Journal of Archaeological Science: Reports 6, 1-13.

Plomp, E., von Holstein, I.C.C., Kootker, L.M., Verdegaal-Warmerdam, S.J.A., Forouzanfar, T., Davies, G.R. (2020). Strontium, oxygen, and carbon isotope variation in modern human dental enamel, Am J Phys Anthropol 172, 586-604.

[1] Among the cattle incisors, a wild horse incisor was also found. Skeletal elements of horses are very scarce in the animal bone assemblage of site 1, so it is possible that it concerns another grave good. However, the tip of the root -where a perforation would have been- is missing. Therefore, we cannot know for certain that this is a modified ornament

#### 2.4 Tiel Medel de Roeskamp (NBBM6) (Gelderland, the Netherlands)

**Analyzed samples:**

I35542 (grave 4_S1137_V4957): ca. 3700 BCE

I35543 (grave 1_S1147_V2084): ca. 3700 BCE

I35544 (grave 3_S2386_V4910): ca. 3700 BCE

**Contact information:** Theo ten Anscher

**Site information and excavation history:**

At Medel -De Roeskamp, in the Dutch central river district, large-scale excavations were carried out between 2016-2017 . In the excavated area of approximately 5 ha, stratigraphically separated prehistoric landscapes were uncovered, revealing settlement remains dating to the Early Neolithic Swifterbant culture, the Middle Neolithic Hazendonk Group and the Early Bronze Age, with traces extending to the late Neolithic and well into the late Bronze Age.

Situated on a crevasse/levee along a channel of the river Rhine system approximately 1,5 ha of a large Swifterbant settlement was excavated, yielding more than 550.000 finds. Around 40 $^{14}$C-dates place the settlement firmly in Phase SW2 (4400-3900 BCE). Occupation began around 4300 with the clearing of the local hardwood riparian forest. Over time, the rapidly rising groundwater table and the associated deposition of clay sediments in the surrounding floodplain and channel had an initially imperceptible but steadily worsening impact on the levee's suitability as a settlement location. The area that remained above the surrounding floodplain gradually diminished. From 4050 BCE onwards, the site was only sporadically visited. By that time, the levee was largely submerged, while the local channel had become a narrow and shallow residual channel.
Approximately 30-40 two aisled, rectangular or slightly tapering house plans point to a permanent settlement consisting of at least a few houses at any one time. Evidence of local cereal cultivation is abundant, including about 2500 chaff and kernel remains of naked barley, einkorn, emmer wheat and free-threshing wheat (durum), as well as grain based products such as ground cereals and charred bread lumps. More than 75 % of all identified mammalian remains (7500 fragments identifiable to species or family level) are from livestock, dominated by large pigs kept for meat, and cattle kept for milk, meat, and as draught animals. Less than 25 % of mammalian remains belong to game, mostly red deer. In particular, bones of fur animals, abundant at other Swifterbant sites, are scarce. Fishing appears to have been substantial. Among the gathered food plants, hazelnuts and crab apples were prominent. Apart from a few scattered human remains, no traces of human graves were found on the levee. Some in situ skeletal remains indicate (partial) dog burials. Polished stone axes and a *durchlochte Breitkeil*, all found in fragments, as well as the spectrum of grains and arable weeds, ceramics (including vessel shapes, decoration rules based on pot format, and the application of a layer on large pots), some flint tools (particularly used for harvesting) and the house types, indicate strong Bischheim influences and shared technological and agricultural practices. This agrarian lifestyle – complemented by fishing, gathering, and some hunting – persisted at Medel until local environmental conditions made it unsustainable.

Around 3800 BCE, the old silted up channel was reactivated, causing extensive erosion of the former riverbank area of the Swifterbant settlement. The sediments in the old channel were also swept away. It took about a century for new inhabitable landscape elements to form over the area of the old channel. Point bars along the new channel, which subsequently silted up rapidly, attracted Hazendonk occupation. This habitation probably started around 3700 BCE and lasted until about 3500 BCE or slightly later, yielding approximately 75.000 finds. The Hazendonk settlement is characterized by a sequence of a few houses, with one occupied at a time. The subsistence economy was dominated by agriculture as is evidenced by a small number of chaff remains and kernels of naked barley and emmer wheat, as well as livestock, mainly cattle and pigs. Ca. 25 % of the mammal bones belong to game, particularly wild boar, followed by red deer and aurochs. In contrast to the Swifterbant period, very few charred hazelnut shells were recovered, suggesting that gathering played a less significant role. Fishing appears to have been of lesser importance too.

In the extensive site monograph, no graves are mentioned for the Hazendonk period (nor for the Swifterbant period). However, new aDNA information has led to a reinterpretation of the data: a flatgrave cemetery formerly ascribed to the Early Bronze Age is now believed to be much older and should be linked to the Hazendonk period. The arguments are outlined below.

Followed by a hiatus of ca. 1500 years a new occupation phase began in the Early Bronze Age (1950-1800 cal.BC), preceded by ephemeral traces of a Late Neolithic Bell Beaker settlement. Nearly 300.000 finds can be ascribed to this period. The Early Bronze Age settlement is characterized by several farmyards and associated structures, along with three monumental burial mounds containing dozens of inhumation and cremation burials. These mounds functioned as cemeteries that were in use from the Early Bronze Age, or perhaps the late Bell Beaker period, well into the Late Bronze Age. During its early phases the largest mound was encircled by a disrupted ditch and seems to have served as a solar calendar. When viewed from a central point within the disrupted circle, the gaps between the ditches align with the position of the rising sun during solstices and equinoxes.

180 m to the southwest of the burial mounds, a small flat grave cemetery was discovered. At the center of this cemetery, precisely aligned with one of the viewing gaps, a ritual pit was discovered. This pit contained animal bones, human skull remains (without the mandible or teeth), a glass bead – the oldest glass bead in the Netherlands, probably crafted in Mesopotamia – and several Early Bronze age sherds. The four surrounding individual graves 1-4 and grave 5, which contained the remains of at least 20 individuals, all lacked datable grave gifts. Only the rather poorly preserved skeletal remains were visible, with no traces of burial pits whatsoever. The teeth (and sometimes the premolars as well) of the adults, who appeared to be exclusively female, showed significantly more wear than the molars, indicating that they were used as tools. This trait was one of the key arguments for the contemporaneity of the individuals. About twenty $^{14}$C-samples were taken from teeth or bone, but none yielded reliable results due to poor collagen preservation as a result of soil formation processes. These soil processes are not restricted to Medel but can be expected for large areas in the Dutch river district.

Initially a Hazendonk date for the graves was contemplated, but when it became clear that the sherds found directly above and among the human remains belong to Bell Beaker and Early Bronze Age pottery, this was deemed highly unlikely. Since the ritual pit must date to the Early Bronze Age (see also below), while its upper fill contained human remains that are complementary to the headless skeleton from the adjacent grave 4, all the indications pointed to an Early Bronze Age date for the flat grave cemetery.

Several aDNA samples from this flat grave cemetery were attempted, but only three individuals provided aDNA samples. The new aDNA evidence, not available at the time, makes it clear that an Early Bronze Age date and all the hypotheses concerning the Early Bronze Age and Beaker burial rituals, has to be rejected. A new scenario that reconciles the archaeological data with the genetic evidence is proposed below.

**Summary of the sampled materials:**

Grave 1 (I35543) contained the remains of a 40–45-year-old female in a strongly flexed position, lying on her left side. No grave goods were associated with this burial.

Grave 3 (I35544) contained the remains of a 30–50-year-old female in a flexed position, lying on her right side. An awl crafted from a metacarpal or metatarsal bone of an herbivore (sheep, goat, or deer) was found within the grave.

Grave 4 (I35542) was partly disturbed, as the skull was discovered in a separate pit adjacent to the burial. The grave itself contained the headless skeleton of a 25–35-year-old female, buried in a strongly flexed position, lying on her right side. Within the upper fill of the pit, the heavily fragmented remains of what is likely her skull were discovered, accompanied by teeth from the upper and lower jar and some bones from the hand.

**Dating:** The $^{14}C$ -dates of bone and teeth are very problematic as the following demonstrates. The cal BCE ranges reflect the 95.4 % probability.

Grave 1: a sample of the right femur yielded no collagen. A piece of the right tibia was dated to 3090-2875 cal BCE (GrM-13370: 4315 ± 50 BP). One burnt fragment of einkorn found in grave 1 was radiocarbon dated to 4227-3947 cal BCE (GrM-24349: 5243 ± 29 BP). This date should be regarded as a *terminus post quem*.

Grave 3: a sample of the left tibia yielded no collagen. Tooth enamel (apatite) was dated to 1970-1885 cal BCE (GrM-13181: 3576 ± 15 BP). By an experimental method a fragment of the left femur (apatite) was dated to 3635-3374 cal BCE (GrM-20888: 4720 ± 40 BP).

Grave 4: no associated $^{14}C$-dates.

No $^{14}C$-dates could be obtained for grave 2. Samples of bone and teeth (dentine) from grave 5 yielded no or hardly any collagen and could not be dated. Three radiocarbon dates from tooth enamel (apatite) range from 2635 to 1530 cal BC. Two other teeth (apatite) from grave 5 were dated using an experimental method, which resulted in dates between 3335 and 2601 cal BC. Two small charred hazelnut shell fragments, sampled together, gave a date between 4331 and 4058 cal BC that should be regarded as a *terminus post quem*.

The individuals from graves 1, 3, and 4 have no steppe ancestry, pleading against a date in the Early Bronze Age. There are multiple IBD links to several individuals from predominantly England and Scotland dating to the first half of the 4th millennium BCE and an individual (**I33738**) from nearby Zoelen de Beldert (~4200-3800 BCE; see Supplementary Table 7). In particular the latter individual must be related around the 8$^{th}$ degree to individual I35543 from Tiel Medel. The maximum distance between these individuals can thus be hypothesized to be ~250 years, assuming an average generation time of 25 years. This too is in conflict with an Early Bronze Age date for graves 1, 2, and 3.

Reconciling the archaeological and the genetic evidence is possible, according to a new scenario: The ritual pit must date to the Early Bronze Age, as is attested by the presence of Early Bronze Age sherds, the oldest glass bead found in the Netherlands and a radiocarbon sample from cereal parenchym dated to 2196-1985 cal BCE (GrM-22836: 3702 ± 24 BP; a *terminus post quem*), all found in its lower fill, and it's position in relation to the 'calendar monument' mentioned before. It has been cut through a much older grave, grave 4. After the rituals involving the deposition of animal bones, a glass bead and a skull were completed, the pit was filled and the disturbed remains from grave 4 were reburied in the top of the pit. Contrary to what was initially considered, the (fragments of) late Bell Beaker and Early Bronze Age sherds found directly on top and among the graves do not provide *termini post quem* for the graves. They should instead be regarded as later intrusions. There is evidence for erosion reaching to the top of the human remains. The combination of erosion and trampling could explain their presence.

The cemetery is located on a point bar that was formed around or after 3800 BCE as is attested by two radiocarbon dates from samples found deep in the sediments that were deposited shortly after the channel reactivation. An uncharred fruit of a willow was dated to 3797-3648 cal BCE (Ua-55971: 4959 ± 32 BP) and a charred hazelnut shell was dated to 3945-3656 cal BCE (GrM-17652: 5005 ± 30 BP). They provide *termini post quem* for the formation of the point bar, clearly ruling out an attribution of the graves to the Swifterbant Culture. The oldest radiocarbon samples from Hazendonk period refuse date to 3791-3641 (charcoal alder: GrM-24350 4935 ± 40 BP), 3765-3640 cal BCE (unburnt cattle tibia GrM-22850: 4915 ± 26 BP), 3765-3638 cal BCE (charcoal alder: GrM-24357 4902 ± 29 BP) and 3709-3533 cal BCE (unburnt cattle phalanx: Ua 56120: 4862 ± 30 BP).

The conclusion is that the flat grave cemetery including graves 1, 3 and 4 can be dated to the first half of the 4$^{th}$ millennium BCE. A date around 3700 seems likely.

**Isotopes:** Despite the poor preservation of the skeletons, complete dental elements or enamel fragments from female individuals in Graves 1, 3, and 4 were available for combined Sr-O-C isotope analysis. However, the O-C isotope data should be interpreted with caution (see Supplementary Information S3). The results are presented in Table S2.4.
The $^{87}$Sr/$^{86}$Sr vary significantly among individuals, providing evidence of both intra-individual mobility and diverse regional origins. The Sr isotope signature of Grave 1 is consistent with the expected bioavailable Sr signature for the Tiel region, i.e., the Dutch central river area (Kootker et al., 2016). Her $\delta^{18}O_{PDB}$ values are relatively low, possibly indicating an origin further inland than Tiel. However, the difference between the obtained $\delta^{18}O_{PDB}$ value and the 'local'/'Dutch' lower limit is minimal, particularly when considering potential intra-tooth variation (Plomp et al., 2020). Female Grave 3 exhibits greater variation in $^{87}$Sr/$^{86}$Sr. The first and third molars align with the local Sr isotope signature, as do the $\delta^{18}O_{PDB}$ values, though they fall on the lower end of the expected range. The second molar, however, exhibits a more radiogenic Sr ratio (0.710393), which is inconsistent with the Dutch river area (Kootker et al., 2016) but aligns with regions to the north and south of the site where Pleistocene sandy deposits occur. While she may not have originated far from Tiel, her isotopic data indicate residential mobility during childhood. A scenario in which she left the Tiel De Roeskamp site at an early age and returned a few years later remains plausible.
The most striking results come from female Grave 4. Her $^{87}$Sr/$^{86}$Sr indicate childhood mobility and are incompatible with both the Tiel region and the rest of the Netherlands. While her $\delta^{18}O_{PDB}$ a values fall within the local or Dutch range, her Sr isotope ratios exclude a local origin. Radiogenic Sr signatures of this nature are found in southeastern Belgium (Sengeløv et al., 2025) and Scotland (British Geological Survey materials © UKRI, 2025), though a more northern origin, such as Scandinavia, cannot be ruled out. Given the IBD genetic link between the individual from Grave 4 and Scotland, this represents an intriguing match that warrants further investigation. However, her $\delta^{18}O_{PDB}$ values are too positive for a British origin. Further analysis of the oxygen isotope composition in the phosphate fraction could provide more reliable insights. The $\delta^{13}C_{PDB}$ values suggest that the females' diets during their first 16 years of life were predominantly based on $C_3$ plants, likely sourced from forested or temperate environments.

**Table S2.4.** Sr-O-C isotope data for females 1, 3, and 4 from Tiel Medel De Roeskamp. The element numbers are according to Fédération Dentaire Internationale (FDI).

| Sample ID | Material | Element | $^{87}$Sr/$^{86}$Sr | 2SE | $\delta^{13}C_{PDB}$ (‰) | SD | $\delta^{18}O_{PDB}$ (‰) | SD |
|---|---|---|---|---|---|---|---|---|
| Grave 1_S1147 (I35543) | Enamel | 46 | 0.708727 | 0.000008 | -16.92 | 0.11 | -7.01 | 0.14 |
| | | 47 | 0.708852 | 0.000010 | -14.62 | 0.05 | -6.22 | 0.08 |

| | | | | | | | | |
|---|---|---|---|---|---|---|---|---|
| Grave 3_S2386 (I35544) | Enamel | 46 | 0.708947 | 0.000008 | -15.16 | 0.06 | -6.21 | 0.15 |
| | | 47 | 0.710393 | 0.000007 | - | - | - | - |
| | | 28 | 0.709581 | 0.000008 | -14.00 | 0.17 | -6.42 | 0.20 |
| Grave 4_S1137 (I35542) | Enamel | 16 | 0.715390 | 0.000012 | - | - | - | - |
| | | 17 | 0.715615 | 0.000007 | -15.21 | 0.09 | -6.16 | 0.19 |
| | | 18 | 0.714950 | 0.000008 | -14.76 | 0.04 | -5.52 | 0.09 |

**Source of the samples:** RAAP Archeologisch Adviesbureau. Samples collected by Steffen Baetsen.

**Author of the entry:** Theo ten Anscher, Lisette Kootker

Kootker, L.M., Van Lanen, R.J., Kars, H., Davies, G.R. (2016). Strontium isoscapes in the Netherlands. Spatial variations in $^{87}Sr/^{86}Sr$ as a proxy for palaeomobility, Journal of Archaeological Science: Reports 6, 1-13.

Plomp, E., von Holstein, I.C.C., Kootker, L.M., Verdegaal-Warmerdam, S.J.A., Forouzanfar, T., Davies, G.R. (2020). Strontium, oxygen, and carbon isotope variation in modern human dental enamel, Am J Phys Anthropol 172, 586-604.

Sengeløv, A., Capuzzo, G., Dalle, S., James, H.F., Sabaux, C., Stamataki, E., Hlad, M., Gerritzen, C.T., Legrand, E.M., Veselka, B., Mulder, G.D., Annaert, R., Boudin, M., Salesse, K., Warmenbol, E., Mattielli, N., Snoeck, C., Vercauteren, M. (2025). From plants to patterns: Constructing a comprehensive online strontium isoscape for Belgium (IsoBel) using high density grid mapping, Geoderma 453, 117123. https://doi.org/10.1016/j.geoderma.2024.117123

Ten Anscher, T.J., S. Knippenberg, C.M. van der Linde, W. Roessingh, N. Willemse (2023). Doorbraken aan de Rijn, Een Swifterbant-gehucht, een Hazendonk-nederzetting en erven en graven uit de bronstijd in Medel-De Roeskamp, RAAP-rapport 6519 / Archol rapport 742 / ADC rapport 6150 / BAAC Rapport A-16.0207, RAAP/Archol/ADC ArcheoProjecten/BAAC, Weesp/Leiden/Amersfoort/‘s-Hertogenbosch.

#### 2.5 Schipluiden-Harnaschpolder (04HP) (Zuid-Holland, the Netherlands)

**Analyzed samples:**

| | | |
|---|---|---|
| I38121 | (V9267): | 3550-3500 BCE |
| I38447 | (Grave 5, Ind. 6): | 3630-3500 calBCE (5170±40 BP, GrA-26672) |
| I38448 | (Grave 6, Ind. 7): | 3550-3490 calBCE (5005±40 BP, GrA-26650) |

**Contact information:** Liesbeth Smits

**Site information and excavation history:** The Schipluiden site was located on a submerged sand dune near the coast, of which the top presently is situated at c. 3m minus Dutch Ordnance Datum (NAP). The site was excavated in 2003 by ARCHOL bv. in cooperation with Leiden University in artificial dry conditions (water management). It yielded a wealth of data about Middle Neolithic habitation and economy, including a palisaded settlement area, houses or huts, preservation of organic material and a small number of burials (Louwe Kooijmans & Jongste 2006). Three phases of habitation between 3800 and 3400 BCE were recognized.

**Summary of the sampled materials:** Six burials were found, which contained seven individuals in a flexed or stretched position. Apart from that, a number of 'loose' bones were discovered. The preservation condition was good, probably because originally the dead were buried close to the groundwater table and became submerged soon after habitation (Smits & Louwe Kooijmans 2006: 93). Grave 1 contained the remains of two male individuals, both in extended supine position. Ind. 2 had most likely died from a blow to the front side of the head. For Ind. 1 there were no indications of traumata (Smits & Louwe Kooijmans 2006: 95). Nevertheless, an unnatural death for both individuals may be the reason for the unusual nature of this burial (double and (semi) extended position vs. single and flexed or strongly flexed) (Smits & Louwe Kooijmans 2006: 95).

Individuals 3, 4, 5, 6, and 7 were buried in individual graves in a strongly flexed position, indicative of being tightly wrapped or bound. Individual 6 and 7 (grave 5 and 6 respectively) were young children, Individual 6 yielded sample **I38447,** Individual 7 yielded **sample I38448**. Apart from these six formal burials, the site yielded 36 human bones without burial context, belonging to at least eight individuals (Smits *et al.* 2010). One of these, upper molar **V09267,** yielded sampled (**I38121**).

The strontium isotopes of the individuals (Table S2.5) showed no signs for mobility, but the Oxygen $\delta^{18}O$ levels showed that two of the bones without context (V05001 at 15.8‰ and V08057 at 18.9‰) could not have been local (Smits *et al.* 2010: 20). In their words: "Using the calibration of Daux et al. (2008) shows that SCH6 *(=V05001)* must have spent his childhood somewhere well to the east or south with a drinking water $\delta^{18}O$ of -9.4 ± 0.5‰ , which is consistent with modern precipitation in a broad band from central Scandinavia, through eastern and southern Germany to the western Alps." (Smits *et al.* 2010: 20, 21).

**Table S2.5**: list of sampled individuals (data from Smits et al., 2010; Smits & Louwe Kooijmans, 2006)

| ***Sample ID*** | ***Ind.*** | ***phase*** | ***sex*** | ***age*** | ***Posture*** | ***$^{14}C$*** | ***Calib BCE*** | | $\delta^{13}C$ | $\delta^{15}N$ |
|---|---|---|---|---|---|---|---|---|---|---|
| I38447 | Gr.5 Ind. 6 | 1-2a | inf. | 8y | strongly flexed | 5170 ± 40 (GrA-26672) | 4049-3811 | | -18,50 | |
| I38448 | Gr.6 Ind. 7 | 2 | inf. | 2y | Flexed | 5070 ± 40 (GrA-26737) | 3863-3779 | | -21,05 | 16,36 |
| I38121 | V09267 | 2a | m | 25-35y | manidible + molars | | | | | |

**Dating:** The human bones yield slightly older dates than the proposed period of habitation, but the $\delta^{13}C$ values indicate a reservoir effect of a maximum of 300 years (Mol *et al.* 2006). This effect may have been only very modest to nonexistent for individual 7, who was only 2 years old. From the archaeological evidence it is clear the habitation ended before 3400 BCE, because no Vlaardingen pottery was recorded, only Hazendonk pottery, which was in use until the first half of the fourth millennium BCE (Mol *et al.* 2006: 35; Raemaekers & Rooke 2006).

**Source of the samples:** Provinciaal archeologisch depot Zuid-Holland; Inge Riemersma, Mark Phlippeau. Samples collected by Eveline Altena.

**Authors of entry:** Liesbeth Smits, Harry Fokkens

**References:**

Louwe Kooijmans, L. P., & Jongste, P. F. B. (2006). *Schipluiden. A neolithic settlement on the Dutch North Sea coast, c. 3500 cal BVC* (Vol. 37/38). Faculty of Archaeology.

Mol, J., Louwe Kooijmans, L. P., & Hamburg, T. D. (2006). 2 Stratigrahy and chronology of the site. In L. P. Louwe Kooijmans & P. F. B. Jongste (Eds.), *Schipluiden: A Neolithic Settlement on the Dutch North Sea Coast c. 3500 cal BC* (pp. 19–38). Leiden University Press.

Raemaekers, D. C. M., & Rooke, M. (2006). The Schipluiden pottery. In L. P. Louwe Kooijmans & P. F. B. Jongste (Eds.), *Schipluiden: A Neolithic Settlement on the Dutch North Sea Coast c. 3500 cal BC* (pp. 113–128). Leiden University Press.

Smits, E., & Louwe Kooijmans, L. P. (2006). 5 Graves and human remains. In L. P. Louwe Kooijmans & P. F. B. Jongste (Eds.), *Schipluiden: A Neolithic Settlement on the Dutch North Sea Coast c. 3500 cal BC* (pp. 91–112). Leiden University Press.

Smits, E., Millard, A., Nowell, G., & Pearson, G. (2010). Isotopic Investigation of Diet and Residential Mobility in the Neolithic of the Lower Rhine Basin. *European Journal of Archaeology*, *13*, 5–31. https://doi.org/10.1177/1461957109355040

#### 2.6a Molenaarsgraaf (Zuid-Holland, the Netherlands)

**Analyzed samples:**

| | | |
|---|---|---|
| I13025 | (h 1967/1._Skelet I; grave I): | 2136-1892 calBCE (3635±40 BP, GrN-5131) |
| I13026 | (h 1967/1._Skelet II; grave II): | 2135-1890 calBCE (3630±40 BP, GrN-5566) |
| I13027 | (h 1967/1._Skelet III; grave III): | 2197-1983 calBCE (3700±25 BP, PSUAMS-7847) |

These samples have previously been reported in Patterson, N., Isakov, M., Booth, T. et al. Large-scale migration into Britain during the Middle to Late Bronze Age. Nature 601, 588–594 (2022). https://doi.org/10.1038/s41586-021-04287-4.

#### 2.6b Molenaarsgraaf_24A (Zuid-Holland, the Netherlands)

**Analyzed sample:**

| | | |
|---|---|---|
| I12896 | (24A: h 1973/3_18,19): | 2864-2500 calBCE (4100±30 BP, PSUAMS-7806) |

**Contact information:** Luc Amkreutz, Leendert Louwe Kooijmans

**Site information and excavation history:** In August 1972 local archaeologists discovered human bones in the side of a newly cleaned ditch. They reported this to the keeper of the National Museum of Antiquities (L.P. Louwe Kooijmans). On a Saturday afternoon with two technicians of the museum, Louwe Kooijmans rescued the remains as well as possible. They documented two concentrations of human bone c. 60 cm below the surface, situated on the sediments of the Schoonrewoerd stream ridge. There was no (grave) pit visible, and the bones were reported not to be in anatomical position (museum diary for find number 1973/3), even though Louwe Kooijmans remembers a person in crouched position. A few flints were recovered from the site as well, no pottery. The site was never published in detail but mentioned as site 24a by Louwe Kooijmans in his dissertation (Louwe Kooijmans 1974).

**Summary of the sampled materials:** The material of site 24A was inventoried and analyzed by Hilde Uytterschaut (unpublished document), who suggests that 3 individuals were present in this collection of bones, mainly based on the dental elements. There was 1 adult at least and two children (4-7 y. old). A tooth from the upper jaw of one adult individual of 30-40 years old (RM1973/18-19) yielded sample **I12896**.

**Dating:** The sample of I12896 is dated to 2864-2500 calBCE (4100¬±30 BP, PSUAMS-7806). This suggests a late Vlaardingen/Corded Ware context. The $\delta^{13}C$ value is -22.0 ‰ and $\delta^{15}N$ value 14.2 ‰. This could imply that a reservoir effect is present.

**Source of samples**: National Museum of Antiquities Leiden; Luc Amkreutz. Sample collected by Eveline Altena.

**Authors of entry**: Luc Amkreutz, Harry Fokkens, Leendert Louwe Kooijmans

**References**:

Louwe Kooijmans, L.P. (1974). *The Rhine/Meuse Delta; four studies on its prehistoric occupation and Holocene geology*. Leiden: Instituut voor Prehistorie.

#### 2.7 Mienakker (OPM'90) (Noord-Holland, the Netherlands)

**Analyzed samples:**

| | |
|---|---|
| I12902 (R9560-11_V2764, S54): | 2852-2574 calBCE (4100±20 BP, PSUAMS-8438); 2554-2202 calBCE (3890±50 BP, GrA-1670) |

**Contact information:** Harry Fokkens

**Site information and excavation history:** The site of Mienakker was excavated in 1990 but was published only in 2013 by a multidisciplinary team of Groningen University, Leiden University and the Cultural Heritage Agency of the Netherlands (Kleijne *et al.* 2013). This team re-analysed all documentary and material culture evidence but was not involved in the original excavation. The spatial analysis of the site was carried out by Nobles. He describes the grave of OPM90, nicknamed by the excavators as ‘Cees’, as

deposited in a pit that was dug into the habitation layer of the site (Nobles 2013, 36; Figure 1). There were no grave goods. Stratigraphically the burial is interpreted as one of the most recent elements at the site.
In his spatial analysis Nobles reconstructs a house-like ritual structure around this grave (Nobles 2013b, esp. 238 ff.). However, there are serious doubts about that reconstruction (Fokkens et al. 2016, 76). Nobles has tried to settle the debate (Nobles 2020), but not convincingly in our view.

**Summary of the sampled materials:** Skeleton OPM'90/S54 yielded **sample I12902**; he was 20-25 years old, may have been about 1.72 m long and there were no tell-tale signs of illnesses (Pasveer et al. 1992, 270). The pit in which OPM'90 was buried, was only 120 cm long and 96 cm wide. The bottom was 25 cm below the excavation surface (Nobles 2013a, 33). Because the skeleton was partly in anatomical position, especially with respect to the labile joints of the left hand, Plomp concludes that it was probably a primary burial (Plomp 2013, 176). Strangely enough, both legs and the right arm were missing. The original examiners of the skeleton, Pasveer and Uytterschout, state that gnaw-marks were present of a dog or wolf especially on the knee joints (femur and tibia) and on the pelvis (Pasveer & Uytterschaut 1992: 274; Plomp 2013: 178). Moreover, the clavicle was broken, but no gnaw marks were reported. In other words: the right arm was (forcefully?) removed, and the lower legs were bitten off. According to Plomp this happened not very long after death because the labile hand joints were still in anatomical position (Plomp 2013, 176). Given the 'functional' dimensions of the grave pit (just large enough to fit the torso), the possibility remains that this was the grave of a person who died elsewhere and the body was exposed to open air for a short time , accessible to scavengers.

**Dating:** From the stratigraphy of the site, it is clear that the grave '…was dug in the final stages of human activity ...' (Nobles 2013a, 33). Kleijne and Weerts distinguish two possible phases of habitation, the first starting in Furholts phase D (2880-2680 BCE; (Furholt 2003; Kleijne et al. 2013), the second in Furholts phase E (2620-2480 BCE; (Furholt 2003; Kleijne et al. 2013)). The grave was dug in the second phase. The skeleton was direct dated between 2676 and 2433 cal BC (GrA-15698: 4010 +/- 50 BP), but the collagen content was considered poor. An earlier sample date, also of bone collagen (GrA-1670 3890±50 BP), was rejected because that and others from the same batch appeared too young (Lanting et al. 2002, 76–77). The $\delta^{13}C$ and $\delta^{15}N$ isotope readings are -18.4 ‰ and 15.5‰ respectively, so a (mild) reservoir effect could be present (Kleijne et al. 2013, 25–26).
**Isotopes:** 'Cees' is currently part of the permanent exhibition at the museum that is part of the *Provinciaal Depot van Noord-Holland* in Castricum. To ensure minimally invasive sampling, three molars from both the maxilla and mandible were collected *in situ*. The results of the Sr-O-C isotope analysis are presented in Table S2.7.
The Sr and O isotope data align with the expected values for the Mienakker region (West-Friesland) and the broader Netherlands, respectively (Kootker et al., 2016; Kootker et al., 2019). This suggests that 'Cees' possibly spent the first 16 years of life in a single location, possibly within West Friesland.
The $\delta^{13}C$ values show some variation (–12 ‰ to –10 ‰). The –10 ‰ value is relatively high compared to other data from the Netherlands for this period. Such elevated values could be attributed to the consumption of $C_4$ plants or fish. Given that a potential reservoir effect has also been detected in the bone collagen, it is plausible that fish consumption, possibly for a limited period during childhood, contributed to this more positive $\delta^{13}C$ signature.

**Table S2.7.** Sr-O-C isotope data for 'Cees' (S54) from Mienakker. The element numbers are according to Fédération Dentaire Internationale (FDI).

| Sample ID | Material | Element | $^{87}Sr/^{86}Sr$ | 2SE | $\delta^{13}C_{PDB}$ (‰) | SD | $\delta^{18}O_{PDB}$ (‰) | SD |
|---|---|---|---|---|---|---|---|---|
| OPM'90/S54/'Cees' (I12902) | Enamel | 26 | 0.709148 | 0.000006 | -12.57 | 0.09 | -5.44 | 0.15 |
| | | 47 | 0.709181 | 0.000006 | -10.89 | 0.08 | -5.79 | 0.10 |
| | | 48 | 0.709230 | 0.000008 | -12.11 | 0.06 | -5.53 | 0.15 |

**Source of samples:** Provinciaal depot voor archeologie van Noord-Holland; Martin Veen, Rob van Eerden. Samples collected by Eveline Altena

**Author of entry:** Harry Fokkens, Eveline Altena, Lisette Kootker

**References:**

Fokkens, H., Steffens, B.J.W., & van As, S.F.M. (2016). *Farmers, fishers, fowlers, hunters. Knowledge generated by development-led archaeology about the Late Neolithic, the Early Bronze Age and the start of the Middle Bronze Age (2850 - 1500 cal BC) in the Netherlands.* Amersfoort: Rijksdienst voor het Cultureel Erfgoed.

Furholt, M. (2003). *Die absolutchronologische Datierung der Schnurkeramik in Mitteleuropa und Südskandinavien*. Bonn: Dr. Rudolf Habelt GMBH.

Kleijne, J.P., O. Brinkkemper, R.C.G.M. Lauwerier, B.I. Smit & E.M. Theunissen. 2013. *A Matter of Life and Death at Mienakker (the Netherlands). Late Neolithic Behavioural Variability in a Dynamic Landscape*. Vol. 45 (Nederlandse Archeologische Rapporten 45). Amersfoort: Cultural Heritage Agency of the Netherlands.

Kleijne, J.P., & Weerts, H.J.T. (2013). 2 Landscape and chronology. In J. P. Kleijne, O. Brinkkemper, R. C. G. M. Lauwerier, B. I. Smit, & E. M. Theunissen (eds) *A Matter of Life and Death at Mienakker (the Netherlands). Late Neolithic Behavioural Variability in a Dynamic Landscape.*Nederlandse Archeologische Rapporten, 19–28. Amersfoort: Cultural Heritage Agency of the Netherlands

Kootker, L.M., Van Lanen, R.J., Kars, H., Davies, G.R. (2016). Strontium isoscapes in the Netherlands. Spatial variations in $^{87}Sr/^{86}Sr$ as a proxy for palaeomobility, Journal of Archaeological Science: Reports 6, 1-13. 10.1016/j.jasrep.2016.01.015

Kootker, L.M., van Lanen, R.J., Groenewoudt, B.J., Altena, E., Panhuysen, R.G.A.M., Jansma, E., Kars, H., Davies, G.R. (2019). Beyond isolation: understanding past human-population variability in the Dutch town of Oldenzaal through the origin of its inhabitants and its infrastructural connections, Archaeological and Anthropological Sciences 11, 755-775. 10.1007/s12520-017-0565-7

Lanting, J.N., & van der Plicht, J. (2002). De $^{14}$C Chronologie van de Nederlandse Pre- en Protohistorie III: Neolithicum. *Palaeohistoria* 41/42 (1999-2000): p.1–110.

Nobles, G.R. (2013a). 3 Features. In J. P. Kleijne, O. Brinkkemper, R. C. G. M. Lauwerier, B. I. Smit, & E. M. Theunissen (eds) *A Matter of Life and Death at Mienakker (the Netherlands). Late Neolithic Behavioural Variability in a Dynamic Landscape*.Nederlandse Archeologische Rapporten 45, 29–36. Amersfoort: Cultural Heritage Agency of the Netherlands

Nobles, G.R. (2013b). 11 Spatial analysis. In J. P. Kleijne, O. Brinkkemper, R. C. G. M. Lauwerier, B. I. Smit, & E. M. Theunissen (eds) *A Matter of Life and Death at Mienakker (the Netherlands). Late Neolithic Behavioural Variability in a Dynamic Landscape*.Nederlandse Archeologische Rapporten 45, 185–240. Amersfoort: Cultural Heritage Agency of the Netherlands

Nobles, G.R. (2020). Settling the monumental issue in the Dutch Wetlands. In A. B. Gebauer, L. Sørensen, A. Teather, & A. C. Valera (eds) *Monumentalising Life in the Neolithic: Narratives of Continuity and Change*, 125–138. Oxford: Oxbow Books Available at: https://doi.org/10.2307/j.ctv13pk66m.16.

Pasveer, J.M., & Uytterschaut, H.T. (1992). Twee Laat-Neolithische skeletten uit Noord-Holland, een fysisch-anthropologisch onderzoek. *Westerheem* 41: p.268–275.

Plomp, E. (2013). 10 The human skeleton. In J. P. Kleijne, O. Brinkkemper, R. C. G. M. Lauwerier, B. I. Smit, & E. M. Theunissen (eds) *A Matter of Life and Death at Mienakker (the Netherlands). Late Neolithic Behavioural Variability in a Dynamic Landscape*. Nederlandse Archeologische Rapporten 45, 175–184. Amersfoort: Cultural Heritage Agency of the Netherlands

#### 2.8 Sijbekarspel-Op de Veken (Noord-Holland, the Netherlands)

**Analyzed samples:**

I33741 ( HvH 8077-01): 2571-2468 calBCE (4015±20 BP) [R_Combine: (4015±20 BP, PSUAMS-12657), (3890±50 BP, GrA-15696)

**Contact information:** Harry Fokkens

**Site information and excavation history:** The site Op de Veken in Sijbekarspel was discovered through corings in 1986 and surveyed in more detail in 1987. In that year also four 1x2m test pits were dug. One of these was extended to 4x4m when a grave was discovered (van Heeringen & Theunissen, 2001). The site was preliminary published by Hogestijn (Hogestijn & Woltering, 1990), and a detailed analysis of the skeleton was performed (Pasveer & Uytterschaut, 1992b, 1992a). As most of the skeletons found in West-Frisia were given names by their excavators, the Sijbekarspel skeleton was nicknamed 'Mies, het woiffie van Soibekarspel' (Hogestijn & Woltering, 1990). IBD analysis (Supplementary Table 7) made clear that I33471 shared a distant kinship (apprximately $8^{th}$ degree) with indidividual I12902 from Mienakker.

**Summary of the sampled materials:** The Sijbekarspel skeleton yielded **sample I33741**; she was a woman aged about 30-35. She was placed on her left side facing south in a crouched position. In her youth she probably had severe health problems, a.o. a shortage of nutrients in her food (Pasveer & Uytterschaut, 1992a, p. 271). Her teeth had remains of dental plaque showing a diet of grain, and even diatoms from a salt to brackish environment, consistent with the West-Frisian wetland landscape at the time (Pasveer & Uytterschaut, 1992a, p. 273).

**Dating:** A first AMS-date of bone collagen yielded an unexpectedly young date: GrA-1644 3550±50 BP. Since this date belonged to a small series with strange results, it was re-dated (Lanting & van der Plicht, 2002, p. 76). This time it yielded 3890±50 BP (GrA-15696; 2475-2202 cal BCE). The $\delta^{13}C$ value of –20. 0 ‰ and the $\delta^{15}N$ value of 14.1 ‰ does not suggest a reservoir effect (Lanting & van der Plicht, 2002, p. 76). Charred hazelnut shells from the site itself yielded a slightly older date: 3960+-60 BP (GrA-107) (van Heeringen & Theunissen, 2001). The stratigraphy of the site places the grave in the Vlaardingen/Corded Ware period of use, most likely slightly after 2500 BCE.

**Isotopes:** Comparable to her distant kin from Mienakker, Mies' $^{87}Sr/^{86}Sr$ aligns with the expected bioavailable local strontium signature (Kootker et al. 2016, Table S2.8). The inter-dental elemental variation in $^{87}Sr/^{86}Sr$ is negligible, as is the case for $\delta^{13}C$ values, which indicate a diet primarily based on $C_3$ plants. In contrast, the $\delta^{18}O$ data exhibit significant variation. The $M^1$ displays the most enriched $\delta^{18}O$ value, which may be an artefact of breastfeeding. While this pattern, where earlier mineralizing dental elements show less negative $\delta^{18}O$ values, does not occur in all individuals, it remains a plausible explanation for the observed variation. The $\delta^{18}O$ values of the $M^2$ and $M^3$ are relatively low, even lower than those of 'Cees'. Although the $\delta^{18}O$ of the $M^2$ exceeds the defined lower limit, the values may still be compatible with the isotopic range of "West-Friesland". However, further research is necessary to assess the reliability of these data before drawing more definitive conclusions.

**Table S2.8.** Sr-O-C isotope data for 'Mies' (S54) from Sijbekarspel. The element numbers are according to Fédération Dentaire Internationale (FDI).

| Sample ID | Material | Element | $^{87}Sr/^{86}Sr$ | 2SE | $\delta^{13}C_{PDB}$ (‰) | SD | $\delta^{18}O_{PDB}$ (‰) | SD |
|---|---|---|---|---|---|---|---|---|
| 'Mies/Woiffie'/ 8077-01 (I33741) | Enamel | 16 | 0.709075 | 0.000008 | -12.97 | 0.04 | -4.85 | 0.05 |
| | | 17 | 0.709181 | 0.000008 | -12.24 | 0.03 | -7.69 | 0.15 |
| | | 18 | 0.709173 | 0.000006 | -12.83 | 0.08 | -6.42 | 0.11 |

**Source of samples:** Provinciaal depot voor archeologie van Noord-Holland; Martin Veen, Rob van Eerden. Sample collected by Eveline Altena.

**Author of entry:** Harry Fokkens, Lisette Kootker

**References:**

Hogestijn, J. W. H., & Woltering, P. J. (1990). 'Het woiffie van Soibekarspel': Een Laat-Neolithisch vrouwengraf te Sijbekarspel. *West-Frieslands Oud En Nieuw*, *57*, 152–164.

Lanting, J. N., & van der Plicht, J. (2002). De $^{14}$C Chronologie van de Nederlandse Pre- en Protohistorie III: Neolithicum. *Palaeohistoria*, *41/42 (1999-2000)*, 1–110.

Pasveer, J. M., & Uytterschaut, H. T. (1992a). Twee Laat-Neolithische skeletten uit Noord-Holland, een fysisch-anthropologisch onderzoek. *Westerheem*, *41*, 268–275.

Pasveer, J. M., & Uytterschaut, H. T. (1992b). Two Late Neolithic human skeletons, a recent discovery in the Netherlands. *International Journal of Osteoarchaeology*, *2*(1), 1–14. https://doi.org/10.1002/oa.1390020102

van Heeringen, R. M., & Theunissen, E. M. (2001). *Kwaliteitsbepalend onderzoek ten behoeve van duurzaak behoud van neolithische terreinen in West-Friesland en de Kop van Noord-Holland* (Vol. 21). ROB.

#### 2.9 Oostwoud-Tuithoorn (Noord-Holland, the Netherlands)

**Analyzed samples:**

I39211 (skeleton 239): 1981-1692 cal BCE (3520±50 BP, GrA-15601)

I39210 (Skeleton S233-A16-005_M125) (contaminated)

**Contact information:** Harry Fokkens

**Site information and excavation history:** In 1956 and 1957, Prof. van Giffen excavated one Late Neolithic and one Early Bronze Age burial mound at Oostwoud-Tuithoorn (van Giffen 1961, 1962). Since van Giffen had not been able to finish the excavations, additional research was conducted in 1963 and 1966 by de Weerd (de Weerd 1963, 1967). Finally in 1977 and 1978, when the site was being threatened by deep-ploughing, a large-scale excavation was conducted under supervision of van der Waals and Lanting (Lanting 1979, 2008). During these excavations a total of 15 well preserved skeletons were recovered, dating to three episodes of activity between c. 2340–1780 cal BCE. A detailed account of all excavations was published by

Fokkens et al. (Fokkens *et al.* 2017), and aDNA samples of other skeletons were published earlier by Olalde et al. (Olalde *et al.* 2018) and Patterson *et al.* 2021.

**Summary of the sampled materials:** For the present study an additional two individuals from the site were sampled: skeleton S233 and S239.

Skeleton 233 was buried in a pit which was disturbed in the Middle Ages, leaving less than 25 % of the skeleton (the skull and part of the pelvis: Veselka 2016). Its orientation was not properly recorded in the field drawings, but a sketch shows a crouched position on the right side with the head pointing south, facing east (Lanting and van der Plicht 2002, 86). That position is in line with that of other female skeletons at Oostwoud, though these were all facing north. Osteoarchaeological analysis considered the individual to be a man of 36-49 years old (Veselka 2016), though earlier analysis had suggested this to be possibly female (Runia 1987). S**ample I39210** of skeleton 233 was suspected of contamination and not used in the analysis (Supplementary table 2), but indicated the molecular sex as female.

Skeleton 239 yielded **sample I39211**. This skeleton was of a relatively tall male individual (181.4 ± 3.27 cm) who was buried in extreme flexed position, probably because he was wrapped in a mat of some kind (Fokkens et al. 2017, 141).

**Dating:** Skeleton 233 (I39210) was not direct dated, but she probably should be placed in the same episode of burial as S242 (I4074), S236 (I4073) and S228 who were related in the second or third degree (Supplementary table 7); Fokkens *et al.* 2017). These were buried between 2140 and 2070 BCE.

Skeleton 239 (I39211) was dated to the Early Bronze Age (GrA-15601: 3520 +/1 60 BP; 1981-1662 calBCE), though still buried in a Late Neolithic tradition in terms of position and orientation. Therefore, we suggest he dates to Episode 4, between 1900 and 1800 BCE.

**Source of samples:** Provinciaal depot voor archeologie van Noord-Holland; Martin Veen, Rob van Eerden. Samples collected by Eveline Altena.

**Author of entry:** Harry Fokkens

**References:**

DE WEERD, M.D. 1963. Protocolboek opgraving Oostwoud. Instituut voor Prae- en Protohistorie, Universiteit van Amsterdam.

DE WEERD, M.D. 1967. Medemblik [nabij Oostwoud]. *Nieuwsbulletin KNOB* 1967, 2e afl. februari, kolom *31-*32.

Fokkens, H., Veselka, B., Bourgeois, Q., Olalde, I., & Reich, D. (2017). Excavations of Late Neolithic arable, burial mounds and a number of well-preserved skeletons at Oostwoud-Tuithoorn; a re-analysis of old data. *Analecta Praehistorica Leidensia*, *47*, 95–150.

Lanting, J. N. (1979). Medemblik: Oostwoud. *Archeologische Kroniek van Noord-Holland over 1978*, 250–251.

Lanting, J. N. (2008). De NO-Nederlandse/NW-Duitse Klokbekergroep: Culturele achtergrond, typologie van het aardewerk, datering, verspreiding en grafritueel. *Palaeohistoria* 49/50 (2007-2008): 11–326.

Olalde, I. et al. (2018). The Beaker phenomenon and the genomic transformation of northwest Europe. *Nature* 555: 190–96. https://doi.org/10.1038/nature25738 https://www.nature.com/articles/nature25738#supplementary-information.

Patterson, N. *et al*. (2021). Large-scale migration into Britain during the Middle to Late Bronze Age. *Nature* 601: 588–94. https://doi.org/10.1038/s41586-021-04287-4.

Runia, L.T. (1987). *The chemical analysis of prehistoric bones. A paleodietary and ecoarcheological study of Bronze Age West-Friesland* (British Archaeological Reports International Series 363). Oxford: BAR publishing.

Van Giffen, A.E. (1961). Settlement traces of the Early Bell Beaker Culture at Oostwoud (N.H.). *Helinium* 1: 233–228.

Van Giffen, A.E. (1962). Grafheuvels uit de midden-bronstijd met nederzettingssporen van de Klokbekercultuur bij Oostwoud. *West-Frieslands Oud en Nieuw* 29: 199–209.

Veselka, B. (2016). Fysisch antropologische analyse van het menselijk skeletmateriaal uit Oostwoud. Leiden.

#### 2.10 Ottoland-Kromme Elleboog (Zuid-Holland, the Netherlands)

**Analyzed samples:**

| | | |
|---|---|---|
| I12900 | (h 1982/7._4_Skelet II) | 2457-2145 calBCE (3820±45 BP, GrN-6384) |
| I13028 | (h 1982/7._4_Skelet I) | 2500-2100 BCE |

These two individuals have previously been reported in Patterson, N., Isakov, M., Booth, T. et al. Large-scale migration into Britain during the Middle to Late Bronze Age. Nature 601, 588–594 (2022). https://doi.org/10.1038/s41586-021-04287-4. We produced additional genetic data for I12900.

#### 2.11 Doggerland (the Netherlands)

**Used samples:**

| | | |
|---|---|---|
| DOG007 | (U 2014/12.3;) | 7576-7201 calBCE (8370±50 BP, GrA-11642) |
| DOG001 | (A10-007) | 7730-7586 calBCE (8627±35 BP, MAMS-48201) |
| DOG002 | (U 2014/12.4) | 8421-8238 calBCE (9091±37 BP, MAMS-34582) |

A detailed archaeological context was published in Posth, C., Yu, H., Ghalichi, A., Rougier, H., Crevecoeur, I., Huang, Y., Ringbauer, H., Rohrlach, A. B., Nägele, K., Villalba-Mouco, V., Radzeviciute, R., Ferraz, T., Stoessel, A., Tukhbatova, R., Drucker, D. G., Lari, M., Modi, A., Vai, S., Saupe, T., … Krause, J. (2023). Palaeogenomics of Upper Palaeolithic to Neolithic European hunter-gatherers. *Nature*, *615*(7950), 117–126. https://doi.org/10.1038/s41586-023-05726-0

#### 2.12 Swifterbant (S2) (Dronten, Flevoland, the Netherlands)

**Analyzed individuals:**

| | | |
|---|---|---|
| SWA001 | (Skelet II): | 4180-4030 BCE |
| SWA002 | (Skelet III): | 4180-4030 BCE |
| SWA004 | (Skelet IV): | 4180-4030 BCE |

**Contact person:** Daan Raemaekers

**Site information and excavation history:**

The archaeological sites near Swifterbant were discovered in the early 1960's as a result of the creation of the polder *Oostelijk Flevoland*. There are two landscapes in which sites were found. The first is that of sandy ridges that were located along the small river Hunnepe. These sand ridges were occupied during the Mesolithic and Neolithic and have in general a poor preservation condition. Nevertheless, 13 human burials were documented at location S21-S24 (Meiklejohn & Constandse-Westermann 1979), of which 12 were $^{14}$C dated, between 4600-4000 calBCE (Raemaekers et al. 2014). Visual inspection of these remains by E. Altena resulted in their exclusion from the current analysis. The second landscape with archaeological sites is that of the levees that accompanied the Hunnepe river. Here, the sites S2, S3 and S4 are of relevance.

The sites are located on one small river system. S3 and S4 are within meters distance from one another, while S2 is located at some 500 m distance. Until recently, the sites were dated c. 4300-4300 calBCE, a relatively unprecise date because of the presence of a plateau in the calibration curve. Now, the chronology of S3 and S4 has been re-analyzed, making use of short-lived samples of cereal grains, high-precision AMS dating and Bayesian modelling. As a result, S4 is now dated 4240-4160 calBCE and S3 is now dated 4180-4030 BCE (Dreshaj et al. 2024).

S2 was not included in this re-analysis. What can be said about the date of the site and the human burials that were documented? The single $^{14}C$ date from the settlement is on charcoal (4252-3995 calBCE: GrN-5443. 5300 ± 40 BP). Because this date fits the S3 dates perfectly and S2 is located on the same river system, we propose that the site of S2 dates between 4180-4030 BCE as well. During fieldwork, the contours of the burial pits could be seen to have been dug through the settlement layer, implying a younger date than the settlement activities.

Detailed subsistence data are available for all three sites. They make clear that the Swifterbant occupants practiced hunting and husbandry, gathering and cultivation. Here, we focus on those elements that underline that these sites should be interpreted as Early Neolithic, rather than Late Mesolithic. Evidence for cereal cultivation is abundant. At all three sites, cultivated fields were documented (Huisman & Raemaekers 2014), charred cereal remains were found in large numbers (Van Zeist & Palfenier-Vegter 1981; Schepers 2020), many ceramic vessels were present (Raemaekers et al.2014), and all coprolites (Kubiak-Martens and Van der Linden 2022) yielded microscopic or chemical evidence of cereal remains. While most mammal bones at these sites come from pigs, the domestic status remains ambiguous. Cattle bones are found in smaller numbers and testify to full control over fodder, mobility (Brusgaard et al. 2024) and reproduction (Erven et al. in prep.). This indicates the social importance of domestic cattle to these people.

**Summary of the sampled materials:**

The human remains from S2 comprise ten burials in nine burial pits. The burials share an NNW-SSE orientation and are found on the eastern levee of the Hunnepe river. The age and gender distributions are as follows (Meiklejohn & Constandse-Westermann 1979):

| | | |
|---|---|---|
| Burial II (sample SWA001) | Female (probable) | 35+ years |
| Burial III (sample SWA002) | Male (probable) | 20-35 years |
| Burial IV (sample SWA003) | Male | 20-55 years |

**Source of the samples:** Provinciaal Archeologisch Depot Flevoland; Tineke Heise-Roovers. Samples collected by Eveline Altena.

**Author of the entry**: Daan Raemaekers

**References:**

Brusgaard, N. Ø., Kooistra, J., Schepers, M., Dee, M., Raemaekers, D., & Çakırlar, C. (2024). Early animal management in northern Europe: Multi-proxy evidence from Swifterbant, the Netherlands. *Antiquity*, *98*(399), 654–671. https://doi.org/10.15184/aqy.2024.58

Meiklejohn, C., & Constandse-Westerman, T. S. (1978). The human skeletal material from swifterbant, Earlier Neolithic of the northern Netherlands: I. Inventory and demography. *Palaeohistoria*, 39–89.

Dreshaj, M., D.C.M. Raemaekers & M. Dee, (2024). Chronological modelling on a calibration plateau: implications for the emergence of agriculture in the Dutch wetlands, *Radiocarbon* 65(6), 1280-1298. https://doi.org/10.1017/RDC.2023.126

Huisman, D.J. & D.C.M. Raemaekers, (2014). Systematic cultivation of the Swifterbant wetlands (The Netherlands). Evidence from Neolithic tillage marks (c. 4300–4000 cal. BC), *Journal of Archaeological Science* 49, 572-584.

Kubiak-Martens, L. & Van der Linden, M. (eds) (2022). *Neolithic Human Diet. Based on Studies of Coprolites from the Swifterbant Culture Sites, the Netherlands*. Nederlandse Archeologische Rapporten 77. Amersfoort: Cultural Heritage Agency of the Netherlands.

Raemaekers, D.C.M., J. Geuverink, I. Woltinge, J. van der Laan, A. Maurer, E.E. Scheele, T. Sibma & D.J. Huisman, (2014). Swifterbant-S25 (gemeente Dronten, provincie Flevoland). Een bijzondere vindplaats van de Swifterbant-cultuur (ca. 4500-3700 cal. BC), *Palaeohistoria* 55/56, 1-56.

Schepers, M. & N. Bottema-Mac Gillavry, (2020). The vegetation and the exploitation of plant resources. In D.C.M. Raemaekers & J.P. de Roever (red.), *Swifterbant S4 (the Netherlands). Occupation and exploitation of a Neolithic levee site (c. 4300-4000 cal. BC),* Groningen (Groningen Archaeological Studies 36),51-75.

Van Zeist, W. , & R.M. Palfenier-Vegter, (1981). Seeds and fruits from the Swifterbant S3 site. Final Reports on Swifterbant IV, *Palaeohistoria* 23, 105-168.

### Belgium

#### 2.13 Abri des Autours (Dinant, Namur, Belgium)

**Used sample:**

| | | |
|---|---|---|
| AAT001 | (AA3): | 9160-8623 calBCE (9500±75 BP, OxA-4917) |

A detailed archaeological context was published in: Semal, P., Polet, N., & Cauwe, N. (2023). Abri de Autours, Belgium Supplementary information; section 1 archaeological contact information to Posth et al. 2023. *Nature*, *615*(7950). https://doi.org/10.1038/s41586-023-05726-0

#### 2.14 Malonne Petit Ri (Namur, Belgium)

**Used sample:**

| | | |
|---|---|---|
| MPR001 | (MPR-1): | 8731-8294 calBCE (9270±90 BP, OxA-5042) |

A detailed archaeological context was published in: Semal, P., & Jadin, I. (2023). Malonne Petit Ri, Belgium Supplementary information; section 1 archaeological contact information to Posth et al. 2023. *Nature*, *615*(7950). https://doi.org/10.1038/s41586-023-05726-0

#### 2.15 Trou Al'Wesse (Modave, Liège, Belgium)

**Analyzed sample:**

I13627 (1 / x4; AF004): 3334-3028 calBCE (4466±21 BP, OxA-39060)

I13642. (2 / 3120; AF018) 3500-3100 BCE (contaminated)
I13648 (4 / 3262; AF024) 3500-3100 BCE (contaminated)

**Contact information:** Maria Pala, John Stewart

**Site information and excavation history:**

Trou Al'Wesse ('Wasp Cave' in the Walloon dialect) is part of a limestone karstic cave system located in the Hoyoux valley. The cave extends almost horizontally for 35m inside a cliff. At the back, the ceiling opens through a 9m long chimney that connects to the plateau above. The cave has been known since excavations first occurred in the 1860s, and it has been the object of numerous investigations since (Masy, 2020-2022, Miller et al., 2011, Flas et al., 2019). The Pleistocene and Holocene deposits show different phases of human occupation from the Middle and Upper Palaeolithic, Mesolithic and Neolithic.

**Summary of the sampled materials:** The human remains included in this study were retrieved from the deposits filling the chimney at the back of the cave, and were excavated in the 1880s (Masy, 1993)

**Dating:** Previous radiocarbon dates on two human 5$^{th}$ left metatarsals established a Neolithic chronology for some of these remains: 4560±30 (Beta-319269 / 5436–5052 cal. BP) and 4450±30 (Beta-319270 / 5284–4885 cal. BP) (Miller *et al*., 2012).

Genetic Identifier: I13627. Grave Identifier: 1 / x4; AF004. Grave type: chimney burial. Skeletal information: left petrous. Grave goods: none. Dating: radiocarbon date 3334-3028 calBCE (4466±21 BP, OxA-39060; skeletal element: left petrous). Additional information: collective 'grave' excavated during the late 19th century.

The following two samples were not used in the analysis because of suspected contamination (Supplementary Table 2)

- Genetic Identifier: I13642. Grave Identifier: 2 / 3120; AF018. Grave type: chimney burial. Skeletal information: left humerus. Grave goods: none. Dating: context date 3500-3100 BCE. Additional information: collective 'grave' excavated during the late 19th century.

- Genetic Identifier: I13648. Grave Identifier: 4 / 3262; AF024. Grave type: chimney burial. Skeletal information: right tibia. Grave goods: none. Dating: context date 3500-3100 BCE. Additional information: collective 'grave' excavated during the late 19th century.

**Source of the samples:** University of Huddersfield

**Authors of the entry:** Maria Pala, Alessandro Fichera

**References**:

Flas D., Zwyns N., Stewart J., Wilkinson K., Barrett N., Knul M. & Noiret P. (2019). Modave/Modave: Trou Al'Wesse, fouilles 2018. *Chronique de l'Archaeologie Wallonne* 27, 161-164.

Masy P. (1993). La sépulture collective néolithique du trou Al'Wesse à Modave (province de Liège). *Bulletin des Chercheurs de la Wallonie***,** XXXIII, 81-99.

Masy P. (2020-2022). Historique du Trou Al'Wesse (Modave) avant les fouilles modernes commencées en 1988. *Bulletin des Chercheurs de la Wallonie* 55**,** 5-23.

Miller R., Collin F., Otte M. & Stewart J. (2011). Le Trou Al'Wesse: du Moustérien au Néolithique dans la vallée du Hoyoux. *Le Paléolithique moyen en Belgique. Mélanges Marguerite Ulrix-Closset.* Liège: ERAUL 128.

Miller R., Stassart E., Otte M., Austin P. & Stewart J. (2012). Interprétation chronostratigraphique de la séquence holocène du Trou Al'Wesse à la lumière des nouvelles datations: du Mésolithique ancien au Néolithique moyen (Modave, B). *Notae Praehistoricae* 32**,** 133-139.

#### 2.16 Grotte du Mont Falise (Liège, Belgium)

**Analyzed samples:**

| | | |
|---|---|---|
| I13629 | (213 / 3259; AF006): | 2950-2600 BCE |
| I13631 | (213 / 15.019); AF008 | 2950-2600 BCE |
| I13638 | (213 / 3252); AF014 | 2886-2668 calBCE (4180±25 BP, PSUAMS-12142) |
| I13649 | (212 / 1x.021); AF026 | 3050-2600 BCE |
| I13651 | (212 / 3245); AF028 | 3050-2600 BCE |

**Contact information:** Maria Pala, Damien Flas, Pierre Noiret

**Site information and excavation history:**

The Grotte du Mont Falise, is a cave located near Antheit (Liege province). It opens into a limestone cliff in the basin of the Mehaigne. The cave was excavated in the 1890s by Julien Fraipont, and in 1958 by Haeck (Haeck 1964, Fraipont 1897). Apart from human remains, the site yielded archaeological material from the Middle and Upper Palaeolithic, Neolithic, Iron Age, Roman, and Medieval periods (Haeck 1964).

**Summary of the sampled materials:** Genetic Identifier: I13629. Grave Identifier: 213 / 3259; AF006. Grave type: collective grave. Skeletal information: juvenile (12 years); molar (LLM2). Grave goods: none. Dating: context date 2950-2600 BCE (PSUAMS generated no collagen; skeletal element: left mandibular condyle). Additional information: excavated during the late 19th century.

Genetic Identifier: I13630. Grave Identifier: 213 / 3274; AF007. Grave type: collective grave. Skeletal information: molar. Grave goods: none. Dating: radiocarbon date 2882-2669 calBCE (4174±21 BP, OxA-39062; skeletal element: molar). Additional information: excavated during the late 19th century.

Genetic Identifier: I13631. Grave Identifier: 213 / 15.019; AF008. Grave type: collective grave. Skeletal information: left femur. Grave goods: none. Dating: context date 2950-2600 BCE. Additional information: excavated during the late 19th century.

Genetic Identifier: I13638. Grave Identifier: 213 / 3252; AF014. Grave type: collective grave. Skeletal information: molar. Grave goods: none. Dating: radiocarbon date 2886-2668 calBCE (4180±25 BP, PSUAMS-12142; skeletal element: tooth). Additional information: excavated during the late 19th century.

Genetic Identifier: I13649. Grave Identifier: 212 / 1x.021; AF026. Grave type: collective grave. Skeletal information: femur. Grave goods: none. Dating: context date 3050-2600 BCE. Additional information: excavated during the late 19th century.

Genetic Identifier: I13651. Grave Identifier: 212 / 3245; AF028. Grave type: collective grave. Skeletal information: femur. Grave goods: none. Dating: context date 3050-2600 BCE. Additional information: excavated during the late 19th century.

Genetic Identifier: I13653. Grave Identifier: 213 / 3251; AF030. Grave type: collective grave. Skeletal information: molar. Grave goods: none. Dating: context date 3050-2600 BCE. Additional information: excavated during the late 19th century.

**Dating:** Radiocarbon dates on two right ulnas suggest a Late Neolithic chronology for some of the human remains (Toussaint 2003): 4195±40 (OxA-10687 / 4846–4581 cal. BP) and 4265±40 (OxA-10688 / 4959–4648 cal. BP).

**Source of the samples: University of Huddersfield**

**Authors of the entry:** Maria Pala, Alessandro Fichera, Damien Flas

**References**:

Fraipont, J. (1897). La grotte du mont Falhise [Anthée]. *Bulletins de l'Académie royale des sciences, des lettres et des beaux-arts de Belgique.* Bruxelles: Académie royale des sciences, des lettres et des beaux-arts de Belgique.

Haeck, J. (1964). La grotte du Mont Falise à Antheit, vallée de la Méhaigne, province de Liège. *Bulletin de la Société royale belge d'Anthropologie et de Préhistoire* 74, 39-54.

Toussaint, M. (2003). Wanze/Antheit: apport des datations radiocarbones d'ossements humains de la grotte du Mont Falise à la problématique des sépultures protohistoriques en milieu karstique. *Chronique de l'Archéologie Wallonne* 11, 99-101.

#### 2.17 Abri Sandron (Liège, Belgium)

**Analyzed samples:**

| | | |
|---|---|---|
| I13633 | (97; AF010): | 2916-2781 calBCE (4260±25 BP, PSUAMS-12141) |

| | | |
|---|---|---|
| I13635 | (89 / 6763; AF012): | 2660-2468 calBCE (4035±30 BP PSUAMS-11920) |
| I13654 | (X1; AF031): | 2950-2650 BCE |
| I13655 | (88 / 6165; AF032); | 2950-2650 BCE |
| I13656 | (90; AF033): | 2950-2650 BCE |
| I13657 | (91; AF034): | 2917-2786 calBCE (4265±25 BP, PSUAMS-12143) |
| I13659 | (94; AF036): | 2911-2706 calBCE (4245±25 BP, PSUAMS-12144) |
| I13660 | (95; AF037): | 2865-2502 calBCE (4103±29 BP, UBA-42622) |

**Contact information:** Maria Pala, Damien Flas, Pierre Noiret **Site information and excavation history:**

Abri Sandron is a rock-shelter located in the Mehaigne valley, near Huccorgne (Liege Province). The site has been excavated several times between the 1880s and 1960s. The main excavations were organised by Julien Fraipont and Fernand Tihon in 1887 and 1888 (Fraipoint 1898) and yielded the human remains sampled here. Besides hundreds of bones, corresponding to at least 15 individuals (Toussaint, 2002, Fraipont, 1898), the archaeological material includes Pleistocene (Middle and Upper Palaeolithic; (Otte, 1979) and Holocene (mostly Neolithic) fauna and artefacts.

**Summary of the sampled materials:** The human remains included in this study were part of a collection excavated in 1887 and 1888 by Julien Fraipont and Fernand Tihon (Fraipoint 1898).

Genetic Identifier: I13633. Grave Identifier: 97; AF010. Grave type: collective grave. Skeletal information: molar (LRM3). Grave goods: none. Dating: radiocarbon date 2916-2781 calBCE (4260±25 BP, PSUAMS-12141; skeletal element: mandible). Additional information: excavated during the late 19th century.

Genetic Identifier: I13635. Grave Identifier: 89 / 6763; AF012. Grave type: collective grave. Skeletal information: adult; left petrous. Grave goods: none. Dating: radiocarbon date 2660-2468 calBCE (4035±30 BP, PSUAMS-11920; skeletal element: cranium fragment). Additional information: excavated during the late 19th century.

Genetic Identifier: I13654. Grave Identifier: X1; AF031. Grave type: collective grave. Skeletal information: molar (M1). Grave goods: none. Dating: context date 2950-2650 BCE. Additional information: excavated during the late 19th century.

Genetic Identifier: I13655. Grave Identifier: 88 / 6165; AF032. Grave type: collective grave. Skeletal information: molar (RM3). Grave goods: none. Dating: context date 2950-2650 BCE. Additional information: excavated during the late 19th century.

Genetic Identifier: I13656. Grave Identifier: 90; AF033. Grave type: collective grave. Skeletal information: molar (LLM3). Grave goods: none. Dating: context date 2950-2650 BCE. Additional information: excavated during the late 19th century.

Genetic Identifier: I13657. Grave Identifier: 91; AF034. Grave type: collective grave. Skeletal information: juvenile; molar (ULM3). Grave goods: none. Dating: radiocarbon date 2917-2786 calBCE (4265±25 BP, PSUAMS-12143; skeletal element: maxilla). Additional information excavated during the late 19th century.

Genetic Identifier: I13659. Grave Identifier: 94; AF036. Grave type: Collective grave. Skeletal information: none. Grave goods: molar (LLM). Dating: radiocarbon date 2911-2706 calBCE (4245±25 BP, PSUAMS-12144; skeletal element: mandibular molar). Additional information excavated during the late 19th century.

Genetic Identifier: I13660. Grave Identifier: 95; AF037. Grave type: collective grave. Skeletal information: juvenile; molar (LLM2). Grave goods: none. Dating: radiocarbon date 2865-2502 calBCE (4103±29 BP, UBA-42622; skeletal element: mandible). Additional information: excavated during the late 19th century.

**Dating:** Radiocarbon dating on three human 2nd right metacarpals suggests a Late Neolithic chronology (Toussaint, 2002): 4235±45 (OxA-10555 / 4868–4587 cal. BP), 4183±38 (OxA-10556 / 4839–4580 cal. BP) and 4280±40 (OxA-10557 / 4963–4655 cal. BP).

**Source of the samples:**

**Authors of the entry:** Maria Pala, Alessandro Fichera, Damien Flas

**References**:

Fraipont, J. (1898). Les Néolithiques de la Meuse. Type de Furfooz. *Bulletin de la Société d'Anthropologie de Bruxelles.* Bruxelles.

Otte, M. (1979). *Le paléolithique supérieur ancien en Belgique*, Bruxelles : Musées royaux d'art et d'histoire.

Toussaint, M. (2002). Problématique chronologique des sépultures du Mésolithique mosan en milieu karstique. *Notae Praehistoricae,* 22**,** 141-166.

#### 2.18 Pommerœul (Hainault, Belgium)

**Analyzed samples:**

I18068 (T26-C): 3011-2890 cal BCE (4320 ± 27 BP, RICH-27887)

I21570 (T26-J): 3017-2906 cal BCE (4278+/-27 BP, RICH-27891)

These samples have been reported in detail by Veselka, B. et al. 2024. Assembling Ancestors: the manipulation of Neolithic and Gallo-Roman skeletal remains from Pommeroeul, Belgium. *Antiquity online*, 1-16.

#### 2.19 Grotte de la faille du Burin (Namur, Belgium)

**Analyzed samples:**

I7010 (BELG_265): 8000-5500 BCE

**Contact person:** Michel Toussaint

**Site information and excavation history:** The Faille du Burin is a small cave at the base of the rocks surmounted by the medieval castle of Samson, just above the slopes that descend towards the Samson river, a tributary of the Meuse river. The cavity opens to the northeast and appears as a narrow crack that extends southwest for nearly 8 m before opening into a small room of about 1.50 $m^2$, where the human bones were found by P. Lacroix who excavated dozens of cranial and postcranial bones (Toussaint & Lacroix, 2002). The bones correspond to at least four adults and two children. No archaeological material was collected in direct contact with the bones; a flint blade was discovered in the access corridor. It seems however that, during previous speleologist work, the anterior part of the cavity has also yielded human bones dating back to the Neolithic.

**Summary of the sampled materials:**The human bone analyzed in this article comes from the small room located about ten meters from the entrance of the cave. The sampled individual in this study is I7010.

**Dating:** Four AMS radiocarbon dates were obtained from four left navicular bones from four different subjects found in the small room at the back of the cavity. They provided the following results:

| Date (BP) | Lab code | Sample material | $\delta^{13}C$ | $\delta^{15}N$ | cal. BP (2σ) | cal. BCE (2σ) |
|---|---|---|---|---|---|---|
| 9520 ± 55 BP | OxA-10585 | Left navicular bone (adult human) | - 19.6 | | 11090-10595 | 9140-8645 |
| 9345 ± 75 BP | OxA-8938 | Left navicular bone (adult human) | -19.6 | | 10750-10290 | 8800-8340 |
| 9335 ± 65 BP | OxA-10595 | Left navicular bone (adult human) | -19.6 | | 10715-10295 | 8765-8345 |
| 9315 ± 50 BP | OxA-10564 | Left navicular bone (adult human) | -20.0 | | 10660-10300 | 8710-8350 |

These radiocarbon dates are consistent with the already rich corpus of radiocarbon results obtained from early Mesolithic burials of our regions (Toussaint, 2010 ; Meiklejohn *et al.*, 2014).

**Source of the samples:** The material discovered at the "Faille du Burin" is in the process of being deposited in public collections.

**Authors of the entry:** Michel Toussaint

**References**:

Meiklejohn, C., Miller, B., & Toussaint, M. (2014). Radiocarbon dating of Mesolithic human remains in Belgium and Luxembourg. *Mesolithic Miscellany* 22, 10–39.

Toussaint M. (2010). Les sépultures mésolithiques du bassin mosan wallon : où en est la recherche en 2010 ? *Bulletin des Chercheurs de la Wallonie*, hors-série n° 2, 69-86.

Toussaint M. & Lacroix Ph. (2002). Andenne/Thon : la Faille du Burin à Samson, une nouvelle sépulture collective du Mésolithique ancien. *Chronique de l'Archéologie wallonne*, 10/2002, 228-230.

#### 2.20 Grotte Rousseau (Lustin, Profondeville, Namur, Belgium)

**Analyzed samples:**

I7012 (BELG_267): 2618-2468 calBCE (4020±25 BP, PSUAMS-7872)

I7018 (BELG_7764): 8547-8293 calBCE (9190±45 BP, PSUAMS-7873)

**Contact person:** Michel Toussaint

**Site information and excavation history:** This small cave was discovered during speleological and archaeological surveys by Philippe Lacroix in the 1980s and 1990s, near the Meuse river. It has not been the subject of detailed excavations, but only small surveys. The most spectacular piece collected is a part of a right human foot with bone ankylosis (Masy, 1997).

**Summary of the sampled materials:** The human bones analyzed in this article were found near the entrance of the cave. Two individuals have been sequenced and have been directly radiocarbon dated. Interestingly one dates to the Mesolithic (I7018), the other to the Neolithic (I7012).

**Dating**: Two AMS radiocarbon dates were obtained from fifth metatarsals from two different individuals found near the entrance of the cavity. They provided the following results (Toussaint *et al.*, 2020):

| **Date (BP)** | **Lab code** | **Sample material** | **cal. BP (2σ)** | **cal. BCE (2σ)** |
|---|---|---|---|---|
| 4270 ± 70 BP | OxA-8877 | Adult fifth metatarsal | 5040-4580 | 3090-2630 |
| 4150 ± 50 BP | OxA-8809 | Adult fifth metatarsal | 4830-4530 | 2880-2580 |

Two direct dates were obtained on the sampled individuals (I7012 and I7018).

| **Date (BP)** | **Lab code** | **Sample material** | **cal. BP (2σ)** | **cal. BCE (2σ)** |
|---|---|---|---|---|
| 4020 ± 25 | PSUAMS-7872 | Human bone (I7012) | 4565-4415 | 2620-1470 |
| 9190 ± 45 BP | PSUAMS-7873 | Human bone (I7018) | 10495-10240 | 8545-8295 |

**Source of the samples:** The anthropological material discovered at the Rousseau cave are in the process of being deposited in public collections.

Authors of the entry: Michel Toussaint

**References** :

Masy Ph. (1997). Notice, in descriptive catalogue in *Le secret des dolmen*s. Wéris, Musée des Mégalithes, 145.

Toussaint M., Smolderen A., Bocherens H., Cattelain L., Collin J.-Ph. & Cattelain P. (2020). La Grotte Ambre à Matagne-la-Grande (Doische, Namur, Belgique) : étude anthropologique, biogéochimique et archéologique d'un amas d'ossements humains du Néolithique final du bassin mosan wallon. *Archeo-Situla* 39, 63-100.

#### 2.21 Wéris (Durbuy, Luxembourg, Belgium)

**Analyzed samples:**

I7014 (BELG_273): 3500-2500 BCE

**Contact person:** Michel Toussaint

**Site information and excavation history:** The megalithic complex of Wéris (Durbuy, province of Luxembourg) stretches over some 8 km long and 300 m wide, on two plateaus aligned on either side of the Aisne, an Ardennes tributary on the right bank of the Ourthe river. The sites consists of two passage graves (Wéris I and Wéris II) with associated standing stones, and six sites composed of one or more menhirs. All these monuments are made of puddingstone, a rock that has formed in natural benches on the Ardennes ridge overlooking the Wéris plateau (Toussaint, 2003; Toussaint *et al.*, 2009).

The interest of the Wéris megalithic field has long been recognised. The two passage graves and the site of the menhirs of Oppagne were the subject of initial excavations at the end of the 19th century and the beginning of the 20th century. Recent excavations at the end of the 20th century and the beginning of the 21st century were still carried out in the two passage graves but also in a series of sites with standing stones discovered on this occasion

The two passage graves are built according to the same rectangular plan. They both have an ante-chamber, an elongated burial chamber and a posterior slab lying behind the back stone. It is in these two alleys, clearly formerly disturbed, that very rare archaeological and anthropological material was discovered both during the first excavations and recent excavations.

**Summary of the sampled materials:** The human bone analyzed in this article comes from the Wéris II covered alley (I7014).

**Dating**: Five $^{14}C$ dates were obtained at the Wéris megalithic field, all using human bones:

*Wéris I* :

| Date (BP) | Lab code | Sample material | $\delta^{13}C$ | $\delta^{15}N$ | cal. BP (2σ) | cal. BCE (2σ) |
|---|---|---|---|---|---|---|
| 4240 ± 65 BP | OxA-6457 | Adult phalange | | | 4960-4575 | 3010-2625 |
| 4170 ± 60 BP | OxA-6458 | Fragment of maxilla | | | 4840-4530 | 2895-2580 |

*Wéris II* :

| Date (BP) | Lab code | Sample material | $\delta^{13}C$ | $\delta^{15}N$ | cal. BP (2σ) | cal. BCE (2σ) |
|---|---|---|---|---|---|---|
| 4240 ± 45 BP | OxA-8956 | Adult second left metacarpal | - 20.4 | | 4875-4585 | 2925-2640 |
| 4180 ± 40 BP | OxA-8939 | Adult five left metatarsal | - 20.5 | | 4840-4580 | 2890-2630 |

*Heyd standing stone* :

| Date (BP) | Lab code | Sample material | $\delta^{13}C$ | $\delta^{15}N$ | cal. BP (2σ) | cal. BCE (2σ) |
|---|---|---|---|---|---|---|
| 4425 ± 45 BP | OxA-8828 | Human clavicle | - 20.9 | | 5280-4865 | 3330-2920 |

**Source of the samples:** The archaeological and anthropological material discovered in Wéris is very poor due to old alterations of the two passage graves. The few discoveries from the first excavations, at the end of the 19th century and the beginning of the 20th century, are kept at the Archaeological Museum in Arlon. The archives and the few objects from the recent excavations are kept in Namur at the "Agence wallonne du Patrimoine, Service public de Wallonie.

Authors of the entry: Michel Toussaint

References :

Toussaint M. (ed.), (2003). Le « champ mégalithique de Wéris ». Fouilles de 1979 à 2001. Volume 1. Contexte archéologique et géologique, Namur, Études et Documents, Archéologie, 9, 448 p..

Toussaint M. Frébutte C. & hubert F. (eds.), (2009). *Le « champ mégalithique de Wéris ». Fouilles de 1979 à 2001, volume 2. Rapports de fouilles*. Namur, Études et Documents, Archéologie 15, 320 p.

#### 2.22 Claminforge (Sambreville, Namur, Belgium)

**Sample analyzed**:

I7015 (BELG_6598): 9000-8500 BCE

**Contact person:** Michel Toussaint

**Site information and excavation history:** Claminforge is a small cave site on the left bank of the Bième, a tributary of the Sambre. The site was first discovered in 1988 by a group of speleologists of the Centre Spéléologique de la Basse Sambre(Meiklejohn et al. 2014). In 1995 a small rescue excavation was carried out by M. Toussaint (Toussaint 2019). The front part of the site was formerly destroyed by the activity of a limestone quarry. Therefore the human remains discovered represent only part of the original bone deposit. All of them, often fragmentary, have been discovered at the end narrow gallery of barely two square meters, most of them by speleologists in 1988. Seven individuals have been recovered from the cave. Direct dating of two bones, carried out just after the 1995 excavation, corresponds to the early Mesolithic. However, new, as yet unpublished datings carried out recently at Ghent University show that some of the other individuals correspond to the Neolithic. This discovery could possibly be interpreted as two successive secondary burials, one Mesolithic and one Neolithic, but more probably as rejections of bones intended to make room for new corpses.

**Summary of the sampled materials:** The present assemblage gives evidence for the presence of at least seven individuals: four adults and three children aged between 8 and 11 years (Toussaint 2019). One of these individuals yielded sample I7015.

**Dating:** Two of the individuals have been dated: after the rescue excavation in 1995 Toussaint discovered, a human third metacarpal, which yielded a first $^{14}$C date; a cervical vertebra found a year later, yielded a second date (Toussaint 2019, 255).

| Date (BP) | Lab code | Sample material | $\delta^{13}C$ | $\delta^{15}N$ | cal. BP (2σ) | cal. BCE (2σ) |
|---|---|---|---|---|---|---|
| 9320 ± 75 | OxA-5451 | Cervical vertebra | -19.4 | 10.5 | 10705-10264 | 8755-8315 |
| 9525 ± 60 | OxA-10552 | 3rd metacarpal | -19.4 | --- | 11105-10590 | 9156-8642 |

**Source of the samples:** The bones of Claminforge are preserved at the Prehistomuseum of Ramioul, in Flémalle, in the province of Liège, Belgium.

**Authors of the entry:** Michel Toussaint

**References :**

Meiklejohn, C., Miller, B., & Toussaint, M. (2014). Radiocarbon dating of Mesolithic human remains in Belgium and Luxembourg. *Mesolithic Miscellany* 22, p.10–39.

Toussaint, M. (2019). Les ossements humains du Mésolithique ancien de la grotte de Claminforge (Sambreville, province de Namur, Belgique). *Bulletin des Chercheurs de la Wallonie, Tome LIV*, 251–281.

### Germany

#### 2.23 Blätterhöhle cave (Westphalia, Germany)

**Samples used:**

| | | |
|---|---|---|
| I1563 | Bla5+Bla7+Bal13+Bla26(o)+Bla30+Bla54 (Excavation 2004) \| Bla5+Bla7+Bal13+Bla26(o)+Bla30+Bla54 (Excavation 2014) | 3626-3378 calBCE (4726±17 BP) [R_combine: (4580±30, KIA-28844, Bla5); (4860±30, KIA-45011, Bla7); (4730±25, KIA-45010, Bla13)] |
| I1565 | Bla8+Bla9+Bla11+Bla24+Bla26(x)+Bla45 (Excavation 2004) \| Bla8+Bla9+Bla11+Bla24+Bla26(x)+Bla45 (Excavation 2014) | 3725-3655 calBCE (4965±15 BP) [R_combine: (4950±30, KIA-45006, Bla8); (4905±25, KIA-45008, Bla9); (5145±30, KIA-45007, Bla11); (4845±35, KIA-37507, Bla24)] |
| I1593 | Bla16+Bla27+Bla59 (Excavation 2004) \| Bla16+Bla27+Bla59 (Excavation 2014) | 3644-3528 calBCE (4810±23 BP) [R_combine: (4615±30, KIA-28845, Bla16); (5055±35, KIA-37508, Bla27)] |
| I1594 | Bla28 (Excavation 2004) | 3338-3024 calBCE (4465±30 BP, KIA-28846) |

A detailed description and analysis is published by Lipson, M., Szécsényi-Nagy, A., Mallick, S., Pósa, A., Stégmár, B., Keerl, V., Rohland, N., Stewardson, K., Ferry, M., Michel, M., Oppenheimer, J., Broomandkhoshbacht, N., Harney, E., Nordenfelt, S., Llamas, B., Gusztáv Mende, B., Köhler, K., Oross,

K., Bondár, M., … Reich, D. (2017). Parallel palaeogenomic transects reveal complex genetic history of early European farmers. *Nature*, *551*(7680), 368–372. https://doi.org/10.1038/nature24476

#### 2.24 Niedertiefenbach (Germany)

**Samples used:**

| | | |
|---|---|---|
| KH150189_KH150632_KH150636 | NT1 | 3346-3095 calBCE (4493±26 BP, KIA-53052) |
| KH150190 | NT21 | 3500-2800 BCE |
| KH150191 | NT39 | 3500-2800 BCE |
| KH150193_KH150286 | NT9A+ 9 | 3500-2800 BCE |
| KH150195_KH150196_KH150616 | NT23/123/128b | 3342-3096 calBCE (4492±24 BP, KIA-52274) |
| KH150197 | NT122.1 | 3500-2800 BCE |
| KH150198 | NT186 | 3500-2800 BCE |
| KH150200 | NT136.2 | 3500-2800 BCE |
| KH150203 | NT121 | 3339-3036 calBCE (4481±24 BP, KIA-52267) |
| KH150204_KH150634 | KH150204_KH150634 | 3352-3099 calBCE (4507±25 BP, KIA-53051) |
| KH150208_KH150210 | NT26 /111 | 3500-2800 BCE |
| KH150287 | NT54 | 3346-3098 calBCE (4499±25 BP, KIA-52268) |
| KH150289 | NT98 | 3346-3098 calBCE (4499±24 BP, KIA-52270) |
| KH150418 | NT41+45+43 | 3500-2800 BCE |
| KH150419 | NT27 | 3500-2800 BCE |
| KH150422 | NT46 | 3366-3103 calBCE (4538±24 BP, KIA-52272) |
| KH150610 | NT17 | 3365-3102 calBCE (4532±25 BP, KIA-52273) |
| KH150612 | NT5+6.2 | 3500-2800 BCE |
| KH150613_KH180043 | NT107 | 3346-3098 calBCE (4499±26 BP, KIA-53045) |
| KH150614_KH150615 | NT142.1 | 3334-3025 calBCE (4462±24 BP, KIA-53046) |
| KH150618 | NT48 | 3336-3028 calBCE (4468±25 BP, KIA-52275) |
| KH150619 | NT42 | 3333-3021 calBCE (4455±24 BP, KIA-52276) |
| KH150620 | NT148 | 3344-3094 calBCE (4491±25 BP, KIA-53047) |
| KH150621 | NT50 | 3500-2800 BCE |
| KH150622 | NT130 | 3264-2926 calBCE (4417±19 BP, KIA-53048) |
| KH150623 | NT135 | 3500-2800 BCE |
| KH150625 | NT83 | 3348-3096 calBCE (4497±27 BP, KIA-52277) |
| KH150626 | NT58 | 3500-2800 BCE |

| | | |
|---|---|---|
| KH150627 | KI11 | 3335-3024 calBCE (4462±27 BP, KIA-52278) |
| KH150628 | NT150.1 | 3334-2938 calBCE (4448±26 BP, KIA-53049) |
| KH150629 | NT30 | 3500-2800 BCE |
| KH150630 | KI12 | 3487-3106 calBCE (4564±25 BP, KIA-53050) |
| KH150633 | KI13 | 3344-3036 calBCE (4486±29 BP, KIA-52279) |
| KH150635 | KI14 | 3322-2924 calBCE (4425±28 BP, KIA-52280) |
| KH150637 | NT110 | 3328-2926 calBCE (4432±28 BP, KIA-52281) |
| KH150639 | NT49 | 3500-2800 BCE |
| KH150640 | NT98 | 3334-3024 calBCE (4461±25 BP, KIA-53053) |
| KH150641 | KI15 | 3339-3028 calBCE (4473±28 BP, KIA-52282) |
| KH180044 | NT136.1 | 3500-2800 BCE |
| KH180045 | NT146 | 3500-2800 BCE |

A detailed analysis was published in: Immel, A., Pierini, F., Rinne, C., Meadows, J., Barquera, R., Szolek, A., Susat, J., Böhme, L., Dose, J., Bonczarowska, J., Drummer, C., Fuchs, K., Ellinghaus, D., Kässens, J. C., Furholt, M., Kohlbacher, O., Schade-Lindig, S., Franke, A., Schreiber, S., … Krause-Kyora, B. (2021). Genome-wide study of a Neolithic Wartberg grave community reveals distinct HLA variation and hunter-gatherer ancestry. *Communications Biology*, *4*(1), 113. https://doi.org/10.1038/s42003-020-01627-4

#### 2.25 Spiekeroog, Wittmund, Lower Saxony (Germany)

**Sample Used:**

SPI001 Site No.: 2212/2:1 5558-5373 calBCE (6510±40 BP; Poz-103001)

**Contact information:** Jan F. Kegler

**Site information and excavation history:**

The site is located on the northern beach of the North Sea island of Spiekeroog about 3 km northeast of the town center. The find was discovered by volunteers (M. Huus) on 04.07.2016 and handed over to the Archaeological Service of the Ostfriesische Landschaft. No further archaeological finds could be made. The object comes from an undefined archaeological context. No follow-up investigation has been carried out on site.

**Summary of the sampled materials:**

Skeletal information: mandible, male, adolescent, min. 40 years. Grave goods: none. Dating: AMS-radiocarbon date 5566 - 5355 calBCE (6510 ± 40 BP; Poz-103001). Additional information:

Wetland find of a human mandible, found during survey in 2016. The bone shows all the characteristics of wet soil preservation. Very robust mandible without knots, dentate with heavily abraded molars. Toothed with right P2, M1 and M2, left M1, M2 and M3. Carious dentition (interdental caries right P2).

An anthropological examination by Dr. S. Grefen-Peters, Braunschweig, confirmed the archaic character of the lower jaw, which comes from a man who probably died at the age of at least 40 years.

Isotope analyses by the University of Warsaw and the Curt Engelhorn Center for Archaeometry confirm a diet with a high content of marine food (probably fish, waterfowl and seal). The 87Sr/86Sr analyses from CEZA underline a local resident population. Genetic material has been extracted from the M2 Molar.

**Dating:**

The molar was dated by the $^{14}$C laboratory in Poznan to 5558-5373 cal BCE (6510±40 BP; Poz-103001)
**Source of the samples:** Archäologischer Dienst & Forschungsinstitut Ostfriesische Landschaft, Aurich

**Authors of the entry:** Jan F. Kegler

**References**:

Kegler, J.F., Grefen-Peters, S. (2019a): Männer aus dem Meer. Archäologie in Deutschland 02-2019, 59.

Kegler, J.F. u. Grefen-Peters, S. (2019b): Meermänner - Anthropologische Spülsaumfunde von Spiekeroog und Baltrum. Archäologie in Niedersachsen 22, 110-115.

#### 2.26 Baltrum, Wittmund, Lower Saxony (Germany)

**Sample Used:**

BLR001 Site No.: 2210/5:2-1 3795-3655 calBCE (4905 ± 30 BP; Poz-103000)

**Contact information:** Jan F. Kegler

**Site information and excavation history:**

The site is located on the northern beach of the North Sea island of Baltrum about 4,5 km northeast of the town center. The find was discovered by volunteers (Chr. Groger) on 21.03 2018 and handed over to the Archaeological Service of the Ostfriesische Landschaft. No further archaeological finds could be made. The object comes from an undefined archaeological context. No follow-up investigation has been carried out on site

**Summary of the sampled materials:**

An anthropological examination by Dr. S. Grefen-Peters, Braunschweig, confirmed the archaic character of the lower jaw, which comes from an adult man who probably died at the age in between 20 to 50 years. Isotope analyses by the University of Warsaw and the Curt Engelhorn Center for Archaeometry (CEZA) confirm a diet with a high content of marine food (probably fish, waterfowl and seal). 87Sr/86Sr analyses from CEZA underline a local resident population.
Genetic material has been extracted from the M2 Molar.

The fragment of site 2210/5:2-1 (sample BLR001) is of a mandible that belonged to an adolescent male between 20-50 years old. There were no grave goods. The bone shows all the characteristics of wet soil preservation. The mandible fragment is very robust. The right side of the jaw is incomplete, the right branch and the corpus with the premolar and molar tooth fan are missing. Toothed with left M1, M2 and M3.

**Dating:**

$^{14}C$ dating (AMS) by the 14C laboratory in Poznan confirms a Neolithic age: 3795-3655 calBCE (4905 ± 30 BP; Poz-103000).

**Source of the samples:** Archäologischer Dienst & Forschungsinstitut Ostfriesische Landschaft, Aurich

**Authors of the entry:** Jan F. Kegler

**References:**

Kegler, J.F., Grefen-Peters, S. (2019a): Männer aus dem Meer. Archäologie in Deutschland 02-2019, 59.

Kegler, J.F. u. Grefen-Peters, S. (2019b): Meermänner - Anthropologische Spülsaumfunde von Spiekeroog und Baltrum. Archäologie in Niedersachsen 22, 110-115.

### SI 3 Analytical details Sr-O-C isotope analysis

The dental elements selected for combined Sr-O-C isotope analysis (N=23) were mechanically cleaned at the Archaeological and Forensic Sample Preparation Laboratory of the Vrije Universiteit Amsterdam or, in case of 'Cees' from Mienakker (supplementary data 2.7), at the depot where the skeleton was housed, using a Proxxon drill equipped with a ball-shaped, acid-cleaned (10% HCl), diamond-coated grinding bit. Approximately 10 mg of white enamel powder was collected in clean glass vials.

For Sr isotope analysis, around 2 ± 1 mg of enamel powder was subsampled into acid-cleaned (6–7M HCl) Eppendorf® centrifuge tubes and transferred to the USA class 100 (ISO 5) clean laboratory, equipped with USA class 10 (ISO 4) laminar flow hoods at the same university. Additionally, 0.3 mg ± 10% was subsampled into clean, screw-capped Exetainer® vials and sent to the Stable Isotope Laboratory at the Vrije Universiteit Amsterdam for O-C isotope analysis.

Depending on the physical quality of the sample, the enamel powder underwent leaching with 0.1M acetic acid, followed by Milli-Q water rinsing and dissolution in 500 µl 3M $HNO_3$. Strontium extraction and sample loading were conducted following previously established protocols[110]. Isotope compositions were measured using a Thermo Scientific™ Triton Plus™ instrument housed at the Vrije Universiteit Amsterdam. Strontium ratios were determined via a static routine and corrected for mass fractionation to an $^{86}Sr/^{88}Sr$ of 0.1194. The long-term reproducibility was ± 0.000008 based on repeated analysis of the NIST® SRM® 987 standard during the course of the study (2019-2022: n = 393, 1σ, loading size: 200 ng). Procedural blanks (n = 7) contained between a negligible amount strontium. The $^{87}Sr/^{86}Sr$ are reported with a ±2 standard error (2SE), representing the analytical uncertainty derived from 240 measurements (12 blocks of 20 cycles) per run.

The $\delta^{18}O$ and $\delta^{13}C$ values were analyzed using a Thermo Finnigan GasBench II preparation device connected to a Thermo Finnigan Delta+ mass spectrometer. The data were normalized to the Vienna Peedee Belemnite (VPDB) scale using an in-house carbonate reference material (VICS), calibrated against NBS19 and LSVEC certified reference materials. Instrument performance was verified using the international control standard IAEA-603, which yielded average values of 2.40‰ for $\delta^{13}C$ and −2.55 ‰ for $\delta^{18}O$ (n = 31). The reproducibility of IAEA-603 during the analytical session was 0.12‰ (1σ).

### SI 4. *qpAdm* modeling of ancestry proportions

We used *qpAdm*[111] to estimate ancestry proportions. We set the parameters allsnps: YES and inbreed: YES to account for the use of pseudo-haploid data.

As outgroups for our models, we used the same population set as in Patterson *et al.*[112], but adding West Siberian hunter-gatherers (WSHG) and a group composed by Caucasus hunter-gatherers (CHG) and Iran Neolithic individuals. The outgroup list is as follows:

- OldAfrica: a pool of diverse ancient African individuals with no evidence of recent West Eurasian-related admixture that we use as a deeply divergent outgroup.
- Afanasievo: a group of individuals from the Altai region of Russia and Mongolia with very similar ancestry to Steppe_EBA.
- WHGB: European hunter-gatherers that were genetically very similar to WHG but with slightly more Eastern European Hunter-Gatherer relatedness and mostly from the Iron Gates region of the Danube river in southeastern Europe
- Anatolia_N: Anatolian Neolithic farmers very similar genetically to Early European farmers (EEF).
- WSHG
- CHG_IranNeolithic

We found that the addition of these two populations to the outgroup set adds leverage to tease apart WHG-related ancestry, Eastern hunter-gatherer-related ancestry (EHG) and Steppe Early Bronze Age-related ancestry.

We began by computing by-individual ancestry proportions using a one-way model with 12 Late Paleolithic and Mesolithic individuals from Western Europe (WHG) as a proxy for the ancestry present in Western Europe before the arrival of Neolithic farmers (Supplementary Table 3). Of the 109 individuals from the Rhine-Meuse delta region in our dataset, only the 10 Mesolithic individuals and two Swifterbant individuals can be modelled successfully (Supplementary Table 3). The remaining individuals fail with extremely low p-values, which indicates that they harbored additional ancestry layers. By adding a second source of ancestry, the Balkan Neolithic individuals (Balkan_N) as a proxy for European Early Farmers (EEF) with western Anatolian farmer-derived ancestry, a large number of individuals can be successfully modelled. This includes the vast majority of the Neolithic individuals, one Vlaardingen/Corded Ware individual, and the Tiel Medel individuals with uncertain chronology (most likely Middle Neolithic according to IBD data). The remaining Neolithic individuals fail with p-values 0.00015 and 0.049, but for all except two we obtained a good fit by adding Eastern European hunter gatherers (EHG) as a third source, with proportions 4-10%.

The remaining 20 individuals without a good p-value in all the previous models all come from Vlaardingen/Corded Ware, Bell Beaker and Early Bronze Age contexts. By adding a steppe related source, either a Steppe Early Bronze Age group (Steppe_EBA) or a Corded Ware group from Germany (Germany_CordedWare), we were able to successfully model most of these individuals with very high proportions of steppe-associated ancestry, with the exception of the two Vlaardingen/Corded Ware individuals who required lower proportions than the rest.

Next, we wished to test for sex bias in the admixture between EEF and Mesolithic hunter-gatherers. To increase resolution, we merged all the Neolithic individuals from the Rhine-Meuse delta (excluding the Late Neolithic ones with steppe ancestry) with capture data under one group (Lower_Rhine_Neolithic) and tested the following two-way models:

Germany_EN_LBK+ WHG

Czechia_N_TRB+ WHG

We repeated this analysis for autosomal data only, and separately for the X-chromosome only (Supplementary Table 4).

We used Germany Early Neolithic LBK farmers and Czechia Neolithic TRB farmers because they represent more proximal sources for the EEF-related ancestry (as opposed to Balkan_N), and our goal is to study evidence for sex-bias in the admixture that occurred in the Rhine-Meuse delta region, rather than studying an integrated signal that also might reflect admixture between people of Anatolian farmers and people of Mesolithic European hunter-gatherer ancestry as they interacted across the European continent prior to arriving in our study region.

If one of the two populations participating in the admixture event is predominantly composed by women, and the other population predominantly by men, we expect the ancestry proportions of the first population to be higher in the X-chromosome than those in the autosomes, as women carry two X-chromosomes for every one carried by men, while the autosomes are equally carried by both biological sexes. In both models we find EEF-related proportions to be ~10% higher than in the autosomes, with Z-scores ranging between 2-3 standard deviations above zero for the EEF ancestry being higher on the X-chromosome. This provides moderately (but not highly) significant evidence of sex bias in the admixture event between Neolithic groups with EEF-related ancestry and hunter-gatherer groups, with EEF-related ancestry being predominantly contributed by females. Together with the independent evidence from mtDNA and Y-chromosome lineages, the data provide a compelling case for sex bias.

Next, we attempted to model the Rhine-Meuse delta individuals with steppe-associated ancestry using proximal models. We grouped them into three clusters of individuals: the two Corded Ware/Vlaardingen individuals with steppe ancestry, the Bell Beaker individuals, and the Early Bronze Age individuals. As a proximal steppe-related source we used Germany_CordedWare, and as a Neolithic-related source we used one of the following populations or a combination of two (Supplementary Table 5):

-MLN_Belgium: Late Neolithic individuals from Belgium with high levels of hunter-gatherer ancestry.

-MN_Wartberg: Late Neolithic Wartberg individuals from Niedertiefenbach (Northwestern Germany) with high levels of hunter-gatherer ancestry, but lower than the Late Neolithic individuals from Belgium.

-Germany_Baalberge_MN: Middle Neolithic Baalberge individuals from Germany.

-Czechia_N_TRB: Neolithic TRB individuals from the Czech Republic.

-Poland_GlobularAmphora: Late Neolithic Globular Amphora individuals from Poland.

-England_Neolithic: Neolithic individuals from England.

-France_Neolithic: Neolithic individuals from France.

-Iberia_Neolithic_Chalcolithic: Neolithic-Chalcolithic individuals from the Iberian Peninsula.

These Neolithic populations represent both local groups from the Rhine-Meuse delta area, as well as Neolithic groups from other areas. Thus, this analysis allowed us to test whether there was evidence for a local component of Neolithic ancestry.

The group formed by the two CordedWare/Vlaardingen individuals (I12902 and I33741) can only be modelled using MLN_Belgium (88.8% ± 2%) as a source for the Neolithic component, which is consistent with the PCA results showing a close affinity between these two groups of individuals. Models combining MLN_Belgium with one of the other Neolithic sources give good-fitting models but with close to zero ancestry proportions for the non-MLN_Belgium source. In contrast, the Bell Beaker and Early Bronze Age groups can be modelled either using MN_Wartberg (17.4% ± 1.6% and 23.6% ± 2%, respectively) alone as proxy for the Neolithic component, or using MLN_Belgium (9.4-11.6% and 7.4-11.9%, respectively) together with one of the sources from non-Rhine-Meuse delta regions (6.0-8.8% and 12.3-17.4%, respectively).

When using MLN_Belgium alone as a source for the Neolithic component, *qpAdm* D-scores clearly show that hunter-gatherer-related outgroups shared significantly more alleles with the fitted model than with the data, indicating that the model includes too much hunter-gatherer ancestry, and thus local MLN_Belgium-like groups cannot be the sole source of the non-Steppe ancestry. Conversely, when using one of the sources from nearby regions alone as a source for the Neolithic component, *qpAdm* D-scores clearly show that hunter-gatherer-related outgroups shared significantly more alleles with the data than with the fitted model, indicating that the model includes too low hunter-gatherer ancestry. In summary, the Neolithic ancestry in Bell Beaker and Early Bronze Age groups derives, at least in part, from the local Neolithic populations from the Rhine-Meuse area with high hunter-gatherer ancestry.

Given the previously observed strong genetic similarity between Bell Beakers-associated individuals from the Rhine-Meuse area and Bell Beaker-associated individuals from England[106], we ran the same models for a group including Bell Beakers from England, but excluding four outlier individuals with higher levels of European Early Farmer ancestry (I14200, I1767, I2416, I5379). These four individuals very likely harbour ancestry from local Neolithic from England and thus are not good representatives of the groups that moved to Britain at the beginning of the Chalcolithic period, and so we tested them as a separate group.

The main group of Bell Beakers from England behave very similarly to the Rhine-Meuse delta Bell Beakers and with matching ancestry proportions (Supplementary Table 5); i.e. non-fitting models when featuring MLN_Belgium or one of the sources from nearby regions alone as a source for the Neolithic component; and a good fit either using MN_Wartberg alone (19.6% ± 1.6%) or a mixture between MLN_Belgium (8.8-11.9%) and one of the sources from nearby regions (8.1-11.5%) as a source for the Neolithic component.

These results again highlight the common origin between the main group of Bell Beakers from England and the Rhine-Meuse delta area. In contrast, the Bell Beaker outliers from England show a very poor fit ($P=1.29x10^{-09}$) when using MN_Wartberg as a source for the Neolithic component and can be modelled (P-value between 0.027-0.105) as a mixture of Germany_CordedWare (61.5-65.0%) and Neolithic populations from outside the Rhine-Meuse area such as Poland_GlobularAmphora, Iberia_Neolithic_Chalcolithic and England_Neolithic (35.0-38.5%).

Our interpretation is that that these Bell Beaker outliers, unlike the main Bell Beaker group, likely represent recent mixtures with local Neolithic populations from Britain, and consequently most their European Neolithic ancestry component is best modeled by British Neolithic populations or other Neolithic populations with similar levels of hunter-gathered ancestry, such as Poland_GlobularAmphora and Iberia_Neolithic_Chalcolithic. As a final check, we tested a Bell Beaker group outside Britain and the Rhine-Meuse area (South-East Germany_BellBeaker) and found very poor fit ($P<1.50x10^{-26}$) when using Neolithic populations from the Rhine-Meuse area with high hunter-gatherer ancestry, corroborating that only Bell Beakers from the Rhine-Meuse area and the main Bell Beaker group from England have evidence of deriving part of their ancestry from Neolithic populations from the Rhine-Meuse area.

Finally, we tested more proximal 2-way and 3-way models for the group of four Bell Beaker outliers from England and for the England Early Bronze Age populations, using either Rhine-Meuse area Bell Beaker-associated individuals or the main England Bell Beaker cluster as sources of steppe-related ancestry.

### SI 5. IBD sharing analysis

We called IBD segments between the Rhine-Meuse delta individuals with high quality data (n=55) and all the previously published ancient individuals from Eurasia with high quality data (n=5169). We followed the same procedure described in Ringbauer *et al.*[113], which involves imputing and phasing the aligned sequenced data with GLIMPSE[114] using haplotypes in the 1000 Genome Project as the reference panel[115], and detecting IBD segments with ancIBD (https://github.com/hringbauer/ancIBD). Pairs of individuals connected by IBD are displayed in Supplementary Table 7.

Three individuals from the site of Tiel Medel have uncertain chronology. The site has a Middle Neolithic Swifterbant occupation but also a Bronze Age occupation phase, and the sampled individuals were not amenable to radiocarbon dating We therefore used their IBD connections to estimate an approximate chronology. Their largest IBD sharing is with I33738, a Middle Neolithic Swifterbant individual from Zoelen de Beldert (Netherlands) dated to 4200-3800 BCE, with whom one of Tiel Medel individuals shares four IBD segments longer than 8 cM (the longest being 20.5 cM), for a total share of 53 cM (Supplementary Table 7). The second and third largest IBD sharing are with a Neolithic individual from Hazleton North (England) who lived 3750-3500 BCE (3 IBD segments for a total of 39 cM) and with a Neolithic individual from Gurgy les Noisats dated to 4836-4606 calBCE (5855±40 BP, Lyon-4446, SacA-8629) (2 IBD segments for a total of 33 cM). Based on these IBD results, a Bronze Age chronological attribution is extremely implausible for these individuals, and we thus approximate their date to the range 3800-3600 BCE, hence within the Middle Neolithic. This chronology fits well with their lack of steppe-associated ancestry in the autosomal genome, which already suggested a pre-2500 BCE date.

**References**

1. Louwe Kooijmans, L. P. Schipluiden: a synthetic view. in *Schipluiden: A Neolithic Settlement on the Dutch North Sea Coast c. 3500 cal BC* (eds. Louwe Kooijmans, L. P. & Jongste, P. F. B.) 485–516 (Leiden University Press, Leiden, 2006).

2. Toussaint, M. Les ossements humains du Mésolithique ancien de la grotte de Claminforge (Sambreville, province de Namur, Belgique). *Bulletin des Chercheurs de la Wallonie, Tome LIV* 251–281 (2019).

3. Toussaint, M. & Lacroix, P. Andenne/Thon: la Faille du Burin à Samson, une nouvelle sépulture collective du Mésolithique ancien. *Chronique de l'Archéologie Wallonne 10* 228–230 (2002).

4. Vos, P. C., Bazelmans, J., van der Meulen, M. & Weerts, H. J. T. *Atlas van Nederland in Het Holoceen*. (Prometheus, Amsterdam, 2024). https://data.overheid.nl/dataset/069b3d67-fa6e-423c-8f80-2bf62c66be94

5. Coles, B. J. Doggerland: a speculative survey. *Proceedings of the Prehistoric Society* **64**, 45–81 (1998).

6. Polet, C. & Cauwe, N. Les squelettes mésolithiques et néolithiques de l'abri des Autours (province de Namur, Belgique). *Comptes Rendus Palevol* **1**, 43–50 (2002).

7. Posth, C. *et al.* Palaeogenomics of Upper Palaeolithic to Neolithic European hunter-gatherers. *Nature* **615**, 117–126 (2023).

8. *Doggerland. Lost World under the North Sea*. (Sidestone Press, Leiden, 2022).

9. *Archeologie in de Betuweroute. Hardinxveld-Giessendam Polderweg: Een Mesolithisch Jachtkamp in Het Rivierengebied (5500-5000 v. Chr.)*. vol. 83 (NS Railinfrabeheer, Utrecht, 2001).

10. Smits, E. & Louwe Kooijmans, L. P. 13 Menselijke skeletresten. in *Archeologie in de Betuweroute. Hardinxveld-Giessendam De Bruin: Een kampplaats uit het Laat-Mesolithicum en het begin van de Swifterbant-cultuur (5500-4450 v.Chr.)* (eds. Louwe Kooijmans, L. P., Koot, C. W., ten Anscher, T. J., Van Wijngaarden, G. J. & Goudswaard, B.) vol. 83 487–498 (Rijksdienst voor het Oudheidkundig Bodemonderzoek, Amersfoort, 2001).

11. Gaffney, V. L., Fitch, S. & Smith, D. N. *Europe's Lost World: The Rediscovery of Doggerland*. (Council for British Archaeology, York, England, 2009).

12. Amkreutz, L. W. S. W. A view from Doggerland – interpreting the Mesolithic-Neolithic transition in the wetlands of the Rhine-Meuse delta (5,500 – 2,500 calBC). in *Stone Age borderland experience: Neolithic and Late Mesolithic parallel societies in the north European plain* (eds. Klimscha, F., Heumueller, M., Raemaekers, D. C. M., Peeters, H. & Terberger, T.) 311–326 (Marie Leidorf GmbH, Rahden, 2022).

13. Walker, J. *et al.* A great wave: the Storegga tsunami and the end of Doggerland? *Antiquity* **94**, 1409–1425 (2020).

14. Ritchie, K. The Ertebølle Fisheries of Denmark, 5400-4000 B. (University of Wisconsin, Madison, 2010).

15. Andersen, S. H. 'Køkkenmøddinger' (Shell Middens) in Denmark: a Survey. *Proceedings of the Prehistoric Society* **66**, 361–384 (2000).

16. Enghoff, I. B. Freshwater fishing at Ringkloster, with a supplement of marine fishes. *Journal of Danish Archaeology* **12**, 99–106 (1995).

17. Raemaekers, D. *et al.* Timing and Pace of Neolithisation in the Dutch Wetlands (c. 5000–3500 cal. BC). *Open Archaeology* **7**, 658–670 (2021).

18. Brusgaard, N. Ø. *et al.* Early animal management in northern Europe: multi-proxy evidence from Swifterbant, the Netherlands. *Antiquity* **98**, 654–671 (2024).

19. Teetaert, D. & Crombé, P. The start of pottery production by hunter-gatherers in the Low Countries (Swifterbant Culture, 5th millennium BC) : a critical assessment of the available radiocarbon dates. *Notae Praehistoricae* **41**, 173–186 (2021).

20. Andersen, S. H. The first pottery in South Scandinavia. in *Pots, Farmers and Foragers. Pottery traditions and social interaction in the earliest Neolithic of the Lower Rhine Area* (eds. Vanmontfort, B., Louwe Kooijmans, L. P., Amkreutz, L. W. S. W. & Verhart, L. B. M.) 167–213 (Leiden University Press, Leiden, 2010).

21. Dreshaj, M., Raemaekers, D. & Dee, M. Chronological modeling on a calibration plateau: implications for the emergence of agriculture in the Dutch wetlands. *Radiocarbon* **65**, 1280–1298 (2023).

22. Crombé, P. *et al.* New evidence on the earliest domesticated animals and possible small-scale husbandry in Atlantic NW Europe. *Sci Rep* **10**, 20083 (2020).

23. Hulst, R. S., Hogestijn, J. W. H., de Haan, M. J. A., Lauwerier, R. C. G. M. & Marswijk, R. W. Buren Zoelen. *Jaarverslag Rijksdienst voor het Oudheidkundig Bodemonderzoek 1992* 69 (1993).

24. Bakels, C. C. *The Western European Loess Belt : Agrarian History, 5300 BC - AD 1000*. (Springer Netherlands, Dordrecht, 2009).

25. Bakels, C. C. *Four Linearbandkeramik Settlements and Their Environment: A Paleoecological Study of Sittard, Stein, Elsloo and Hienheim*. vol. 11 (Leiden (proefschrift), 1978).

26. Kreuz, A. M. *Die Ersten Bauern Mitteleuropas - Eine Archäobotanische Untersuchung Zu Umwelt Und Landwirtschaft Der Ältesten Bandkeramik*. vol. 23 (Leiden (proefschrift), 1991).

27. Čerevková, A. The Subsistence Strategy of Linear Pottery Culture in Moravia (Czech Republic): Current State of Knowledge. **7**, 1473–1491 (2021).

28. Bakels, C. Archaeobotanical investigations in the Aisne valley, northern France, from the neolithic up to the early Middle Ages. *Veget Hist Archaebot* **8**, 71–77 (1999).

29. Denaire, A. *et al.* The Cultural Project: Formal Chronological Modelling of the Early and Middle Neolithic Sequence in Lower Alsace. *Journal of Archaeological Method and Theory* **24**, (2017).

30. Kirschneck, E. The Phenomena La Hoguette and Limburg – Technological Aspects. *Open Archaeology* **7**, 1295–1344 (2021).

31. Constantin, C., Illett, M. & Burnez-Lanotte, L. La Hoguette, Limburg and the Mesolithic: some questions. in *Pots, Farmers and Foragers. Pottery traditions and social interaction in the earliest Neolithic of the Lower Rhine Area* (eds. Vanmontfort, B., Louwe Kooijmans, L. P., Amkreutz, L. W. S. W. & Verhart, L. B. M.) 41–49 (Leiden University Press, Leiden, 2010).

32. Hofmann, D. Keep on walking> the role of migration in Linearbandkeramik life. *Documenta Praehistorica 43* 235–251 (2016).

33. Crombé, P. Mesolithic projectile variability along the southern North Sea basin (NW Europe): Hunter-gatherer responses to repeated climate change at the beginning of the Holocene. *PLoS ONE* **14**, e0219094 (2019).

34. Crombé, P. & Cauwe, N. The Mesolithic. *Anthropologica et praehistorica* **112**, (2001).

35. Cauwe, N., Vander Linden, M. & Vanmontfort, B. The Middle and Late Neolithic. in *Prehistory in Belgium. Special issue on the occasion of the XIVth Congress of the International Union for Prehistoric and Protohistoric Sciences* 77–89 (SRBAP, Brussel, 2001).

36. van Berg, P. L. & Hauzeur, A. Le Néolithique ancien. *Anthropologica et Præhistorica 112* 63–76 (2001).

37. Kruk, J. *The Neolithic Settlement of Southern Poland*. (Archaeopress, Oxford, 1980).

38. Price, T. D. The introduction of farming in northern Europe. in *Europe's First Farmers* (ed. Price, T. D.) 260–300 (Cambridge University Press, Cambridge, 2000). doi:10.1017/CBO9780511607851.011.

39. Regenye, J. *et al.* Narratives for Lengyel funerary practice. *Bericht der Römisch-Germanischen Kommission Bd. 97 2016(2020)* 5–80 (2020) doi:10.11588/DATA/2EVBVW.

40. Louwe Kooijmans, L. P. Mesolithic/Neolithic transformation in the lower Rhine basin. in *Case Studies in European Prehistory* (ed. Bogucki, P. I.) 95–145 (CRC Press, Boca Raton, 1993).

41. Fokkens, H., Steffens, B. J. W. & van As, S. F. M. *Farmers, Fishers, Fowlers, Hunters. Knowledge Generated by Development-Led Archaeology about the Late Neolithic, the Early Bronze Age and the Start of the Middle Bronze Age (2850 - 1500 Cal BC) in the Netherlands.* vol. 53 (Rijksdienst voor het Cultureel Erfgoed, Amersfoort, 2016).

42. Orschiedt, J., Gehlen, B., Schön, W. & Gröning, F. The Neolithic and Mesolithic Cave site 'Blätterhöhle' in Westphalia (D). *Notae Prehistoricae 32* 73–88 (2012).

43. Bollongino, R. *et al.* 2000 years of parallel societies in Stone Age Central Europe. *Science* **342**, 479–481 (2013).

44. Lipson, M. *et al.* Parallel palaeogenomic transects reveal complex genetic history of early European farmers. *Nature* **551**, 368–372 (2017).

45. *Doorbraken Aan de Rijn. Een Swifterbant-Gehucht, Een Hazendonk-Nederzetting En Erven En Graven Uit de Bronstijd in Medel-De Roeskamp*. (RAAP/Archol/ADC ArcheoProjecten/BAAC, Weesp/Leiden/Amersfoort/'s-Hertogenbosch, 2023).

46. Louwe Kooijmans, L. P. & Jongste, P. F. B. *Schipluiden. A Neolithic Settlement on the Dutch North Sea Coast, c. 3500 Cal BVC*. vol. 37/38 (Faculty of Archaeology, Leiden, 2006).

47. Mol, J., Louwe Kooijmans, L. P. & Hamburg, T. D. 2 Stratigrahy and chronology of the site. in *Schipluiden: A Neolithic Settlement on the Dutch North Sea Coast c. 3500 cal BC* (eds. Louwe Kooijmans, L. P. & Jongste, P. F. B.) 19–38 (Leiden University Press, Leiden, 2006).

48. Raemaekers, D. C. M. & Rooke, M. The Schipluiden pottery. in *Schipluiden: A Neolithic Settlement on the Dutch North Sea Coast c. 3500 cal BC* (eds. Louwe Kooijmans, L. P. & Jongste, P. F. B.) 113–128 (Leiden University Press, Leiden, 2006).

49. ten Anscher, T. J. Leven met de vecht. schokland-p14 en de noordoostpolder in het neolithicum en de bronstijd. (Amsterdam University, Amsterdam, 2012).

50. Raemaekers, D. C. M. *et al.* The submerged pre-drouwen trb settlement site wetsingermaar, C. 3500 CAL. BC (province of Groningen, The Netherlands). *Palaeohistoria* **53**, 1–24 (2012).

51. Louwe Kooijmans, L. P. The Neolithic at the Lower Rhine. Its structure in chronological and geographical respect. *Dissertationes Archaeologicae Gandenses* **16**, 149–173 (1976).

52. Raemaekers, D. C. M. & de Roever, J. P. The Swifterbant pottery tradition (5000-3400 BC). Matters of fact and matters of interest. in *Pots, Farmers and Foragers. Pottery traditions and social interaction in*

*the earliest Neolithic of the Lower Rhine Area* (eds. Vanmontfort, B., Louwe Kooijmans, L. P., Amkreutz, L. W. S. W. & Verhart, L. B. M.) 135–149 (Leiden University Press, Leiden, 2010).

53. Vanmontfort, B. The Group of Spiere as a New Stylistic Entity in the Middle Neolithic Scheldt Basin. *Notae Praehistoricae* **21**, 139–143 (2001).

54. Vanmontfort, B. Can we attribute the middle Neolithic in the Scheldt and middle Meuse basins to the Michelsberg Culture? in *Impacts interculturels au Néolithique Moyen. du terroir au territoire: sociétés et espaces* (ed. Duhamel, P.) 109–116 (Artehis Éditions, Dijon, 2006).

55. Crombé, P., Boudin, M. & Van Strydonck, M. Swifterbant pottery in the Scheldt Basin and the emergence of the earliest indigenous pottery in the sandy lowlands of Belgium. in *Early pottery in the baltic - dating, origin and social context: International workshop at Schleswig from 20th to 21st October 2006* (eds. Hartz, S., Lüth, F. & Terberger, T.) vol. Bericht der Römisch-Germanischen Kommission band 89 465–484 (Philipp von Zabern, Darmstadt, 2011).

56. Crombé, P. & Vanmontfort, B. The neolithisation of the Scheldt basin in western Belgium. in *Going Over: The Mesolithic-Neolithic Transition in North-West Europe* (eds. Whittle, A. W. R. & Cummings, V.) 263–285 (The British Academy, London, 2007).

57. Frébutte, C., Toussaint, M., Masy, P., Pirson, S. & Hubert, F. Campagne archéologique 2001 sur le site du «champ mégalithique de Wéris» à Durbuy (province de Luxembourg). *Notae Praehistoricae 21* 157–173 (2001).

58. Veselka, B. *et al.* Assembling Ancestors: the manipulation of Neolithic and Gallo-Roman skeletal remains from Pommeroeul, Belgium. *Antiquity* 1–16 (2024).

59. Immel, A. *et al.* Genome-wide study of a Neolithic Wartberg grave community reveals distinct HLA variation and hunter-gatherer ancestry. *Commun Biol* **4**, 113 (2021).

60. Salanova, L. *et al.* Du Néolithique récent à l'âge du Bronze dans le centre nord de la France : les étapes de l'évolution chrono-culturelle. in *Le Néolithique du Nord de la France dans son contexte européen :*

*habitat et économie aux 4e et 3e millénaires avant notre ère. Actes du 29e colloque interrégional sur le Néolithique Villeneuve-d'Ascq 2-3 octobre 2009* (eds. Bostyn, F., Martial, E. & Praud, I.) vol. Revue archéologique de Picardie. Numéro spécial 28 77–102 (2011).

61. Modderman, P. J. R. The Neolithic burial vault at Stein. *Analecta Praehistorica Leidensia 1* 3–16 (1964).

62. Verhart, L. B. M. & Amkreutz, L. W. S. W. *Een Nieuwe Blik Op de Grafkelder van Stein*. (2017).

63. Amkreutz, L. W. S. W. Funerary practices on the fringe. The social dimensions of the Neolithic burial chamber of Stein and its European connections. in *The Early Neolithic of northern Europe. New approaches to migration, movement and social connection* (eds. Hofmann, D., Cummings, V., Bjørnevad-Ahlqvist, M. & Iversen, R.) 21–34 (Sidestone Press, Leiden, 2025).

64. *Ypenburg-Locatie 4: Een Nederzetting Met Grafveld Uit Het Midden Neolithicum in Het West-Nederlandse Kustgebied*. (Hazenbeg Archaeologie, Leiden, 2008).

65. Bakker, J. A. *The TRB West Group. Studies in the Chronology and Geography of the Makers of Hunebeds and Tiefstich Pottery*. (University of Amsterdam, Amsterdam, 1979).

66. Deichmüller, J. Die neolithische Moorsiedlung Hüde I am Dümmer, Kreis Grafschaft Diepholz, Vorläufiger Abschlussbericht. *Neue Ausgrabungen und Forschungen in Niedersachsen* **4**, 28–36 (1969).

67. Allentoft, M. E. *et al.* 100 ancient genomes show repeated population turnovers in Neolithic Denmark. *Nature* **625**, 329–337 (2024).

68. Iversen, R. The Pitted Ware Complex in a large scale perspective. *Acta Archaeologica* **81**, 5–41 (2010).

69. Iversen, R., Philippsen, B. & Persson, P. Reconsidering the Pitted Ware chronology: A temporal fixation of the Scandinavian Neolithic hunters, fishers and gatherers. *Praehistorische Zeitschrift* **96**, 44–88 (2021).

70. Coutinho, A. *et al.* The Neolithic Pitted Ware culture foragers were culturally but not genetically influenced by the Battle Axe culture herders. *American journal of physical anthropology* **172**, (2020).

71. Vanhanen, S. *et al.* Maritime Hunter-Gatherers Adopt Cultivation at the Farming Extreme of Northern Europe 5000 Years Ago. *Scientific Reports* **9**, 4756 (2019).

72. Louwe Kooijmans, L. P. *The Rhine/Meuse Delta; Four Studies on Its Prehistoric Occupation and Holocene Geology*. (Instituut voor Prehistorie, Leiden, 1974).

73. Cauwe, N. Les sépultures collectives néolithiques en grotte du Bassin mosan. Bilan documentaire. *Anthropologica et Præhistorica* **115**, 217–224 (2004).

74. Toussaint, M. *et al.* La Grotte Ambre à Matagne-la-Grande (Doische, Namur, Belgique) : étude anthropologique, biogéochimique et archéologique d'un amas d'ossements humains du Néolithique final du bassin mosan wallon. in *Deuxièmes Journées d'actualité de la recherche archéologique en Ardenne-Eifel Actes du colloque tenu à Viroinval 17-19 octobre 2019* (eds. Smolderen, A. & Cattelain, P.) 63–100 (Centre d'Études et de Documentation Archéologiques (Cedarc), Treignes, 2020).

75. Haeck, J. La grotte du Mont Falise à Antheit, vallée de la Méhaigne, province de Liège. B 74: 39-54. *ulletin de la Société royale belge d'Anthropologie et de Préhistoire* **74**, 39–54 (1964).

76. Blanchet, J.-C. *Les Premiers Métallurgistes En Picardie et Dans La Nord de La France. Chalcolitique, Age Du Bronze et Début Du Premier Age Du Fer*. (CTHS, Paris, 1984).

77. Brunet, P. *et al.* La céramique de la fin du 4e et du 3e millénaire dans le Centre-Nord de la France: Bilan documentaire. in *Le troisième millénaire dans le nord de la France et en Belgique. Actes de la journée d'études SRBAP-SPF, 8 mars 2003, Lille* (eds. Vander Linden, M. & Salanova, L.) 155–178 (SRBAP, Brussel, 2004).

78. Lambot, B. Le site chalcolithique du Gord à Compiègne (Oise) note préliminaire. *Cahiers Archéologique de Picardie* **8**, 5–18 (1981).

79. Cottiaux, R. La céramique du site éponyme du ‘Gord’ à Compiègne (Oise). *bspf* **92**, 97–106 (1995).

80. Blanchet, J.-C. & Lambot, B. Quelques aspects du Chalcolithique et du Bronze ancien en Picardie. *Revue Archéologique de Picardie* **3–4**, 79–118 (1985).

81. Martial, E., Praud, I. & Bostyn, F. Recherches récentes sur le Néolithique final dans le nord de la France. *Anthropologica et Præhistorica* **115**, 49–71 (2004).

82. Piningre, J.-F. Un aspect de la fin du Néolithique dans le Nord de la France. Les sites de Seclin, Houplin-Ancoisne et Saint-Saulve (Nord). *pica* **3**, 53–69 (1985).

83. Demeyere, F., Bourgeois, J. & Crombé, P. Plan d’une maison du groupe de Deûle-Escaut à Waardamme (Oostkamp, Flandre occidentale). (2004).

84. Oueslati, T., Leroy, G. & Salvador, P.-G. Fowling on the banks of the Scheldt river in the recent Neolithic (France, 3300-2900 cal BC). *Quaternary International* **626–627**, 52–61 (2022).

85. Sergant, J. *et al.* Een tweede vindplaats van de Deûle-Escaut groep in de Vlaamse zandstreek De site van Hertsberge – Papenvijvers 3 (gem. Oostkamp, West-Vlaanderen, België). *Notae Praehistoricae* **29**, 93–99 (2009).

86. Delcourt-Vlaeminck, M. Les exportations du silex du Grand-Pressigny et du matériau tertiaire dans le nord-ouest de I’Europe au Néolithique final / Chalcolithique. *Anthropologica et Præhistorica* **115**, 139–154 (2004).

87. Mallet, N., Ihuel, E. & Verjux, C. La diffusion des silex du Grand-Pressigny au néolithique / Diffusion of Grand-Pressigny flint during Neolithic. *Supplément à la Revue archéologique du centre de la France* **38**, 131–147 (2012).

88. Ihuel, E., Mallet, N., Pelegrin, J. & Verjux, C. The dagger phenomenon: circulation from the Grand-Pressigny region (France, Indre-et-Loire) in Western Europe. in *The Bell Beaker transition in Europe.*

*Mobility and local evolution during the 3rd Millennium BC* (eds. Prieto Martinez, M. P. & Salanova, L.) 113–126 (Oxbow Books, Oxford, 2015).

89. Fokkens, H. The structure of Late Neolithic and Early Bronze Age settlements and houses in the Netherlands. in *Siedlungsarchäologie des Endneolithicums und der frühen Bronzezeit. 11. Mitteldeutsche Arcäologentag vom 18. bis 20. Oktober 2018 in Halle (Saale)* (eds. Meller, H., Friederich, S., Küssner, M., Staüble, H. & Risch, R.) 915–936 (Landesmuseum für Vorgeschichte, Halle, 2019).

90. Martial, E. & Praud, I. Une nouvelle occupation du Néolithique final dans le Nord, à Baisieux : présentation liminaire. *InterNéo 12* 127–138 (2018).

91. Nobles, .G. R. 3. Features. in *A Mosaic of Habitation at Zeewijk (the Netherlands) Late Neolithic Behavioural Variability in a Dynamic Landscape* (eds. Theunissen, E. M., Brinkkemper, O., Lauwerier, R. C. G. M., Smit, B. I. & Van der Jagt, I. M. M.) 39–54 (Cultural Heritage Agency of the Netherlands, Amersfoort, 2014).

92. van Kampen, J. C. G. & van den Brink, V. B. *Archeologisch Onderzoek Op de Habraken Te Veldhoven. Twee Unieke Nederzettingen Uit Het Laat Neolithicum En de Midden Bronstijd En Een Erf Uit de Volle Middeleeuwen*. (2013).

93. Toussaint, M. Les sépultures mésolithiques du bassin mosan wallon: où en est la recherche en 2010? *Les sépultures mésolithiques du bassin mosan wallon: où en est la recherche en 2010?* 69–89 (2010).

94. de Groote, I. *et al.* Report on the latest excavation campaigns at Grotte de La Faucille, Sclayn (BE) : new radiocarbon dates for a better understanding of burial practice during the Final Neolithic. *(2022) NOTAE PRAEHISTORICAE* 161–177 (2022).

95. Kroon, E. J. *Serial Learners. Interactions between Funnel Beaker West and Corded Ware Communities in the Netherlands during the Third Millennium BCE from the Perspective of Ceramic Technology*. (Sidestone Press, Leiden, 2024).

96. Bourgeois, Q. P. J., Kroon, E. J. & Olerud, L. S. Parallel societies: evidence for the co-existence of Late Funnel Beaker West and Early Corded Ware communities, in. in *The Eve of Destruction? Local groups and large-scale networks during the late fourth and early third millennium BC in central Europe* (eds. Hofmann, D., Mischka, D. & Scharl, S.) (Sidestone Press, Leiden, 2025).

97. Beckerman, S. *Corded Ware Coastal Communities. Using Ceramic Analysis to Reconstruct Third Millennium BC Societies in the Netherlands*. (Sidestone Press, Leiden, 2015).

98. Kroon, E. J., Huisman, D. J., Bourgeois, Q. P. J., Braekmans, D. J. G. & Fokkens, H. The introduction of Corded Ware Culture at a local level: An exploratory study of cultural change during the Late Neolithic of the Dutch West Coast through ceramic technology. *Journal of Archaeological Science: Reports* **26**, 101873 (2019).

99. Furholt, M. Mobility and social change: understanding the european Neolithic period after the archaeogenetic revolution. *J Archaeol Res* **29**, 481–535 (2021).

100. Lanting, J. N. De NO-Nederlandse/NW-Duitse Klokbekergroep: Culturele achtergrond, typologie van het aardewerk, datering, verspreiding en grafritueel. *Palaeohistoria* **49/50 (2007-2008)**, 11–326 (2008).

101. Needham, S. P. Transforming Beaker Culture in North-West Europe; Processes of Fusion and Fission. *Proceedings of the Prehistoric Society* **71**, 171–217 (2005).

102. Fitzpatrick, A. P. The arrival of the Beaker Set in Britain and Ireland: rethinking the Bronze Age and the arrival of Indo-European in Atlantic Europe. in *Celtic from the West 2* (eds. Koch, J. T. & Cunliffe, B. W.) 41–70 (Oxbow books, Oxford, 2013).

103. Lanting, J. N. & van der Waals, J. D. *Glockenbecher Symposium Oberried 1974*. (Fibula-Van Dishoeck, Bussum, 1976).

104. Wentink, K. *Stereotype: The Role of Grave Sets in Corded Ware and Bell Beaker Funerary Practices*. (Sidestone Press, Leiden, 2020).

105. Fokkens, H., Veselka, B., Bourgeois, Q., Olalde, I. & Reich, D. Excavations of Late Neolithic arable, burial mounds and a number of well-preserved skeletons at Oostwoud-Tuithoorn; a re-analysis of old data. *Analecta Praehistorica Leidensia* **47**, 95–150 (2017).

106. Olalde, I. *et al.* The Beaker phenomenon and the genomic transformation of northwest Europe. *Nature* **555**, 190–196 (2018).

107. Lohof, E., Hamburg, T. & Flamman, J. *Steentijd Opgespoord. Archeologisch Onderzoek in Het Tracé van de Hanzelijn-Oude Land*. vol. Archol rapport 138 & ADC rapport 2576 (Archol bv & ADC ArcheoProjecten bv, Amersfoort, 2011).

108. Besse, M. Bell Beaker Common Ware during the third Millennium BC in Europe. in *Similar but different. Bell beakers in Europe* (ed. Czebreszuk, J.) 127–148 (Adam Mickiewicz University, Poznan, 2004).

109. Arnoldussen, S. *A Living Landscape. Bronze Age Settlement Sites in the Dutch River Area (*c. *2000-800 BC)*. (Sidestone Press, Leiden, 2008).

110. Kootker, L. M. & De Coster, M. R. A. L. Chromatographic separation of strontium in archaeological human and faunal enamel for Thermal Ionisation Mass Spectrometry (TIMS) analysis. protocols.io. (2024) doi:https://dx.doi.org/10.17504/protocols.io.bp2l628nkgqe/v1.

111. Haak, W. *et al.* Massive migration from the steppe was a source for Indo-European languages in Europe. *Nature* **522**, 207–211 (2015).

112. Patterson, N. *et al.* Large-scale migration into Britain during the Middle to Late Bronze Age. *Nature* **601**, 588–594 (2021).

113. Ringbauer, H. *et al.* Accurate detection of identity-by-descent segments in human ancient DNA. *Nature Genetics* **56**, 143–151 (2024).

114. Rubinacci, S., Ribeiro, D. M., Hofmeister, R. J. & Delaneau, O. Efficient phasing and imputation of low-coverage sequencing data using large reference panels. *Nature Genetics 53* 120–126 (2021).

115. The 1000 Genomes Project Consortium. An integrated map of genetic variation from 1,092 human genomes. *Nature 491* 56–65 (2012).
